# Structural basis of mitochondrial protein import by the TIM complex

**DOI:** 10.1101/2021.10.10.463828

**Authors:** Sue Im Sim, Yuanyuan Chen, Eunyong Park

## Abstract

Mitochondria import nearly all their ∼1,000–2,000 constituent proteins from the cytosol across their double membrane envelope. Genetic and biochemical studies have shown that the conserved protein translocase, termed the TIM complex (also known as TIM23 complex), mediates import of presequence-containing proteins into the mitochondrial matrix and inner membrane. Among ∼10 different subunits of the complex, the essential multi-pass membrane protein Tim23, together with the evolutionarily related protein Tim17, has long been postulated to form a protein-conducting channel. However, the mechanism of TIM-mediated protein import remains uncertain due to a lack of structural information on the complex. Here, we have determined the cryo-EM structure of the core TIM complex (Tim17–Tim23–Tim44) from *Saccharomyces cerevisiae*. We show that, contrary to the prevailing model, Tim23 and Tim17 do not form a water-filled channel, but instead have separate, lipid-exposed concave cavities that face in opposite directions. Remarkably, our data suggest that the cavity of Tim17 itself forms the protein translocation path whereas Tim23 plays a structural role. We also show how the Tim17–Tim23 heterodimer associates with the scaffold protein Tim44 and J-domain proteins to mediate Hsp70-driven polypeptide transport into the matrix. Our work provides the structural foundation to understand the mechanism of TIM-mediated protein import and sorting, a central pathway in mitochondrial biogenesis.

## INTRODUCTION

Mitochondria are endosymbiotically-derived double membrane-bound organelles that perform numerous vital cellular functions in eukaryotes, such as oxidative phosphorylation. Nearly 99% of ∼1,000–2,000 mitochondrial proteins are encoded by nuclear genes and imported into mitochondria shortly after synthesis on cytosolic ribosomes. The import processes are largely mediated by two conserved molecular machines—the translocase of the outer membrane (TOM) and translocase of the inner membrane (TIM) complexes (for reviews, see ref. 1-5). The TOM complex mediates the initial translocation of precursor proteins (preproteins) across the outer membrane (OM) (6-10).

The TIM complex (also frequently termed the presequence translocase or TIM23 complex) further translocates preproteins into the inner membrane (IM) and across the IM into the mitochondrial matrix. Its client proteins are typically targeted to the TIM complex by a short N-terminal cleavable localization signal (also known as a presequence), which adopts a positively-charged amphipathic α-helix (11, 12). If clients contain a membrane-spanning hydrophobic sorting signal, the TIM complex also integrates such proteins into the IM (13, 14). It has been generally thought that the TIM complex forms a water-filled channel that transports polypeptides across the IM (1, 2, 4, 5, 15). It has been further postulated that the channel can open laterally for hydrophobic sorting signals to enter the lipid phase (16-18).

The TIM complex consists of ∼10 different proteins, with the multi-pass integral membrane proteins Tim17 and Tim23 forming its indispensable core (19-24). Tim17 and Tim23 are evolutionarily related to each other. Frequently, Tim23 has been described as the primary channel-forming subunit, whereas Tim17 has been regarded as a regulatory subunit (25-30). The TIM complex exists in two overlapping but distinct compositional states: a motor-associated (TIM^MOTOR^) form and a motor-free (TIM^SORT^) form (16, 31, 32) (Fig. S1 A and B). TIM^MOTOR^ is required for translocation of soluble matrix-targeted proteins, whereas TIM^SORT^ is thought to be specialized for integration of membrane proteins. Both forms share, in addition to Tim17 and Tim23, another essential subunit, Tim50. Tim50, a single-spanning membrane protein containing a soluble domain in the intermembrane space (IMS), may guide preproteins to the Tim17–Tim23 complex (33, 34).

TIM^MOTOR^ includes four additional essential proteins, Tim44, Pam16, Pam18, and mitochondrial heat-shock protein 70 (mtHsp70), collectively known as the presequence translocase-associated motor or PAM (Fig. S1A). Tim44 serves as a scaffold to recruit the mtHsp70 ATPase to the matrix side of the complex (35-38), which provides the driving force for directional transport via its interactions with the translocating polypeptide (39, 40). Pam18 and Pam16 are Hsp40-related J-domain and J-domain-like proteins that form a heterodimer that act in conjunction with the mtHsp70 (41-44). In the TIM^SORT^ form, the core TIM complex associates with two nonessential membrane proteins Tim21 and Mgr2 instead of the PAM (16, 17, 31, 32) (Fig. S1B).

Here, we present the near-atomic-resolution structure of the core TIM^MOTOR^ complex (Tim17– Tim23–Tim44) from *S. cerevisiae* determined by cryo-electron microscopy (cryo-EM).

Combining insights from the structure with mutational analysis, we provide evidence for several surprising conclusions. First, Tim23, Tim17, and Tim44 form a 1:1:1 heterotrimer as the core TIM^MOTOR^ complex. Second, Tim23 and Tim17 do not form an aqueous pore as hitherto assumed. Third, protein translocation is mediated by a lipid-exposed concave cavity formed by Tim17 itself, not by Tim23. These findings and other observations presented here support a paradigm-shifting model for TIM-mediated mitochondrial protein import and sorting.

## RESULTS

### Purification and cryo-EM analysis of the yeast TIM complex

We could purify an endogenous TIM complex, likely a mixture of TIM^MOTOR^ and TIM^SORT^, from *S. cerevisiae* by attaching an affinity tag to Tim17. The isolated complex contained mainly Tim17, Tim23, Tim44, Tim50, Tim21, and mtHsp70 (Fig. S1C), similar to previous reports (31, 34). However, due to low cellular abundance, it was difficult to obtain sufficient quantities for biochemical and structural analysis. We thus co-overexpressed in yeast all known subunits of the TIM complex (except for mtHsp70)—six essential proteins, Tim17, Tim23, Tim44, Tim50, Pam16, and Pam18, and three nonessential proteins, Pam17, Tim21, and Mgr2 (17, 31, 45). The complex purified using this approach exhibited properties similar to the endogenous complex (Fig. S1D). In addition, Pam16 and Pam18 (referred to as the Pam16/18 complex), components known to be weakly associated with the Tim17–Tim23 core (41, 45), appeared to be present in higher stoichiometry, as judged by their greater visibility in SDS gels. A stable, well-behaved complex could also be formed and purified by overexpressing only Tim17, Tim23, and Tim44 (Fig. S1E), consistent with the idea that Tim50 and Pam16/18 are peripherally tethered and do not contribute to the core membrane structure (33, 44, 46-48). When the affinity tag was attached to Tim21 instead of Tim17, the PAM did not co-purify (Fig. S1F), in agreement with the composition expected for the TIM^SORT^ form (Fig. S1B). Thus, the purification results recapitulate known biochemical characteristics of the TIM complex and indicate that the complexes are properly assembled.

Our initial cryo-EM analysis of the purified TIM complex yielded disc-like reconstructions, but only to low (∼8-Å) resolution (Figs. S1D and S2). The poor resolution seemed to originate from the small overall molecular size of the complex and its lack of large protruding features. In one class (Class 2), however, we observed a small central protruding feature (Fig. S2E). We speculated that this protuberance was likely the 20-kDa conserved C-terminal domain (CTD) of Tim44, as it has been shown to interact extensively with both Tim17 and Tim23 (38).

To achieve high-resolution reconstructions, we developed a monoclonal antibody against purified Tim44-CTD. We decorated the TIM^MOTOR^ complex comprised of Tim17, Tim23, Tim44, and Pam16/18 with the antigen-binding fragment (Fab) as a fiducial marker (Table 1 and Fig. 1; and Fig. S3). Inclusion of the Fab dramatically improved the map refinement, allowing us to determine the structure of the complex at 2.7 Å resolution. Tim44-CTD and the transmembrane domains (TMDs) of Tim17 and Tim23 were well resolved, enabling building of an accurate atomic model (Fig. S4A). Density features for Pam16/18 and endogenous mtHsp70 were invisible in this map, likely due to their low occupancy and/or high conformational flexibility (see below).

**Table 1.**
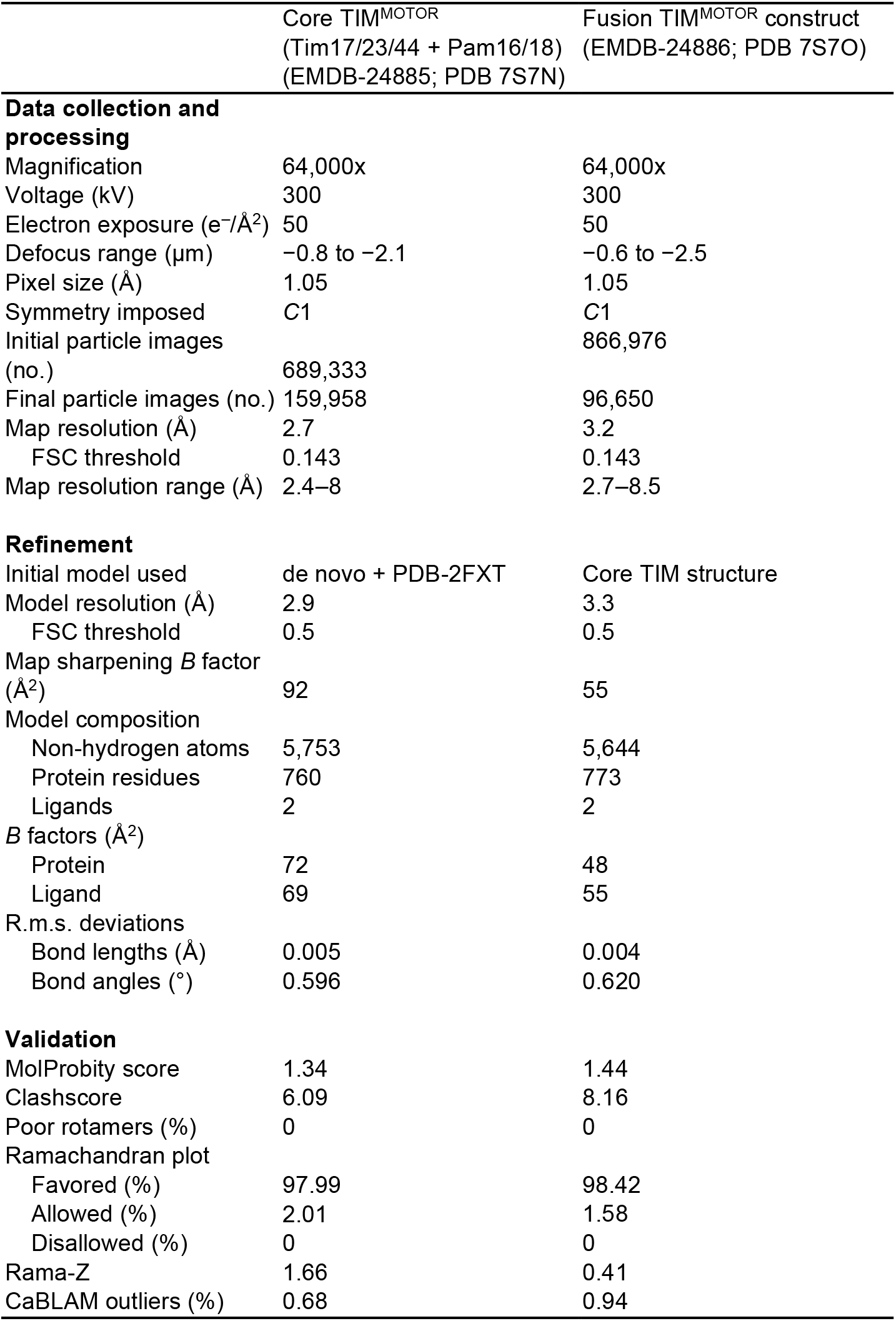
Cryo-EM data collection, refinement and validation statistics.

|  | Core TIM<sup>MOTOR</sup><br>(Tim17/23/44 + Pam16/18)<br>(EMDB-24885; PDB 7S7N) | Fusion TIM<sup>MOTOR</sup> construct<br>(EMDB-24886; PDB 7S7O) |
| --- | --- | --- |
| **Data collection and processing** |  |  |
| Magnification | 64,000x | 64,000x |
| Voltage (kV) | 300 | 300 |
| Electron exposure (e<sup>-</sup>/Å<sup>2</sup>) | 50 | 50 |
| Defocus range (μm) | -0.8 to -2.1 | -0.6 to -2.5 |
| Pixel size (Å) | 1.05 | 1.05 |
| Symmetry imposed | C1 | C1 |
| Initial particle images (no.) | 689,333 | 866,976 |
| Final particle images (no.) | 159,958 | 96,650 |
| Map resolution (Å) | 2.7 | 3.2 |
| FSC threshold | 0.143 | 0.143 |
| Map resolution range (Å) | 2.4–8 | 2.7–8.5 |
| **Refinement** |  |  |
| Initial model used | de novo + PDB-2FXT | Core TIM structure |
| Model resolution (Å) | 2.9 | 3.3 |
| FSC threshold | 0.5 | 0.5 |
| Map sharpening *B* factor (Å<sup>2</sup>) | 92 | 55 |
| Model composition |  |  |
| Non-hydrogen atoms | 5,753 | 5,644 |
| Protein residues | 760 | 773 |
| Ligands | 2 | 2 |
| *B* factors (Å<sup>2</sup>) |  |  |
| Protein | 72 | 48 |
| Ligand | 69 | 55 |
| R.m.s. deviations |  |  |
| Bond lengths (Å) | 0.005 | 0.004 |
| Bond angles (°) | 0.596 | 0.620 |
| **Validation** |  |  |
| MolProbity score | 1.34 | 1.44 |
| Clashscore | 6.09 | 8.16 |
| Poor rotamers (%) | 0 | 0 |
| Ramachandran plot |  |  |
| Favored (%) | 97.99 | 98.42 |
| Allowed (%) | 2.01 | 1.58 |
| Disallowed (%) | 0 | 0 |
| Rama-Z | 1.66 | 0.41 |
| CaBLAM outliers (%) | 0.68 | 0.94 |

**Figure 1.**
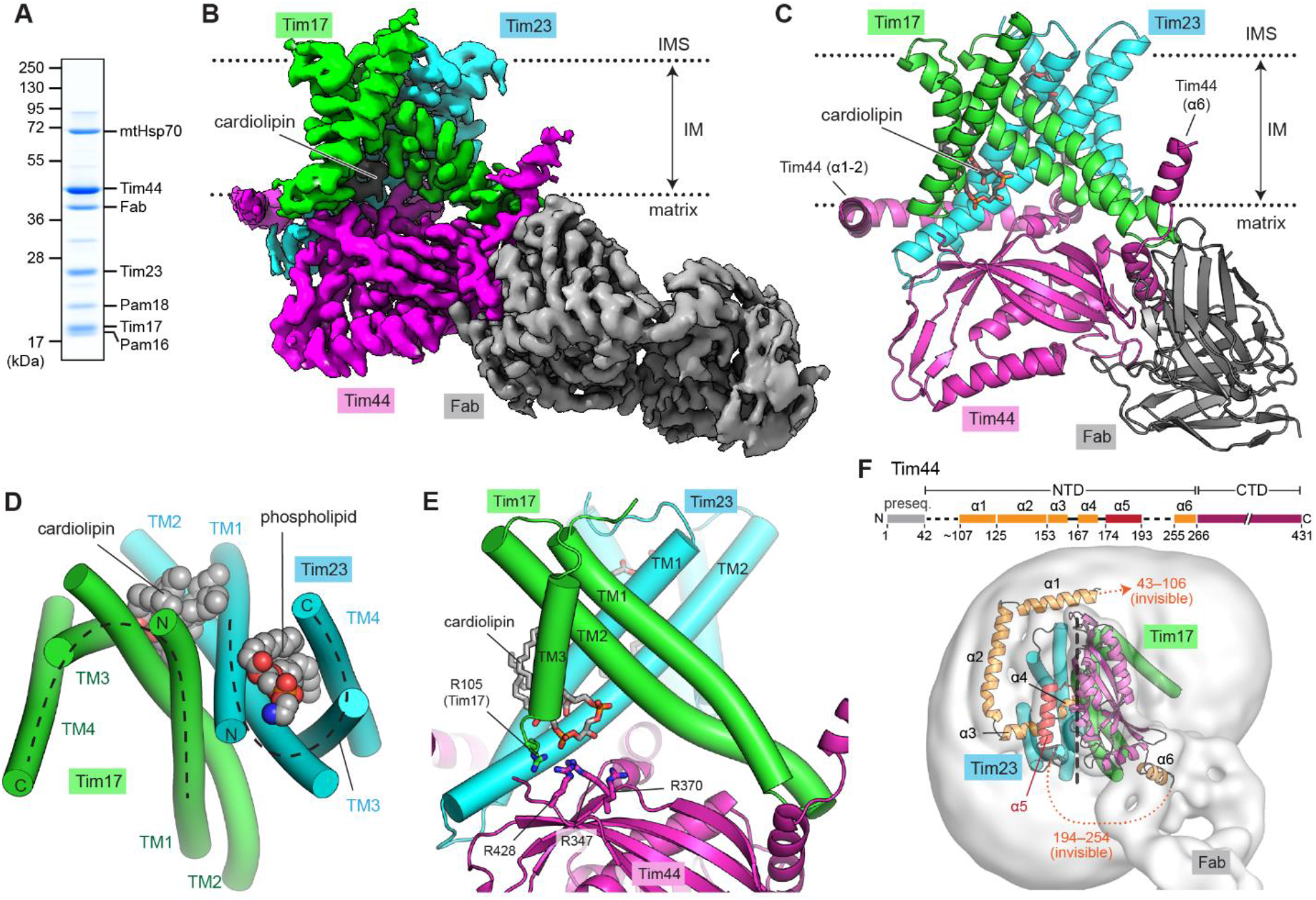
Cryo-EM structure of the TIM complex from *S. cerevisiae*. (**A**) SDS gel image of the affinity-purified, Fab-bound TIM^MOTOR^ complex (Tim17/23/44 + Pam16/18). For full image, see Fig. S3A. (**B**) 2.7-Å-resolution cryo-EM reconstruction of Fab-bound core TIM^MOTOR^ (sample shown in A). The approximate boundaries of the IM are indicated by dotted lines. (**C**) As in B, but showing the atomic model. A model for the constant domains (CH1 and CL) of the Fab was not built. (**D**) Arrangement of the Tim17 and Tim23 TMDs in a view from the IMS. TMs are represented as cylinders (arranged in the N to C order), and lipids are shown as spheres. The phospholipid bound to the Tim23 cavity is likely a phosphatidylethanolamine based on the cryo-EM density (see Fig. S5 A–C). The topology of the cavities is highlighted with dashed lines. (**E**) As in D, but a cutaway view showing the interface between Tim17 and Tim23. Cardiolipin and the neighboring arginine side chains are shown in sticks. (**F**) Organization of Tim44 domains. Upper panel, linear diagram; lower panel, view from the matrix). The detergent micelle is shown with a lowpass-filtered cryo-EM map (gray semitransparent surface). Tim44 is shown as a ribbon diagram (orange, amphipathic helices; red, α5; purple, CTD. Dashed line, plane of the Tim17-Tim23 interface.

### Structure of the Fab-bound Tim17–Tim23–Tim44 complex

Our structure shows that Tim17, Tim23, and Tim44 form a 1:1:1 heterotrimer (Fig. 1 B and C). In the membrane, Tim17 and Tim23 are arranged with pseudo-two-fold rotational symmetry along the membrane normal. Transmembrane segments (TMs) 1–4 of Tim17 are arranged sequentially, but in a tilted fashion, to generate a cavity that is open toward the lipid phase. Looking down from the IMS, the topology resembles a U shape (Fig. 1D). Tim23 also adopts a similar structure but with a substantially narrower cavity, which is occupied by a well-ordered phospholipid molecule (Fig. 1D; and Fig. S5 A–C). Tim17 and Tim23 associate in a “⋂⋃’’ arrangement through their TMs 1–2, which are arranged as crossing diagonals looking from the side (Fig. 1 D and E). TMs 1 and 2 of Tim17 and Tim23 are closely packed against each other with several invariant glycine residues at the interfaces (Fig. S6 A–C). This arrangement explains why the complex was found to be unstable when these glycine residues (e.g., G62L and G66L on Tim17; and G102L and G153L on Tim23) were mutated (49, 50). Because of the pseudo-two-fold symmetry, the cavities of Tim17 and Tim23 face away from each other. The properties of these cavities are discussed in further detail below.

TMs 1–2 of Tim17 and Tim23 create a shallow depression on the matrix side, where Tim44-CTD docks (Fig. 1E). The Tim44-CTD structure could be superimposed (Cα root-mean-square deviation [RMSD] of 0.6 Å) with the previous crystal structure of isolated yeast Tim44-CTD (51) (Fig. S4B). Our structure now reveals how Tim44 is oriented with respect to the Tim17–Tim23 heterodimer. Tim44-CTD has an overall flat structure that associates with Tim17–Tim23 and is vertically aligned along the plane of the Tim17-Tim23 interface, such that one side is oriented towards Tim17 and the other side towards Tim23 (Fig. 1F). The arrangement of Tim44-CTD with respect to Tim17–Tim23 is in excellent agreement with previous data from in-vivo photo-crosslinking experiments (Fig. S6 D and E) (38, 49, 52), strongly supporting that our structure is physiological.

Tim44 is a peripheral membrane protein that is associated with the matrix leaflet of the IM. Our structure reveals that a large part of Tim44-NTD forms membrane-interacting amphipathic helices, which encircle the Tim17–Tim23 heterodimer around the Tim23 side (Fig. 1F; α1–α4 and α6). Although it remains to be elucidated whether the NTD adopts a similarly arced structure in the native membrane, the observed arrangement seems to be enabled in part by α5 segment (residues 174–193) attached to the Tim23-facing side of Tim44-CTD. Mutations in α5 have been shown to cause severe defects in the protein import activity and destabilize the interaction between Tim44 and Tim17–Tim23 (37), suggesting that α5 has a substantial role in the proper arrangement of the Tim44 domains. The ∼60-amino-acid long N-terminal segment preceding the α1 helix is not visible in our structure due to flexibility but appears to be directed towards the Tim17 side (Fig. 1F). This segment is important for an interaction with Pam16 (47, 53). Consistent with this, Pam18, the partner of Pam16, has also been shown to interact with Tim17 (31, 47) (also see Fig. 5 C–E).

Lastly, the structure shows how the TIM complex interacts with the negatively-charged mitochondrial-specific lipid cardiolipin (54-56) (Fig. 1 D and E). A well-resolved density for cardiolipin could be seen in a crevice formed between TMs 2 and 3 of Tim17 and TM2 of Tim23 (Fig. S5D). Multiple arginine side chains from Tim17 (Arg105) and Tim44-CTD (Arg347, Arg370, and Arg428) are positioned near the two phosphate groups of the cardiolipin molecule. In particular, Tim17-Arg105 and Tim44-Arg347 appear to directly interact with one of the cardiolipin phosphate groups (Fig. S5D). Thus, the structure explains previous results that suggested the importance of cardiolipin in maintaining the integrity of the TIM complex (55, 56). Interestingly, the Tim17-R105A mutant has been shown to destabilize the interaction between Tim44 and Tim17–Tim23 (49). It is possible that this defect was caused by decreased affinity of the Tim17 mutant towards cardiolipin.

### Properties of the Tim17 and Tim23 cavities

Contrary to the widely accepted notion, our structure shows that the TIM complex does not form a water-filled protein-conducting channel. Instead, the lipid-exposed concave surface of Tim17 or Tim23 likely constitutes the translocation path. Moreover, the laterally open cavity itself would be compatible with the IM protein integration (sorting) activity of the TIM complex. This idea is consistent with recent cryo-EM structures of another mitochondrial complex called carrier translocase (also termed TIM22 complex) (57, 58), a dedicated insertase that integrates certain multi-pass membrane proteins, such as carrier proteins, into the IM (59). Although the carrier translocase is functionally distinct from the presequence translocase TIM complex, its central subunit (Tim22) belongs to the same protein family as Tim17 and Tim23 and forms a laterally open cavity. Indeed, our structure allowed us to compare all three proteins, and we found that Tim17, Tim23, and Tim22 all share a similar fold (Fig. 2 A and B).

**Figure 2.**
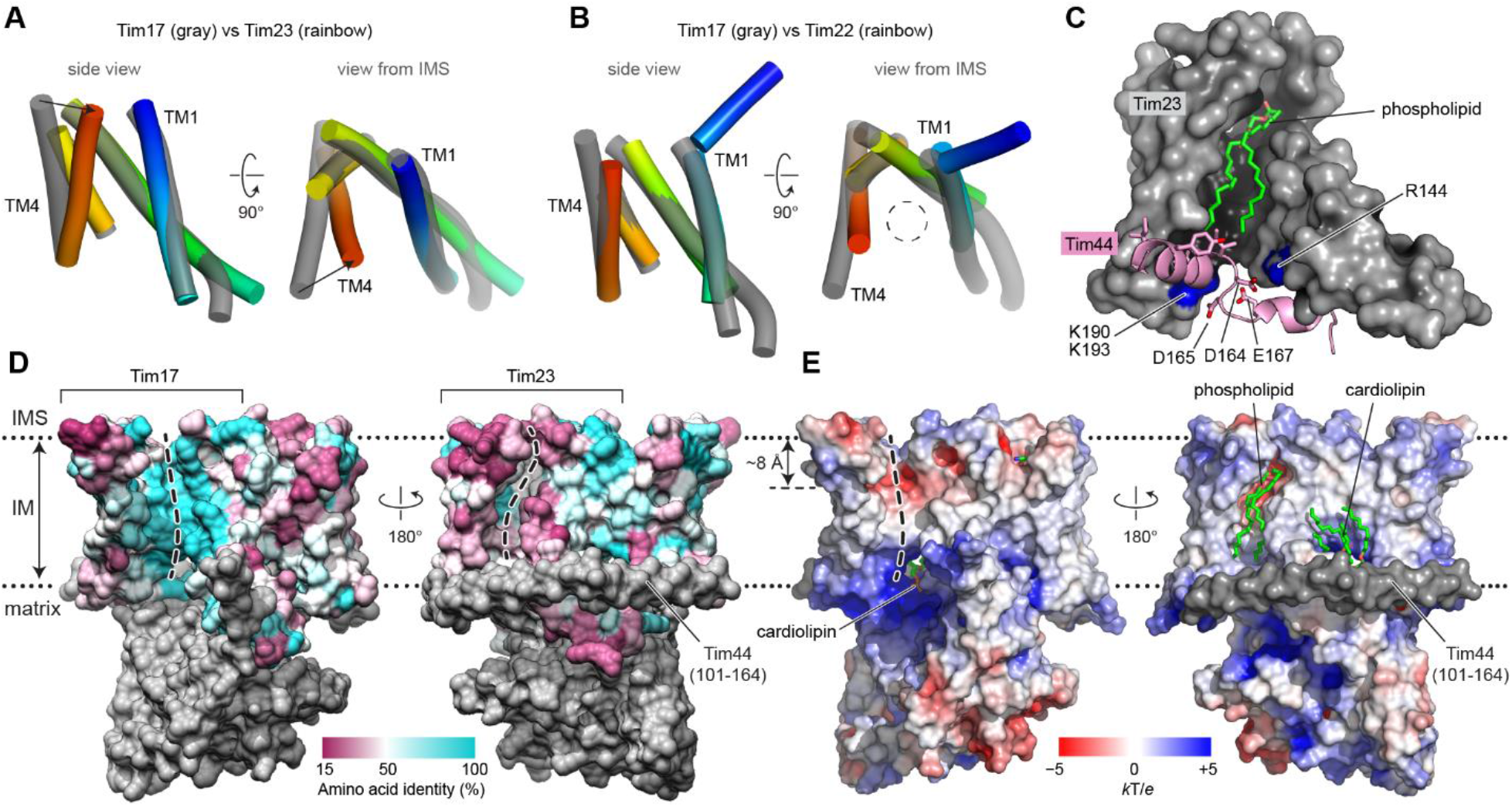
Lipid-exposed concave cavities of Tim17 and Tim23. (**A**) Comparison between the TMDs of Tim17 and Tim23. TM1 to TM4 of Tim23 are shown in rainbow color (blue to red). Arrows highlight a difference between the TM4 positions. (**B**) As in A, but comparing between Tim17 and Tim22 (PDB ID: 6LO8). Note that Tim22 has an extra IMS helix to its N-terminus. (**C**) Side view showing the cavity of Tim23 (gray surface). Amino acid residues 153–174 of Tim44 are shown in pink. Note that residues D164, D165, and E167 interact with basic amino acids (R144, K190, and K193; blue) of Tim23. (**D**) Mapping of amino acid sequence conservation of Tim17 and Tim23 onto the structure. Left, view into the Tim17 cavity; right, view into the Tim23 cavity. Dashed line, path along the cavity. 94 and 102 representative sequences were used for Tim17 and Tim23, respectively. (**E**) Surface electrostatics of the TIM complex. The views are equivalent to those in D. α1–3 (residues ∼107– 164) of Tim44 (modelled as polyalanine) were not included in the calculation.

Both Tim17 and the homologous Tim22 form a similarly sized cavity in the membrane, which is large enough to accommodate a polypeptide chain (Fig. 2B). On the other hand, due to the inwardly tilted position of Tim23-TM4, the cavity of Tim23 is substantially narrower, seemingly too restricted to accommodate even an extended polypeptide (Fig. 2A). Furthermore, the vertical axis of Tim23 is largely obstructed on the matrix side by a segment of Tim44 (positions 153–171; referred to as α3–α4) (Fig. 2C). A partial truncation (Δ150-164) or replacement to alanine (154–157 to Ala_4_) of the α3–α4 segment has been shown to cause growth defects and impair association of Tim44 with the Tim17–Tim23 dimer (37). This suggests that binding of α3– α4 to Tim23 is important for the function and integrity of the TIM^MOTOR^ complex. Taken together, these observations indicate that the cavity of Tim17, rather than that of Tim23, is more likely to be used as the path for protein translocation.

To further evaluate this idea, we mapped the sequence conservation of Tim17 and Tim23 across species onto our structure. We reasoned that among the surface-exposed positions, those exhibiting high sequence identity would more likely mediate protein translocation. Indeed, a sequence comparison indicates that many amino acids lining the cavity of Tim17 are highly conserved whereas those in Tim23 are markedly variable (Fig. 2D). This sequence conservation supports the idea that the cavity of Tim17 constitutes the surface that mediates protein translocation.

### Mutational studies on the Tim17 and Tim23 cavities

Among those highly conserved residues in Tim17, several acidic amino acids (Asp17, Asp76, and Glu126) are positioned on the concave surface in the IMS leaflet, forming a negatively charged patch that extends ∼8 Å deep into the membrane from the IMS (Fig. 2E). On the matrix side, the phosphate groups of the cardiolipin are positioned near the vertical path of the Tim17 cavity. In the middle, the apolar region of the cavity seems to span only ∼10–15 Å of the membrane (Fig. 2E). These features may cause local membrane thinning in the Tim17 cavity, reducing the energy barrier for polypeptide translocation.

To examine the functional importance of the negatively charged surface in Tim17, we mutated the acidic residues and tested cell viability on the basis of the fact that TIM-mediated protein translocation is essential for yeast cell growth (Fig. 3 A–B; and Figs. S7 and S8). Although mutating Asp17, Asp76, and Glu126 individually to a neutral Asn/Gln residue did not affect the cell growth, double or triple mutations severely impaired viability. In addition, single charge-reversal point mutations (D17R and D76R) caused a modest to severe growth defect. Defects by Tim17 mutations were not due to reduced protein expression as stronger expression by an inducible promoter did not alleviate the growth impairment (Fig. S8 C–F). These data suggest that overall negative charges, not individual side chains, are important for function.

**Figure 3.**
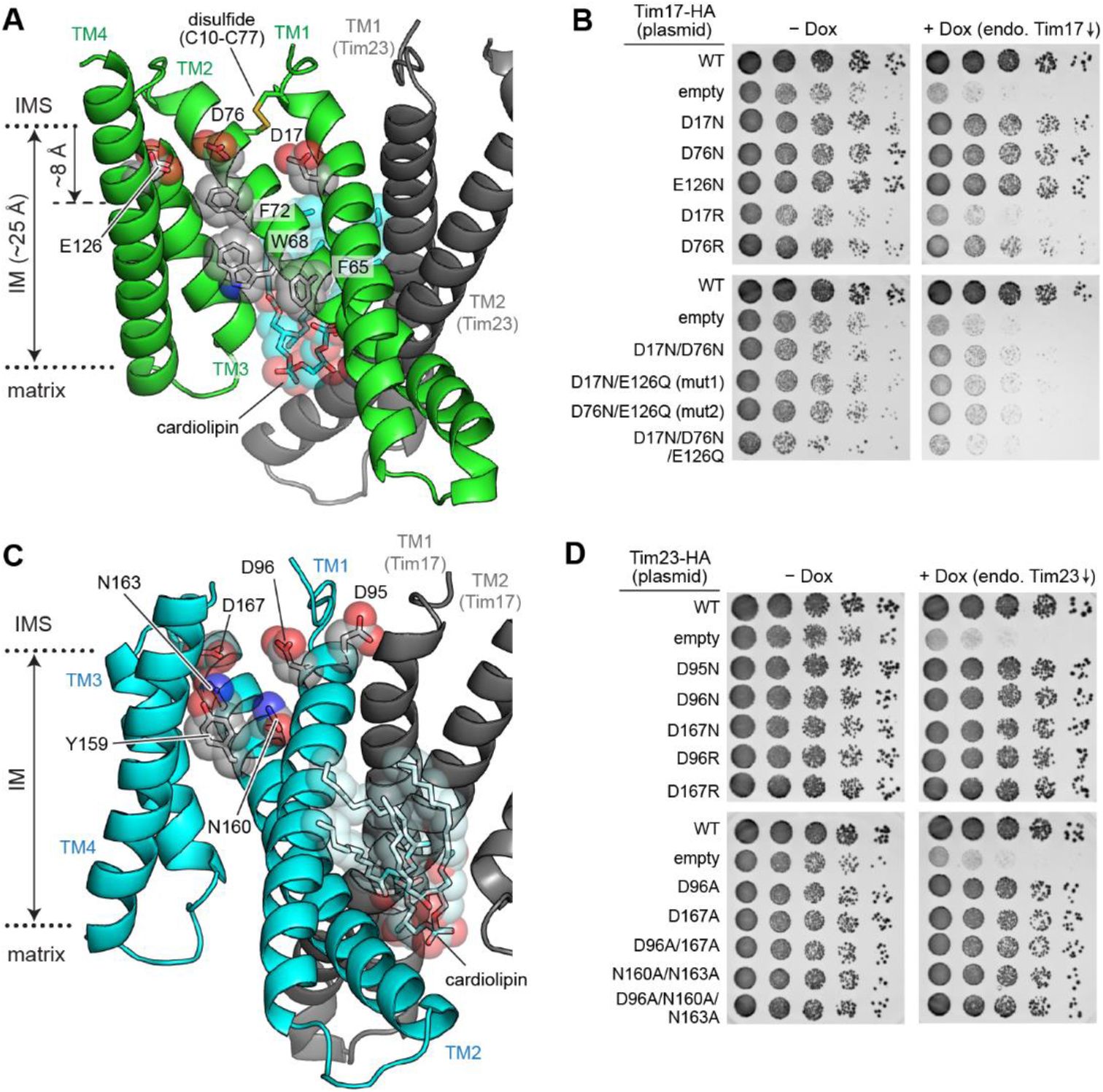
Mutational analysis of the Tim17 and Tim23 cavities. (**A**) Conserved acidic and aromatic amino acids lining the Tim17 cavity. The view is similar to the left panels of Fig. 2 D and E. For clarity, only Tim17 and part of Tim23 are shown. Note that Tim17 contains a disulfide bond between Cys10 and Cys77, as shown previously (79). (**B**) Indicated acidic amino acids lining the Tim17 cavity were mutated, and their functionality was tested by a cell growth assay. The addition of doxycycline (Dox) represses expression of chromosomal WT Tim17. (**C**) As in A, but view into the Tim23 cavity. The view is similar to the right panels of Fig. 2 D and E. (**D**) As in B, but testing mutations of acidic and polar amino acids lining the Tim23 cavity.

Next, we chose two functionally defective Tim17 mutants (D17N/E126Q and D76N/E126Q; referred to as mut1 and mut2, respectively) and examined their defects in the assembly of the TIM complex and translocation of client proteins. Co-immunoprecipitation experiments showed that these Tim17 mutants interact with other Tim and Pam subunits equivalent to WT Tim17 (Fig. 4A). To probe translocation defects in these Tim17 mutants, we generated a stalled translocation intermediate in mitochondria expressing both WT and mutant Tim17 with a model preprotein fused to a dihydrofolate reductase (Cyb2Δ-DHFR) (60) (Fig. 4 B and C; and Fig. S9). When Cyb2Δ-DHFR was pulled down after forming the intermediate under the energized condition (-valinomycin), WT Tim17 could be markedly enriched (Fig. 4C, lanes 2 versus 3). By contrast, mutant Tim17 did not copurify beyond the extent seen without translocation (WT/+valinomycin); only the co-expressed WT Tim17 copies were enriched. Taken together, these data show that mutant Tim17 proteins are properly assembled into the TIM^MOTOR^ complex but cannot engage with preproteins.

**Figure 4.**
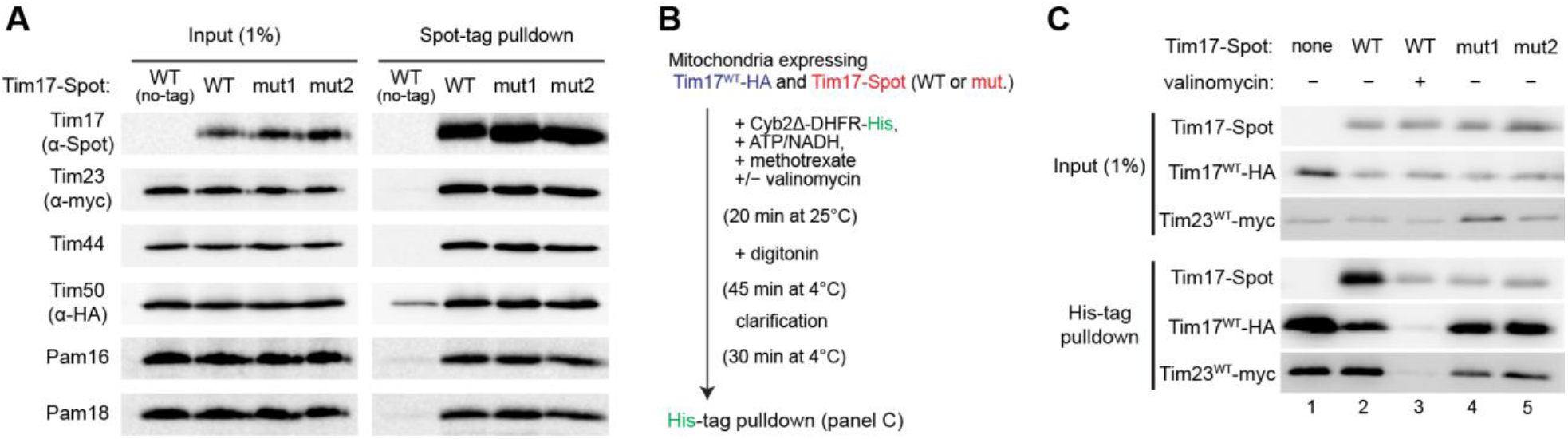
Mutations in the Tim17 cavity cause defects in substrate engagement. (**A**) Co-immunoprecipitation of the subunits of the TIM^MOTOR^ complex with Tim17 mutants. Mitochondria expressing indicated Tim17 variants (mut1= D17N/E126Q; mut2=D76N/E126Q) were solubilized with digitonin, and Tim17 was pulled down with anti-Spot-tag nanobody beads. The samples were analyzed by western blotting with indicated antibodies. (**B**) Schematic for pulldown experiments of stalled translocation intermediates. (**C**) The Cyb2Δ-DHFR-His substrate was pulled down with nickel resin after forming a stalled translocation intermediate, and samples were analyzed by western blotting. Valinomycin dissipates ΔΨ and precludes mitochondrial import.

It has been noted that in the carrier translocase complex, Tim22 exhibits a similar negatively charged patch in its cavity (57, 58). Like Tim17, mutations of those acidic residues in Tim22 impair the function of the carrier translocase (58). We note that Tim23 also has a negatively charged, polar patch in its cavity, mainly from Asp96, Asn160, Asn163, and Asp167 (Figs. 2E and 3C). Unlike Tim17, however, simultaneous mutations of these amino acids to Asn or Ala did not detectably compromise growth (Fig. 3D; and Fig. S7B), providing additional evidence that Tim23 itself is not the route for protein translocation.

In addition to acidic amino acids, the surfaces of the Tim17 and Tim23 cavities also contain conserved aromatic amino acids (Phe65, Trp68 and Phe72 in Tim17; and Tyr159 in Tim23) (Fig. 3 A and C). Mutations of these aromatic amino acids to aliphatic valine did not cause growth defects, suggesting that hydrophobicity may be more important for function than residues identity per se (Figs. S7 C–F, and S8C). Indeed, we note that the F65N mutation on Tim17 greatly reduced cell viability, suggesting that hydrophobicity at this position is important. Taken together, our data strongly suggest that the cavity of Tim17 likely mediates polypeptide translocation, and its acidic patch is critical for this function.

### Possible roles of Tim44 in protein translocation

Our proposal that Tim17 forms the protein translocation path is further supported by the orientation of Tim44-CTD. When a preprotein emerges into the matrix from the membrane, it is expected to be in a largely unfolded state, which would be prone to aggregation. We noticed the Tim17-facing side of the Tim44-CTD presents many solvent-exposed hydrophobic side chains in two clusters. Cluster 1, which is formed by aliphatic amino acids, lines the surface of a twisted β-sheet and is positioned directly below the Tim17 cavity (Fig. 5 A and B). In crystal structures of yeast and human Tim44-CTDs (51, 61), this site is artificially occupied by the membrane-interacting amphipathic helix α6 of the NTD (Fig. S10 A and B). Thus, Cluster 1 may interact with the translocating polypeptide through a hydrophobic interaction. Cluster 2, which is adjacent to Cluster 1 and forms a depressed pocket, may also interact with the translocating polypeptide (Fig. 5 A and B). In the human Tim44-CTD crystal structure, this pocket was occupied by pentaethylene glycol molecules (Fig. S10 C and D) (61). Collectively, these observations suggest that Tim44-CTD tends to associate with other molecules through these hydrophobic surfaces. Thus, Tim44 may function as a molecular chaperone to stabilize the translocating polypeptide against misfolding or aggregation, in addition to its role in recruiting Pam16/18 and mtHsp70.

**Figure 5.**
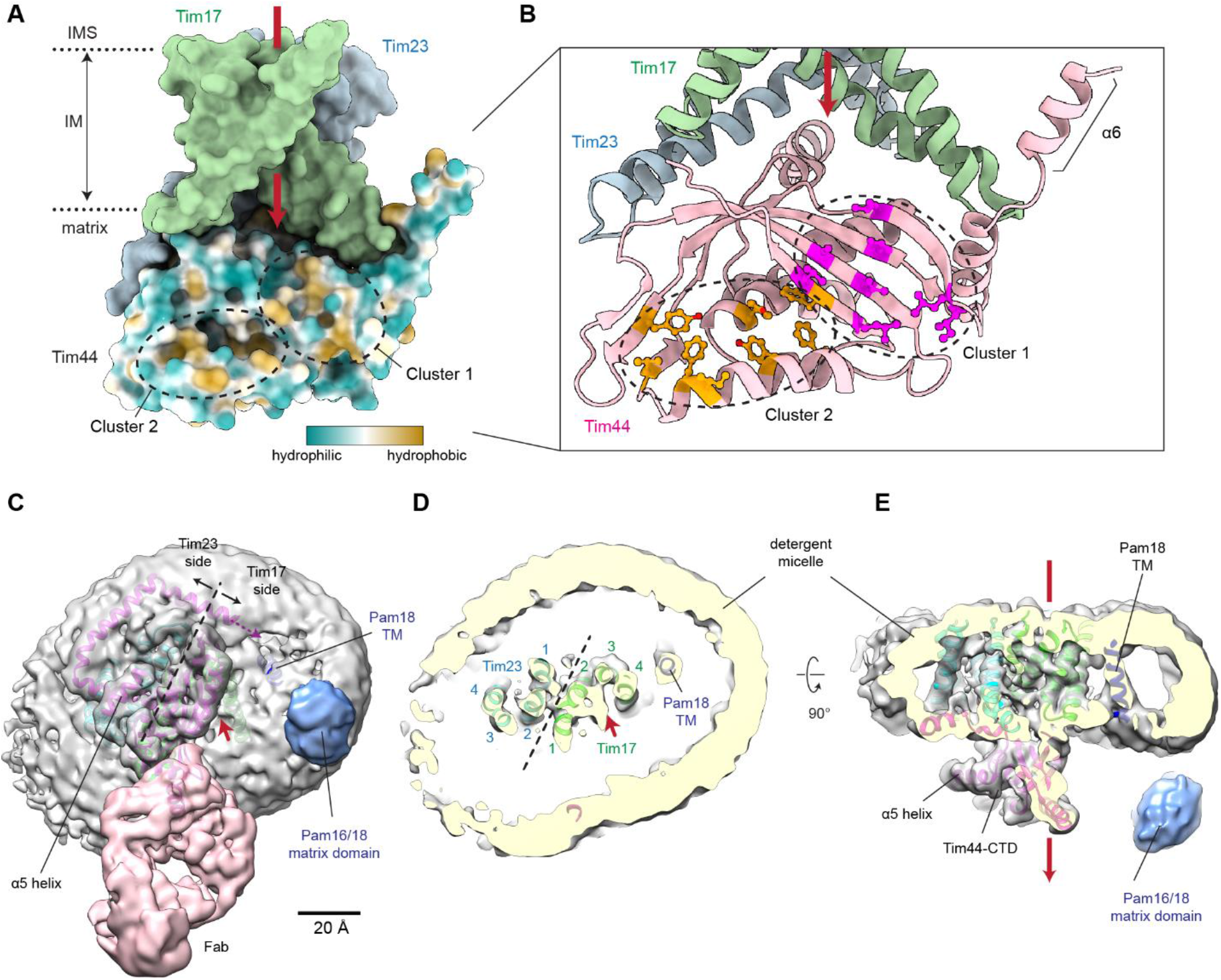
Arrangement of Tim44 and Pam16/18. (**A**) Clusters of solvent-exposed hydrophobic amino acids in Tim44 are shown in a surface representation. The view direction is similar to Fig. 1 B and C. Hydrophobicity is indicated by a color map. A putative path for preprotein translocation is indicated by a red arrow. (**B**) As in A, but a zoomed-in view in a ribbon representation. Solvent-exposed hydrophobic amino acids are shown in purple (Cluster 1) and orange (Cluster 2) with side chains represented as balls and sticks. Note that in the crystal structures of isolated Tim44-CTD, the amphipathic segment α6 folds onto the Cluster 1 (see Fig. S10). (**C**) Cryo-EM structure of the fusion TIM^MOTOR^ construct. To show Pam16/18 features better, the map (surface representation; at a contour level of 5σ) was lowpass-filtered at 5 Å. The atomic model is shown in a ribbon representation (purple, Tim44; green, Tim17; cyan, Tim23; navy, Pam18-TM). The view is from the matrix side. Red arrowhead, putative polypeptide exit site. Dashed line, the plane of the Tim17-Tim23 interface. (**D**) As in C, but showing a cutaway view in the middle of the micelle. Red arrowhead, Tim17 cavity. Numbers, respective TM segments of Tim17 and Tim23. (**E**) As in D, but showing a side view. Red arrow, putative translocation path along the Tim17 cavity (facing back).

### Position of Pam16/18

The Pam16/18 heterodimer is an essential component of the TIM^MOTOR^ complex, which regulates the action of the mtHsp70 ATPase to drive preprotein translocation. To visualize Pam16/18 in the cryo-EM structure, we generated a fusion TIM^MOTOR^ construct, in which Pam18 is fused to Tim17 similarly to a previous report (18), and determined its cryo-EM structure at 3.2 Å resolution (Table 1; and Fig. S11). Although Pam16/18 was not resolved to high resolution, this structure nonetheless revealed that Pam16/18 is positioned next to TMs 3 and 4 of Tim17 (Fig. 5 C–E).

Two additional low-resolution features were visible in this structure—a globular domain on the matrix side and one TM. We note that the former was also visible in our initial low-resolution reconstruction (Figs. S2 F and H, and S12 A–D). The TM feature, which seems to make only minor contacts with TMs 3 and 4 of Tim17 near the membrane boundaries, is directed to the globular domain on the matrix side (Fig. 5 C–E). The globular domain is most likely the J and J-like domains of the Pam16/18 heterodimer as it is one of the two known globular domains on the matrix side (the other is Tim44-CTD; Fig. S1A). The observed TM is likely the TM of Pam18. Consistent with this, the density of the TM was substantially weaker in the class without the globular domain density (Fig. S11C, class 2; Fig. S12E). These assignments agree well with known interactions between the N-terminal segments of Pam16 and Tim44 and between Pam18 and Tim17 (31, 42, 47, 53, 62). Taken together, these results indicate that the J and J-like domains of Pam16/18 is positioned ∼20–30-Å laterally off the Tim17-facing side of Tim44-CTD. The proximity between the Pam16/18 and the Tim17 cavity likely now explain the previous observation in which a stalled translocating polypeptide can crosslink to Pam16 (42).

Hsp70 binds a substrate polypeptide at high affinity upon ATP hydrolysis. A recent co-crystal structure of bacterial Hsp70 (DnaK) and J-domain (DnaJ) shows that, during ATPase activation, the peptide-binding pocket of the Hsp70 comes in close proximity to the J domain (63). Thus, the observed position of Pam16/18 would place mtHsp70 close to the Tim17 cavity such that it can efficiently grasp the emerging polypeptide (Fig. S12F). This is analogous to the Sec complex, which mediates post-translational protein translocation across the endoplasmic reticulum (ER) membrane in concert with the ER-lumenal Hsp70 called BiP. In the Sec complex, BiP is expected to be oriented by the J-domain of Sec63 such that its peptide-binding pocket is positioned below the polypeptide exit site to the ER lumen (64, 65).

## DISCUSSION

Our findings have major implications for understanding the mechanism of TIM-mediated mitochondrial protein import. First, our analyses indicate that Tim17 constitutes the translocation path for preprotein transport. This finding was highly unexpected because it has been generally assumed that the translocation path would be formed mainly by Tim23, either alone or together with Tim17. Our study now suggests that Tim23 plays primarily a structural role in orienting and stabilizing Tim17. Tim23 may also contribute to TIM function as a platform for Tim44 association and for recruitment of Tim50, the putative receptor for presequences (66). In this regard, it is noteworthy that certain protists, such as *Trypanosoma* and *Giardia*, only possess one homolog of Tim17/Tim22/Tim23, which is substantially more related to Tim17 than to Tim23 (67-69), consistent with our conclusion that the translocation process is mediated by Tim17, not by Tim23.

Second, our structure suggests that TIM-mediated protein translocation does not involve an aqueous channel. Our structure indicates that substrate polypeptides move through the laterally open cavity of Tim17 with part of the polypeptide directly exposed to lipids. Although further investigations will be necessary, we propose that the ‘half-channel’-like Tim17 cavity lowers the energy barrier for polypeptide transport through local membrane thinning (Fig. 6). An analogous mechanism has been recently proposed for the Hrd1 protein retrotranslocase at the ER (70, 71). The ‘permanently open’ lateral gate of Tim17 also explains the ability of the TIM complex to detect a transmembrane sorting signal within preproteins and release it into the lipid phase during translocation, analogous to the lateral release of a stop-transfer signal by an open SecY/Sec61 complex (72-74) (Fig. 6).

**Figure 6.**
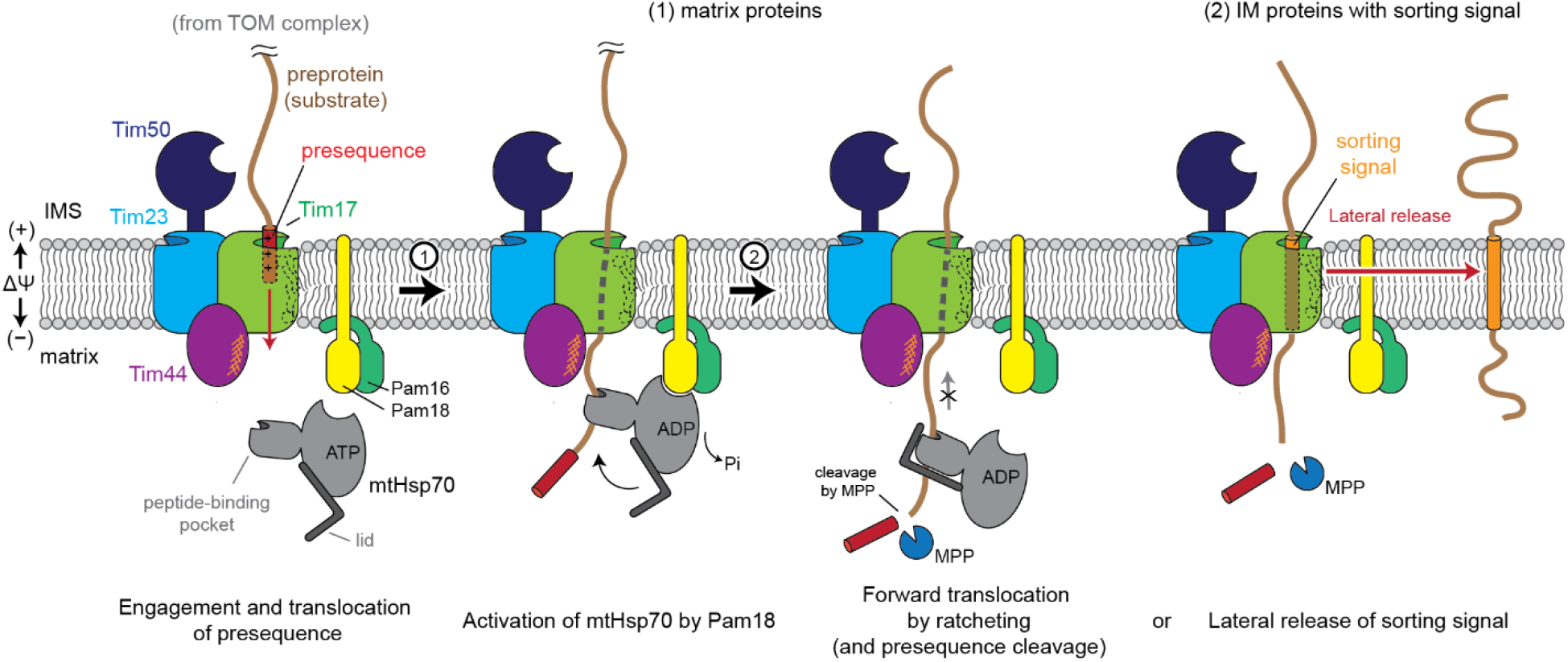
Putative model for the polypeptide transport mechanism of the TIM complex. First, the N-terminal presequence inserts into the cavity of Tim17. This step may be facilitated by Tim50, which may recruit the presequence to the TIM complex. Translocation of the presequence is likely powered by membrane potential (ΔΨ) across the inner membrane. Next, as the N-terminal segment emerges into the matrix, it may first dock on the hydrophobic surface of Tim44-CTD (orange hashes). The presequence could subsequently be grasped by mtHsp70, which would be positioned near the exit of Tim17 (step 1). The interaction between mtHsp70 and the J-domain of Pam18 will trigger ATP hydrolysis and closure of the lid domain, leading to tight binding of mtHsp70 onto the polypeptide (step 2). The tight association of mtHsp70 will promote forward translocation of the polypeptide into the matrix while preventing backsliding. At some point, ADP bound to mtHsp70 will be replaced by ATP by the action of the nucleotide release factor Mge1 (not shown), releasing the polypeptide. Multiple copies of mtHsp70 may bind to the polypeptide simultaneously or successively as translocation progresses (not shown). The presequence will be cleaved by the mitochondrial processing peptidase (MPP; not shown). In the case of preproteins containing a transmembrane sorting signal, the signal will be released into the lipid phase through the open lateral gate of the Tim17 cavity. For clarity, the tethers between the core TIM complex and Pam16–Pam18 and between Tim44 and mtHsp70 are not shown.

Although we cannot exclude the possibility that the TIM complex could form a canonical channel in concert with another protein or by some elaborate conformational changes induced by membrane potential across the mitochondrial IM, we consider these scenarios quite unlikely. Besides the clearly resolved eight TMs from the Tim17–Tim23 heterodimer, the only other TMs in the TIM^MOTOR^ complex are one TM from Tim50, another from Pam18, and two TMs from Pam17 (Fig. S1A), but truncation studies have shown that none of these TMs are essential for protein import (44, 45, 48). Major conformational changes by membrane potential are also unlikely given the lack of an apparent voltage-sensing mechanism, like the highly charged S4 helix of voltage-gated cation channels (75). The requirement of membrane potential for preprotein transport likely originates from a necessity of an electrophoretic force to drive the initial translocation of the positively-charged N-terminal presequence across the IM (76, 77) (Fig. 6).

Our work provides a plausible model for how the core TIM complex coordinates with the mtHsp70 ATPase to provide energy for directional transport of precursor proteins from the IMS to the matrix (Fig. 6). The position of the Pam16/18 heterodimer suggests that the peptide-binding domain of mtHsp70 will be positioned near the polypeptide exit site of Tim17 during the ATPase activation step. This model predicts that the emerging polypeptide may be first stabilized by hydrophobic patches on Tim44 and then will be grasped by mtHsp70. ATP hydrolysis upon activation by the J-domain of Pam18 will close the peptide-binding pocket of mtHsp70 through the action of its lid domain. mtHsp70 can thus act as a molecular ratchet to prevent the preprotein from backsliding into the IMS, in a mechanism analogous to the one first demonstrated for the Sec complex and BiP (78).

In summary, our study reports multiple surprising findings that call for a major revision in the current prevailing model of the TIM complex and its working mechanism. Our work provides a structural framework for interpreting previous results as well as lays the foundation for future investigations. Given the high sequence similarity of the core TIM components, the model we presented here is likely conserved in all eukaryotes. Many questions remain open, including how the TIM complex maintains the permeability barrier during protein transport, how Tim50 associates with Tim23, how the complex converts between the TIM^MOTOR^ and TIM^SORT^ forms, how exactly mtHsp70 interacts with Tim44, Pam16/18, and the translocating polypeptide in different functional states. Addressing these questions will require additional structural and functional studies, including structural determination of a substrate-engaged TIM complex.

## Acknowledgements

We thank D. Toso for electron microscope operational support, J. Thorner for yeast strains and Pgk1 antibodies, and E. Craig for Pam16 and Pam18 antibodies. We thank J. Thorner and T.A. Rapoport for critical reading of the manuscript. This work was supported by the Vallee Scholars Program (E.P.) and Pew Biomedical Scholars Program (E.P.). S.I.S. was supported by an NIH training grant (5T32GM008295) and an NSF Graduate Research Fellowship (DGE 1752814).

## Authors contributions

E.P. conceived and supervised the project; S.I.S. constructed yeast strains and prepared samples for cryo-EM analysis; S.I.S. and E.P. developed anti-Tim44 antibodies, collected and analyzed cryo-EM data, and built atomic models; Y.C. performed mutagenesis experiments; E.P. wrote the manuscript; all authors contributed to data interpretations and manuscript editing.

## Competing interests

Authors declare no competing interests.

## Notes

### Competing Interest Statement

The authors have declared no competing interest.

