## Supplementary material for "Structural basis of mitochondrial protein import by the TIM complex": Materials and Methods, Figures S1 to S12, and Tables S1 to S4

#### Construction of plasmids and yeast strains

A list of yeast strains and plasmids used in this study is given in Tables S1 and S2, respectively. Primers and additional DNA sequences are listed in Tables S3 and S4, respectively.

For purification of the endogenous TIM complex from yeast (Fig. S1C), the *S. cerevisiae* strain yMLT62 (a gift from J. Thorner) was modified to generate the strain ySS078, which expresses C-terminally 2xSpot-tagged Tim17 from the *TIM17* locus. The tag also contains a 16-amino-acid-long linker and an HRV 3C protease cleavage site (amino acid sequence: ...PLQA [Tim17]–ASGTLEVLFQGPT-ASG<u>PDRVRAVSHWSS</u>GGGSGGGST<u>PDRVRAVSHWSS</u>*; Spot-tags are underlined, and the asterisk indicates the C-terminus). To attach the tag to Tim17, a PCR product was first generated containing a 74-bp 5' homology arm immediately before the *TIM17* stop codon, the tag, a hygromycin resistance cassette, and a 74-bp 3' homology arm downstream of *TIM17* (for DNA sequence, see Table S4, TIM17-3C-Spot). DNA was introduced to yeast cells by a standard lithium acetate/polyethylene glycol transformation method. Colonies were selected on a YPD (1% yeast extract, 2% peptone, 2% glucose, and 2% bacto-agar) agar plate containing 400 ug/mL hygromycin B, and chromosomal integration was confirmed by PCR using primers EP_446 and EP_447.

For the overexpression of all TIM complex subunits from an inducible *GAL1* promoter and purification of the full TIM complex (Fig. S1D), the yeast strain ySS025 was used. To construct the ySS025 strain, we first generated two integration plasmids (pSS011 and pSS015) using the MoClo Yeast Tool Kit (YTK) (80). The coding sequences (CDS) for Tim23, Tim17, Tim50, Tim44, Tim21, Pam16, Pam17, Pam18, and Mgr2 were first amplified via PCR using genomic DNA of *S. cerevisiae* BY4741 as a template and cloned individually into a pYTK001 entry plasmid. Internal *Bsa*I and *Bsm*BI sites within CDSs were removed during this step by overlapping PCR. Each CDS part was assembled into expression cassettes in pYTK095 together with connectors, promoter (*pGAL1*), and terminator parts from MoClo YTK and a part for 2xSpot-tag (pYTK-e201). The Tim23, Tim17-Spot, Tim50, Tim44, and Tim21 expression cassettes were inserted into pYTK096 by *Bsm*BI Golden Gate cloning, resulting in the multigene expression plasmid pSS011. Similarly, the Pam16, Pam17, Pam18, and Mgr2 expression cassettes were inserted into pYTK-e106 (ref. 65), resulting in pSS015. The integration plasmids were linearized with the *Not*I endonuclease and sequentially introduced into the yMLT62 yeast strain by a standard lithium acetate/polyethylene glycol transformation method. Colonies were selected on synthetic complete agar medium lacking uracil (SC[–Ura]) for pSS011 or lacking leucine (SC[–Leu]) for pSS015. Chromosomal integration was confirmed by PCR as described previously (80). The yeast strain ySS027, in which Tim21 is Spot-tagged instead of Tim17, is constructed similarly (Tables S1 and S2).

For overexpression and purification of the Tim17/23/44 complex (Fig. S1E), the yeast strain ySS047 was used. To construct the ySS047 strain, we first generated the integration plasmid pSS077 using MoClo YTK. A 2-amino-acid-long GlySer linker at the C-terminus of Tim44 before the stop codon, which was introduced during the construction of the YTK part plasmids, was removed from this expression cassette by PCR, because we found that the inclusion of the GlySer linker somewhat reduces the affinity between Tim44 and the Tim17–Tim23 heterodimer.

The Tim23, Tim17-Spot, and Tim44(–GS) expression cassettes were then inserted into pYTK096 by *Bsm*BI GoldenGate assembly. This integration plasmid was linearized and introduced into the yMLT62 yeast strain, as described above.

For overexpression of the Pam16 and Pam18 subunits in addition to the Tim17/23/44 subunits (Fig. 1A ; and Fig. S3A), the yeast strain ySS055 was used. To construct the ySS055 strain, the Pam16 and Pam18 expression cassettes were inserted into pYTK-e106 to generate the integration plasmid pSS082. This integration plasmid was linearized with *Not*I and introduced into ySS047.

For overexpression and purification of the TIM$^{MOTOR}$ complex containing a fusion between Tim17 and Pam18 (Fig. S11B), the yeast strain ySS107 was used. To make the Tim17–Pam18 fusion construct, a 15-amino-acid-long Gly/Ser linker was introduced between the coding sequences (amino acid sequence: ...PLQA [Tim17] – GGSGGSGGSGGSGGS [linker] – SSQSN... [Pam18]) in the expression cassette. To construct the ySS107 strain, we first generated two integration plasmids pSS107 and pSS109, using MoClo YTK. The Tim23-Spot, Tim17–Pam18, Tim50, Tim44 (–GS), and Pam16 expression cassettes were inserted into pYTK096, resulting in plasmid pSS107. The Pam17-His expression cassette was separately inserted into pYTK-e106, resulting in plasmid pSS109. The integration plasmids were linearized with *Not*I and sequentially introduced into the yMLT62 yeast strain.

For yeast growth complementation assays for Tim17 and Tim23, yeast strains yYC17 and yYC23 were generated, where the chromosomal expression of Tim17 or Tim23 can be repressed in the presence of doxycycline. The tetracycline response element (TRE) and KanMX cassette were PCR amplified from the genome of yeast strain TH_5187 (Horizon Discovery; cat. #YSC1180-202219224) with primers designed for Tim17 (SS_1700 and SS_1701) and Tim23 (SS_1696 and SS_1697). The amplified DNA fragments were then transformed into the R1158 strain (Horizon Discovery; cat. #YSC1210) and selected on YPD agar plates with 300 µg/mL G418. Chromosomal integration was confirmed by PCR with sequencing primers (for Tim17, SS_1702 and SS_1703; for Tim23, SS_1698 and SS_1699).

For expression of Tim17 and its mutants in yeast growth complementation assays (Fig. 3B; and Fig. S7), plasmid pYC17a was used. First, a *Bam*HI cleavage site was introduced before the *Pst*I site of pYTK-e115 by PCR-based mutagenesis. Then, the WT *TIM17* gene, including its native promoter and terminator (426bp upstream and 241bp downstream of CDS), was amplified by PCR (YC_1760 and YC_1761) from the genomic DNA of BY4741 and cloned into this plasmid between the *Eco*RI and *Bam*HI, resulting pYC17a (for DNA sequence, see Table S4). A similar strategy was used to generate pYC23 (288bp upstream and 271bp downstream of Tim23 CDS, PCR with YC_1762 and YC_1763). When indicated, 2 copies of HA tags (amino acid sequence: ...PLQA [Tim17]–TS<u>YPYDVPDYA</u>GS<u>YPYDVPDYA</u>*; HA tags are underlined) were added to the C-terminus by PCR. Point mutations were introduced by PCR-mediated site-directed mutagenesis.

To test Tim17 mutants at variable expression levels (Fig. S8 C-F), pYC17b was constructed to put mutants under the *DDI2* promoter which can be induced in the presence of cyanamide (81). First, a *Kpn*I site was introduced to pYC17a immediately before the start codon of Tim17 by PCR. Then, the native promoter of *TIM17* was replaced with the *DDI2* promoter. To obtain the

*DDI2* promoter, a 938-bp segment upstream of the *DDI*2 CDS was amplified by PCR using YC_1981 and YC_1982 from the genomic DNA of BY4741 and cloned into the *Eco*RI and *Kpn*I site of the modified pYC17a. Lastly, the entire cassette containing the *DDI2* promoter, *TIM17* CDS and *TIM17* terminator (*Eco*RI/*Pst*I fragment) was transferred from the resulting plasmid into pYTK-e122 between the *Eco*RI and *Pst*I sites to generate pYC17b.

For co-immunoprecipitation assays (Fig. 4A), the yeast strain BY4741 was modified to generate the strain ySS121, which expresses C-terminally 2xmyc-tagged Tim23 from the *TIM23* locus and C-terminally 2xHA-tagged Tim 50 from the *TIM50* locus. The myc-tag contains a 16-amino-acid-long linker (amino acid sequence: ...LLEK [Tim23]–ASGTLEVLFQGPTASG-EQKLISEEDLNGEQKLISEEDL*; myc-tags are underlined). To attach the tag to Tim23, a PCR product was generated containing an 87-bp 5’ homology arm immediately before the *TIM23* stop codon, the tag, a hygromycin resistance cassette, and a 66-bp 3’ homology arm downstream of *TIM23* (for DNA sequence, see Table S4, TIM23-myc). The HA-tag contains a 25-amino-acid-long linker and (amino acid sequence: ...AESK [Tim50]–AGGATTASGTGENLYFQGTASGGGS-YPYDVPDYAGSYPYDVPDYA*; HA-tags are underlined). To attach the HA-tag to Tim50, a PCR product was generated containing a 44-bp 5’ homology arm immediately before the *TIM50* stop codon, the tag, a nourseothricin resistance cassette, and a 44-bp 3’ homology arm downstream of *TIM50* (for DNA sequence, see Table S4, TIM50-HA). Yeast cells were transformed sequentially with the PCR products as described above. Chromosomal integration was confirmed by PCR using primers EP_442 and EP_443 for the *TIM23* locus, EP_450 and EP_451 for the *TIM50* locus. The plasmid expressing Spot-tagged Tim17 under the endogenous promoter was made by inserting DNA encoding 426 bp upstream to 241 bp downstream of CDS into the plasmid pYTK-e112 between its *Eco*RI and *Pst*I sites, generating pSS140. A Spot-tag was introduced to the C-terminus of Tim17 by PCR, generating pSS122 (amino acid sequence: ...PLQA [Tim17]–TSPDRVRAVSHWSSGGGSGGGST-PDRVRAVSHWSS*; Spot-tags are underlined). Mutations in Tim17 (D17N/E126Q and D76N/E126Q) were introduced into pSS122 by PCR mutagenesis and verified by Sanger sequencing. The ySS121 yeast strain was transformed with a pSS122 plasmid (encoding WT or respective mutant Tim17).

The Cyb2Δ-DHFR pulldown experiments (Fig. 4 B and C) were performed in yeast strain yYC02, which expresses WT Tim17-HA and WT Tim23-myc from the endogenous loci. Yeast stains yYC03, yYC04, and yYC05 express an additional copy of WT or mutant Tim17-Spot under the endogenous promoter from the *HO* locus (yYC03 for WT Tim17-Spot, yYC04 for D17N/E126Q Tim17-Spot, and yYC05 for D76N/E126Q Tim17-Spot). To generate these strains, a 2xmyc tag was first introduced to the C-terminus of Tim23 in BY4741 as described for ySS121. A PCR product containing a 74-bp 5’ homology arm immediately before the stop codon of Tim17, a 2xHA-tag (amino acid sequence: ...PLQA [Tim17]–AGGATTASGTGENLYFQGTASGGGSYPYDVPDYAGSYPYDVPDYA*; HA tags are underlined and, the asterisk indicates the C-terminus), a nourseothricin resistance cassette, and a 74-bp 3’ homology arm downstream of *TIM*17 was sequentially introduced to the *TIM17* locus (for DNA sequence, see Table S4, TIM17-HA). Chromosomal integration was confirmed with EP_446 and EP_447. To additionally express Spot-tagged Tim17 (WT, D17N/E126Q, or D76N/E126Q), the C-terminally 2xSpot-tagged versions were cloned from respective pSS122 plasmids into the integration vector pYTK-e106 between the *Eco*RI and *Pst*I sites. The integration plasmids were then linearized with *Not*I and introduced to *HO* locus of yeast strain yYC02, resulting in yYC03 (WT), yYC04 (D17N/E126Q), and yYC05 (D76N/E126Q).

For *E. coli* expression of Cyb2Δ-DHFR, DNA encoding the first 165 amino acids of cytochrome b2 (with a.a. 47 to 65 deleted and the endogenous Cys14 and Cys105 mutated to Ser) fused with DHFR was synthesized. This fragment was introduced to pET32a (EMD Millipore) before the His-tag with *Nde*I and *Xho*I to generate pYC002.

To express and purify Tim44-CTD from *E. coli*, the plasmid pSS070 was generated. The coding sequence of Tim44-CTD was amplified via PCR using the genomic DNA of BY4741 as a template and inserted into a pETDuet-1 vector between the *Bam*HI and *Xho*I sites (Tables S2 and S4). To enable affinity purification of the protein, a His-tag (GSSHHHHHHSQDP) and an HRV 3C protease cleavage site (LEVLFQGP) were attached to the N-terminus of Tim44-CTD.

**Purification of the TIM complex for cryo-EM analysis**

Yeast cells were grown in YPEG medium (1% yeast extract, 2% peptone, 2% ethanol and 3% glycerol) in shaker flasks at 30 °C. Cells were harvested upon reaching an optical density of 600 nm ($OD_{600}$) of ~1.5–2. For purification of the endogenous TIM complex (Fig. S1C), yeast strain ySS078 was used. For purification of the TIM complex by Tim17-Spot (Fig. S1D), yeast strain ySS025 was used. For purification of the TIM complex by Tim21-Spot (Fig. S1F), yeast strain ySS027 was used. For purification of core $TIM^{MOTOR}$ (Tim17/23/44; Fig. S1E), core+Pam16/18 (Fig. 1A; and Fig. S3A), and the fusion construct (Fig. S11B), yeast strains ySS047, ySS055, and ySS107 were used, respectively. When the TIM complex was overexpressed, yeast cells were induced with 50 nM β-estradiol upon reaching an OD600 of ~0.7–1.0. After 13–15 h of induction, cells were harvested by centrifugation (3,000 *g* for 5 min).

Crude yeast mitochondria were isolated as described previously (82). Cells were washed with deionized water, resuspended in prewarmed DTT buffer (100 mM Tris-$H_2SO_4$, pH 9.4, 10 mM DTT) (2 mL/g wet weight cells), and incubated at 30°C for 30 min to 1 h. Cells were then harvested by centrifugation (3000 *g* for 5 min), washed with lyticase buffer (1.2 M sorbitol, 20 mM potassium phosphate, pH 7.4) (7 mL/g wet weight cells), and resuspended in lyticase buffer containing a crude preparation of homemade lyticase. After incubation at 30°C for 1–2 h with lyticase, cells were harvested by centrifugation (3000 *g* for 5 min), washed with lyticase buffer, and resuspended in ice-cold homogenization buffer (0.6 M sorbitol, 10 mM Tris-HCl, pH 7.4, 1 mM EDTA, 2 mM PMSF) (6 mL/g wet weight cells). All subsequent steps were performed at 4°C. The yeast spheroplasts were then homogenized with 40 strokes of a Dounce homogenizer and diluted two-fold with homogenization buffer. Crude mitochondria were subsequently isolated through steps of differential centrifugation. The suspension was first centrifuged at 1,500 *g* for 5 min to remove cell debris and nuclei and centrifuged again at 4000 *g* for 5 min. The supernatant from this step was then subjected to further centrifugation at 20,000 *g* for 20 min (Beckman JA-17 rotor) to isolate the mitochondrial fraction. The mitochondrial pellet was washed in SEM buffer (250 mM sucrose, 1 mM EDTA, 10 mM MOPS-KOH, pH 7.2) and harvested at 17,000 *g* for 20 min. The supernatant was removed, and aliquots of mitochondrial pellets (~500 mg) were frozen in liquid nitrogen and stored at -80°C until use.

To purify the TIM complex, mitochondrial pellets were first thawed and resuspended in ice-cold lysis buffer (Buffer L; using 3 times volume of the mitochondrial pellet volume) containing 50 mM Tris pH 7.5, 200 mM NaCl, 1 mM EDTA, 10% glycerol, 5 µg/mL aprotinin, 5 µg/mL leupeptin, 1 µg/mL pepstatin A, and 2 mM PMSF. All subsequent steps were carried out at 4°C. The suspension was briefly homogenized with 10 strokes of a Dounce homogenizer. Mitochondria were then solubilized with a detergent mixture of 0.4% glycol-diosgenin (GDN; Anatrace) and 0.6% lauryl maltose neopentyl glycol (LMNG; Anatrace). After a 1.5 h incubation, the lysate was clarified by centrifugation at 20,000 *g* for 30-45 min (Beckman JA-17 rotor), and the supernatant was incubated with Sepharose beads conjugated with anti-BC2 nanobody for 2 h. The beads were then washed with approximately 30 column volumes of a wash buffer (Buffer W) containing 25 mM Tris pH 7.5, 100 mM NaCl, 1 mM EDTA, 0.02% GDN. The TIM complex was eluted by incubating the beads with ~50 µg/mL HRV 3C protease for 1 h. Cleaved protein was concentrated using an Amicon Ultra (cut-off 100k; GE Life Sciences). The sample was then injected into a Superose 6 Increase 10/300 GL column (GE Life Sciences), equilibrated with Buffer W, containing 25 mM Tris pH 7.5, 100 mM NaCl, 1 mM EDTA, 0.02% GDN. Where indicated, anti-Tim44 Fab was added to the TIM complex at a molar ratio ~1:2 (incubated at 4°C for 15 min) prior to size-exclusion chromatography. Peak fractions were pooled and concentrated to ~5–7 mg/mL using an Amicon Ultra. 3 mM fluorinated Fos-Choline-8 (FFC8; Anatrace) was added to the purified TIM sample prior to cryo-EM grid preparation.

**Antibody generation and purification**

Monoclonal antibodies were generated against Tim44 by immunizing mice with purified Tim44-CTD (residues 210–431). BL21(DE3) *E. coli* was transformed with pSS070, and cells were grown at 37°C to an OD600 of 1.5 in Luria Broth supplemented with 100 ug/mL ampicillin. 0.5 mM IPTG was then added to cultures, and cells were grown for additional 3 h at 37°C. After harvesting by centrifugation, cell pellets were washed with water and stored at −80°C until use. The thawed pellets were then resuspended in 50 mM Tris-HCl pH 7.5, 100 mM NaCl, 1 mM EDTA, 10% glycerol, 5 µg/mL aprotinin, 5 µg/mL leupeptin, 1 µg/mL pepstatin A, and 2 mM PMSF and lysed by sonication for 10 min on ice. All subsequent steps were carried out at 4°C. The lysate was clarified by ultracentrifugation (Beckman Type 45 Ti rotor) at 100,000 *g* for 1 h, and the supernatant was collected and incubated with cobalt resin (Thermo Fisher, Cat. #89966) for 2 h. The beads were then washed with approximately 30 column volumes of 50 mM Tris-HCl pH 7.5, 100 mM NaCl, 1 mM EDTA, 10% glycerol, and 10 mM imidazole. Tim44-CTD was eluted by incubating beads with 4 column volumes of 50 mM Tris pH 7.5, 100 mM NaCl, 1 mM EDTA, 10% glycerol, and 200 mM imidazole. The His-tag was subsequently removed from Tim44-CTD by incubating the beads with ~5 µg/mL HRV 3C protease for 1 h. Cleaved protein was concentrated using an Amicon Ultra (cut-off 10k; GE Life Sciences) and injected into a Superdex 200 Increase 10/300 GL column (GE Life Sciences), equilibrated with 50 mM Tris-HCl pH 7.5, 100 mM NaCl, 1 mM EDTA, 10% glycerol. Tim44-CTD was concentrated to ~2 mg/mL, frozen in 50 µL aliquots with liquid nitrogen, and stored at −80°C until use.

Immunization of mice with purified Tim44-CTD and initial screening of positive hybridoma clones by ELISA was performed by Antibodies, Inc (Davis, CA). Briefly, female Balb/c mice were immunized four times each with protein complex combined with Sigma adjuvant System (Sigma Aldrich). 40 µg of Tim44-CTD in adjuvant was used for the first immunization while 20 µg of

Tim44-CTD in adjuvant was used for the subsequent three immunizations. Test bleeds from each of the mice were assayed by ELISA on a Tim44-CTD coated plate and the mouse with the highest antibody titer was intravenously boosted and then sacrificed 4 days later to obtain the spleen which was used in a fusion reaction with the SP2/0 myeloma cell line to create mouse hybridomas. All procedures were approved by Antibodies Incorporated's Institutional Animal Care and Use Committee and performed under their current Office of Laboratory Animal Welfare assurance.

For ELISA screens of hybridoma, the biotinylated core $TIM^{MOTOR}$ complex (Tim17–Tim23–Tim44) was used. After the core complex was purified (in HEPES instead of Tris buffer) and concentrated, a 2.5x molar excess of EZLink NHS-$PEG_4$-Biotin (Thermo Fisher, Cat. #21362) was added to the sample and incubated at 4°C for 16 h. The sample was then injected into a Superose 6 Increase 10/300 GL column (GE Life Sciences), equilibrated with Buffer W. Peak fractions containing the core $TIM^{MOTOR}$ complex were pooled and concentrated to ~0.5 mg/mL. Biotinylation was confirmed by western blotting using StrepTactin-HRP (Bio-Rad, Cat. #1610381). The biotinylated core $TIM^{MOTOR}$ complex was subsequently immobilized to NeutrAvidin coated ELISA plates.

ELISA-positive hybridoma supernatants were further tested by co-immunoprecipitation and immunoblotting experiments to identify clones producing antibodies that could bind to the core $TIM^{MOTOR}$ complex. We could identify three such clones from thirty ELISA-positive clones. One of these (clone #6) was made monoclonal by a limiting dilution method and used for cryo-EM analysis.

To purify IgG, monoclonal hybridoma (clone #6) cells were grown in high glucose DMEM supplemented with 20% v/v fetal bovine serum (Gibco, Cat. #10437028), 8% v/v macrophage-conditioned medium (Antibodies, Inc), 10% v/v NCTC-109 (Gibco, Cat. #21340039), 1x Penicillin-Streptomycin-Glutamine (Gibco, Cat. #10378016), 1x MEM NEAA (Gibco, Cat. #11140050) in a CELLine Flask (Wheaton, Cat. #WCL1000). The antibody supernatant was harvested and centrifuged at 3000 *g* for 30 min at 4°C to remove any insoluble material. Saturated ammonium sulfate ($[NH_4]_2SO_4$) was slowly added to the supernatant to a final concentration of 25% (w/v) and incubated for 5 h at 4°C. The mixture was centrifuged at 3000 *g* for 30 min at 4°C, and saturated $(NH_4)_2SO_4$ was slowly added to the supernatant to a final concentration of ~50% (w/v) and incubated for 16 h at 4 °C. The precipitate containing antibody was collected by centrifugation at 17,000 *g* (Sorvall SS-34 rotor) for 10 min at 4°C. The pellet was resuspended in TBS buffer containing 25 mM Tris-HCl pH 7.5, 80 mM NaCl and dialyzed against 2 changes of 2 L of TBS buffer for 48 h and 1 change of 1 L of 20 mM Tris-HCl pH 8.0 for 16 h in 10-kDa-MWCO dialysis tubing. Dialyzed samples were loaded directly onto a 5-mL HiTrap Q HP anion exchange column (GE Life Sciences) equilibrated in 20 mM Tris-HCl pH 8.0. After washing with 5 column volumes of 20 mM Tris pH 8.0, antibodies were collected during gradient elution to 20 mM Tris pH 8.0, 1 M NaCl. Fractions containing the eluted IgG were pooled (4 mg/mL).

Fab fragments were generated by digestion of IgG with papain (1:25 w/w; Sigma-Aldrich, Cat. #P3125) in buffer containing 20 mM HEPES-NaOH pH 7.0, 150 mM NaCl, 10 mM L-cysteine HCl, 10 mM β-mercaptoethanol, 10 mM EDTA, and additional 9 mM NaOH at 37°C for 3 h. Cleaved antibodies were diluted in 4 L of 20 mM Tris-HCl pH 8.0 for 16 h in 10-kDa-MWCO dialysis tubing. To collect the Fab fragments, the cleaved IgG sample were applied to a 5-mL HiTrap Q HP anion exchange column (GE Life Sciences) equilibrated in 20 mM Tris-HCl pH 8.0. After washing with 5 column volumes of 20 mM Tris-HCl pH 8.0, Fab fragments were collected during gradient elution to 20 mM Tris-HCl pH 8.0, 1 M NaCl. Fractions containing Fab fragments were pooled, diluted with buffer containing 25 mM Tris-HCl pH 7.5, 100 mM NaCl, 2 mM EDTA, and 20% glycerol, and concentrated using an Amicon Ultra (cut-off 30k; GE Life Sciences) to ~4.5 mg/mL. Samples were either stored at 4°C or frozen in liquid nitrogen and stored at -80°C until use. The sequences of the Fab variable domains were determined by sequencing the cDNAs generated from RNAs that were extracted from the hybridoma cell lines according to (83).

**Cryo-EM grid preparation and data acquisition**

To prepare cryo-EM grids, 3 μL of the sample were applied to a glow-discharged (PELCO easiGlow; 0.39 mBar, 25–30 mA, 40–45 s) gold holey carbon grid (Quantifoil R 1.2/1.3, 400 mesh). The grid was blotted for 3–4 s and plunge-frozen in liquid-nitrogen-cooled liquid ethane using Vitrobot Mark IV (FEI) operated at 4°C and 100% humidity. Whatman No. 1 filter paper was used to blot the samples.

The dataset in which all known subunits of TIM complex (TIM$^{full}$) were overexpressed (Fig. S1D) was collected on a Talos Arctica electron microscope (FEI), operated at an acceleration voltage of 200 kV. The Tim17/23/44+Pam16/18 (Fig. 1A; and Fig. S3) and fusion construct (Fig. S11) datasets were collected on a Titan Krios G3i electron microscope (FEI), operated at an acceleration voltage of 300 kV and equipped with a Gatan Quantum Image Filter (slit width of 20 eV). Dose-fractionated images (~50 electrons per Å$^2$ applied over 42 or 50 frames) were recorded on a K3 direct electron detector (Gatan) using the super-resolution mode. All datasets were collected using SerialEM (84). The datasets were collected with image-beam-shift multiple recording (4 to 9 recordings per stage movement), except for the TIM$^{full}$ dataset, which was collected without image shift. Coma induced by image-beam shift was corrected by beam tilt compensation in SerialEM. The physical pixel size was 1.14 Å for the Arctica dataset and 1.05 Å for the Krios G3i datasets. Target defocus values were set from −0.7 to –2.6 μm for the TIM$^{full}$ dataset, −0.8 to –2.1 μm for the Tim17/23/44+Pam16/18 dataset, and −0.6 to –2.5 μm for the fusion construct dataset.

**Cryo-EM structural determination**

Movies were initially preprocessed using Warp (85), by motion-correcting and estimating contrast transfer function (CTF) and defocus parameters with 7-by-5 tiling (5-by-5 for the TIM$^{full}$ dataset). Micrographs were manually inspected to remove micrographs that were not suitable for image analysis, largely those containing crystalline ice. Particles were automatically picked by Warp and extracted with a box size 256 pixels for the TIM$^{full}$ and fusion construct datasets and 320 pixels for the Tim17/23/44+Pam16/18 dataset. All subsequent image processing was performed in cryoSPARC v2 (86), as described in detail below. While the final maps for the TIM$^{full}$ and the fusion construct datasets were reconstructed from particles picked in Warp, the movies for the Tim17/23/44+Pam16/18 dataset were reprocessed and particles were repicked in cryoSPARC (Fig. S3B). The particles selected from Warp were used to generate initial maps in cryoSPARC, which were subsequently used as a template for the second round of particle picking and heterogenous refinement in cryoSPARC.

(1) TIM$^{full}$ complex

A summary of the single particle-analysis is outlined in Fig. S2B. 854,545 particles were automatically picked in Warp from 2,359 movies. Particles were imported into cryoSPARC for reference-free 2D classification, and classes without clear protein features (mostly empty micelles) were removed. 504,914 particles selected from the 2D classification were subjected to ab initio reconstruction, yielding three initial models (Classes 1 to 3). Class 1 showed a disc-shaped micelle and may represent the TIM$^{SORT}$ complex or the core Tim17–Tim23 complex (Fig. S2D). Class 2 (TIM$^{MOTOR}$) contained two distinguishing bulges (Fig. S2E). The particles in Class 2 (193,357 particles) were used for 3D reconstruction by non-uniform (NU) refinement to yield a map at 8.4-Å resolution (Fig. S2 G and H).

(2) Fab-bound core TIM$^{MOTOR}$ structure (Tim17/23/44 + Pam16/18)

A summary of the single particle-analysis is outlined in Fig. S3B. The initial set of 466,320 particles automatically picked in Warp from 1,506 movies was used for 2D classification. The 268,256 particles selected from the 2D classification (excluding mainly empty micelles) were subjected to ab initio reconstruction, yielding three initial models. Clear features of the core TIM complex and the Fab fragment were visible in one of these classes. These particles (268,256 particles) selected from the 2D classification were classified by a round of heterogeneous refinement using the ab initio reconstructions. The resulting particles (174,222 particles) were used for 3D reconstruction by NU refinement, yielding a map at 2.8-Å resolution for the core+Pam16/18 dataset.

Raw movies (1,506 movies) were then imported into cryoSPARC for tile-based motion correction and CTF estimation. 1,485 micrographs were selected, and a total of 689,333 particles were picked with lowpass filtered templates generated from the 3.1-Å-resolution 3D reconstruction of the TIM complex. Particles were extracted with a box size of 320 pixels, Fourier-cropped to 160 pixels, and subjected to a round of 2D classification. Selected particles from 2D classification (333,513 particles) were subjected to ab initio reconstruction, generating four initial models, and heterogeneous refinement. 188,938 particles were classified into Class 1, which showed clear features of the TIM–Fab complex. This final set of particles (159,958 particles) was subjected to NU refinement and local CTF refinements to yield a map at 2.7-Å resolution.

(3) Fab-bound TIM$^{MOTOR}$ complex with Tim17-Pam18 fusion (fusion construct)

A summary of the single particle-analysis is outlined in Fig. S11C. 866,976 particles were automatically picked in Warp from 2,915 movies. Particles were imported into cryoSPARC for reference-free 2D classification, and classes without clear protein features (mostly empty micelles) were removed. 361,263 particles selected from the 2D classification were subjected to ab initio reconstruction, yielding three initial models (Classes 1 to 3). Only Class 2 displayed good TIM–Fab features (Fig. S11C). The particles in Class 2 (222,430 particles) were subjected to two additional rounds of heterogenous to remove low-quality particles. The resulting 175,317 particles could be refined to a 3.2-Å-resolution reconstruction by NU refinement. Then the particles were further classified into two sets, one with and the other without the Pam16/18 feature by heterogenous refinement. The input reference map without the Pam16/18 feature was made by manually erasing the Pam16/18 density in UCSF Chimera (87). This procedure separated 96,650 particles into a class with a stronger density of the Pam16/18 feature and 78,667 particles into a class with substantially weaker Pam16/18 density. These particle sets were separately used for 3D reconstruction by NU refinement to yield maps at 3.2-Å and 3.3-Å resolution, respectively.

**Atomic Model Building**

The initial atomic model was built de novo into the sharpened map of the core TIM complex dataset using Coot (88). The model for the fusion construct dataset were built after rigid-body fitting into the corresponding map using the initial atomic model and rounds of local refinement in Coot. The Pam18 transmembrane helix, visible in the fusion construct structure, was modeled with alanine because amino acids could not be unambiguously registered. The following sequences were not modeled in the Tim17/23/44+Pam16/18 structure because they were poorly resolved in the density maps: N to 5 (Tim17), 139 to C (Tim17), N to 85 (Tim23), 221 to C (Tim23), N to 106 (Tim44), and 194 to 254 (Tim44). Residue side chains with poor density were truncated at the β-carbon.

Model refinement was performed using Phenix (*phenix.real_space_refine*) (89) with the refinement resolution limit set to the overall resolution of the map. Cryo-EM maps were sharpened using the Phenix Auto-sharpen tool (*auto_sharpen*) (90) (Table 1) and used for refinement. Structural validation was performed using MolProbity (91) in the Phenix package. Protein electrostatics were calculated using the Adaptive Poisson-Boltzmann Solver (92) with default parameters (with monovalent ion concentrations of 0.15 M each) built in PyMOL (Schrödinger). UCSF Chimera, Chimera X (93), and PyMOL were used to prepare structural figures in the paper.

**Multiple sequence alignment**

Amino acid sequences of Tim17 and Tim23 from various organisms were obtained from the UniProt Reference Clusters (UniRef100, 2021_03). For Tim17, any sequence smaller than 130 amino acids or larger than 256 amino acids were excluded. For Tim23, any sequence smaller than 166 amino acids or larger than 257 amino acids were excluded. A total of 94 sequences of Tim17 and 102 sequences of Tim23 were separately aligned using MAFFT (94). Amino acid identity was mapped onto the structure using UCSF Chimera.

**Yeast growth complementation assay (spot assay)**

Yeast strains yYC17 and yYC23 were transformed with plasmid pYC17a and pYC23, respectively. These plasmids encode Tim17 and Tim23 under its endogenous promoter, respectively (Fig. 3 B and D; and Fig. S7). Cells were selected on YPD agar plate with 100 µg/mL nourseothricin. Single colonies were picked and grown overnight in YPD medium with 100 µg/mL nourseothricin at 30°C. The next morning, the overnight culture was then diluted to $OD_{600}$~0.2 in YPD with 100 µg/mL nourseothricin and grown until $OD_{600}$ reached 0.7–1.5. Cells were pelleted and resuspended in fresh medium to $OD_{600}$ of 0.1. After 5-fold serial dilution, 10 μL were spotted on YPD/nourseothricin agar plates with or without 10 μg/mL doxycycline. The plates were incubated at 30°C or 37°C for two days before imaging.

For complementation assays in Fig. S8C, yYC17 was transformed with pYC17b encoding Tim17 mutants under the *DDI2* promoter. Cells were selected on synthetic complete agar plates containing 2% glucose and lacking leucine (SC[−Leu]). Single colonies were picked and grown in the SC(−Leu) medium overnight, back-diluted and grown to $OD_{600}$ ~0.7-1.5. Cells were resuspended to $OD_{600}$ of 0.1 with fresh medium. After 5-fold serial dilution, 10 uL were spotted on SC(−Leu) agar plates with or without 10 μg/mL doxycycline. When indicated, cyanamide (Alfa Aesar, Cat# L2044822) was included in the medium to induce protein expression. The plates were incubated at 30°C or 37°C for two days before imaging.

**Co-Immunoprecipitation of Tim and Pam subunits**

For the co-immunoprecipitation experiments shown in Fig. 4A, ySS121 strains harboring plasmid pSS122, a CEN/ARS plasmid that constitutively expresses WT or mutant Tim17 under the endogenous *TIM17* promoter, were grown in SC(–Leu) containing 0.25% glucose, 2% ethanol, and 3% glycerol at 30°C. The yeast cultures ($OD_{600}$ ~0.8) were then washed in water and diluted to an $OD_{600}$ of 0.05 and grown overnight in YPEG medium at 30°C to an $OD_{600}$ of ~0.8-1.0. Crude mitochondria were prepared as described above, and mitochondria were resuspended in ice-cold lysis buffer containing 50 mM Tris-HCl pH 7.5, 200 mM NaCl, 1 mM EDTA, 10% glycerol, 5 μg/mL aprotinin, 5 μg/mL leupeptin, 1 μg/mL pepstatin A, and 2 mM PMSF and solubilized with 1% digitonin (Calbiochem, Cat. #300410) (the lysate volume was approximately 10 times the mitochondrial pellet volume). All subsequent steps were carried out at 4°C. After a 1 h incubation, the lysate was clarified by centrifugation at 17,000 *g* for 30 min, and the supernatant was incubated with Sepharose beads conjugated with anti-BC2 nanobody for 1 h. The beads were washed four times with 1 mL of a wash buffer containing 25 mM Tris-HCl pH 7.5, 100 mM NaCl, 1 mM EDTA, 10% glycerol, and 0.1% digitonin. Proteins were eluted by addition of SDS sample buffer and heating samples for 30 min at 60°C before analyzsis by SDS-PAGE and western blotting.

**Pulldown with Cyb2Δ-DHFR-stalled translocation intermediates**

C-terminally His-tagged Cyb2Δ-DHFR was purified with a procedure adapted from (60). Briefly, *E. coli* BL21 (DE3) was transformed with pYC002 encoding Cyb2Δ-DHFR. Cells were grown Luria broth (LB) medium to $OD_{600}$ of ~0.6, and the expression was induced with 0.5 mM isopropylthio-β-galactoside (IPTG). After growing the culture overnight at 10°C, cells were harvested, resuspended in 40 mM Tris-HCl, pH 7.5, 300 mM NaCl, 1 mM PMSF, 10 mM DTT, 0.1% Triton-100) and lysed with sonication. The cell lysate was cleared by centrifugation at 14,000 rpm for 1 h at 4°C (Sorvall SS-34 rotor), and Cyb2Δ-DHFR was purified with cobalt resin (Thermo Scientific, Cat. #89966). The eluate was dialyzed against 0.1M phosphate buffer, pH 6.5 and injected into a HiTrap SP HP column (GE Life Science). The Cyb2Δ-DHFR was eluted by a NaCl gradient. Purified Cyb2Δ-DHFR was concentrated to 0.25 mg/mL in storage buffer (10mM MOPS-KOH, pH 7.2, 150 mM NaCl, 1 mM EDTA, 10% glycerol), frozen, and stored at –80°C.

To isolate mitochondria, yeast strains (yYC03, yYC04, and yYC05) expressing WT or mutant Tim17-Spot from the *HO* locus, in addition to Tim17-HA and Tim23-myc from their respective endogenous loci, were grown to $OD_{600}$ ~0.8–1.0 in YPEG medium. Crude mitochondria were prepared as described above, and frozen at –80°C as 5 mg/mL (mitochondrial protein concentration as determined by Bradford assay [Pierce, Cat #23200]) aliquots in SEM buffer (250 mM sucrose, 1 mM EDTA, 10 mM MOPS-KOH, pH 7.2) until use. As a control, mitochondria from yYC02 (strain without Tim17-Spot) were prepared in the same manner.

To generate stalled translocation intermediates, 200 μg of mitochondria (mitochondrial protein) were mixed with 4 μg Cyb2Δ-DHFR in 400 μL import buffer (3% w/v bovine serum albumin [BSA], 250 mM sucrose, 80 mM KCl, 10 mM MOPS-KOH, pH 7.2, 5 mM $MgCl_2$) and supplemented with 2 mM ATP, 2 mM NADH and 2 μM methotrexate (MTX; TCI America, Cat. #M16641). Where indicated, mitochondria were pre-incubated with 2 μM valinomycin (Sigma Aldrich, Cat. #V0627) for 5 min on ice to dissipate the membrane potential, before setting up the import reactions. After the 20-min incubation at 25°C, mitochondria were pelleted and washed twice with sucrose-MOPS buffer (250 mM sucrose, 10 mM MOPS-KOH, pH 7.2). Then, mitochondria were resuspended in 220 uL of 10 mM MOPS-KOH, pH 7.2, 10% glycerol, 200 mM NaCl, 10 mM imidazole, 5 μg/mL aprotinin, 5 μg/mL leupeptin, 1 μg/mL pepstatin A, 2 mM PMSF and solubilized with 1% digitonin for 45 min at 4°C. The lysates were cleared by centrifugation, and the supernatants were incubated with 20 μL Ni-charged resin (GenScript, Cat. #L00223) for 1 h at 4°C. The resin was washed three times with 1 mL wash buffer containing 10 mM MPOS-KOH, pH 7.2, 10% glycerol, 200 mM NaCl, 20 mM imidazole, 0.02% digitonin. Bound proteins were eluted with 40 μL of 10 mM MPOS-KOH, pH 7.2, 10% glycerol, 200 mM NaCl, 250 mM imidazole, 0.02% digitonin. Samples were separated by SDS-PAGE and analyzed by western blotting with antibodies.

#### Western blotting and antibodies

To examine the expression level of the Tim17 mutants, $OD_{600}$ ~0.1 of cells were resuspended in fresh medium (YPD with 100 μg/mL nourseothricin for endogenous promoter constructs; SC[−Leu] with cyanamide at indicated concentration for *DDI2* promoter constructs) and grown overnight at 30°C. Equal numbers (2.5 ODs) of cells were harvested and lysed with lysis buffer (0.1 M NaOH, 0.05 M EDTA, 2% SDS, 2% β-mercaptoethanol). The samples were neutralized with acetic acid and analyzed by SDS-PAGE.

Western blotting experiments were performed with antibodies against Pam16 (a gift from E. Craig), Pam18 (a gift from E. Craig), Pgk1 (a gift from J. Thorner), Myc-tag (Invitrogen, Cat# 13-2500), HA-tag (Invitrogen, Cat# 26183), His-tag (Invitrogen, Cat# MA1-21315). For anti-Tim44 blots, we used the non-monoclonal hybridoma supernatant of Clone #23, which could detect Tim44 on western blots. For Spot-tag blots, we used home-made BC2-Nb fused with a rabbit Fc domain, which was produced in HEK293 cells by transient transfection.

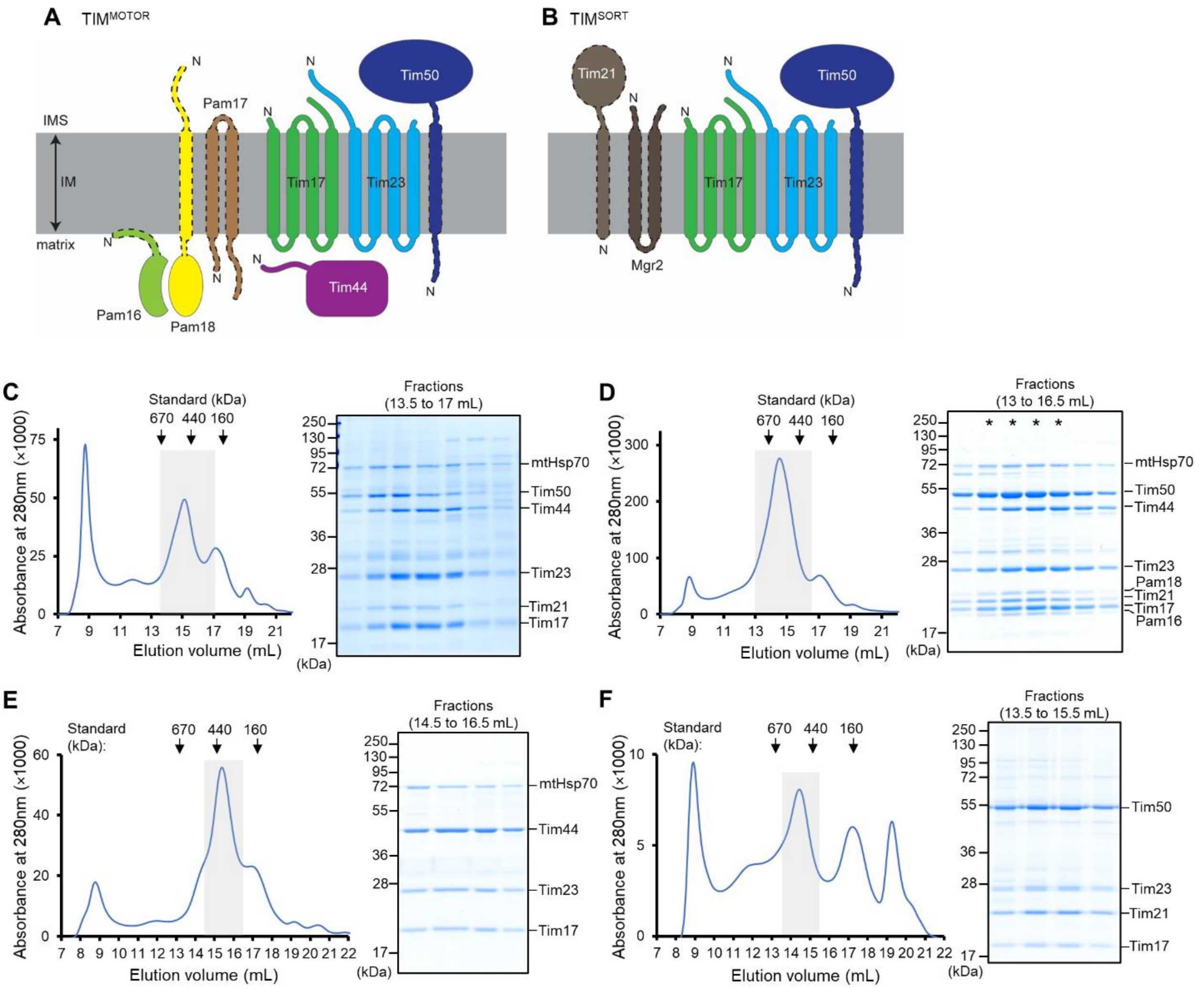

**Fig. S1. Purification of the TIM complex from *S. cerevisiae*.**
(**A**) Schematic diagram of the putative composition of the TIM$^{MOTOR}$ complex. The N-terminus of each subunit is indicated by an "N". All subunits except for Pam17 are essential proteins in yeast. Individual deletions of the portions of Pam16, Pam17, Pam18, and Tim50 outlined by a dashed line are known to be non-lethal (44, 45, 48). Note that a previous report has suggested that Pam17 may not be a component of the mature TIM$^{MOTOR}$ complex (18). (**B**) Schematic diagram of the putative composition of the TIM$^{SORT}$ complex. Tim21 and Mgr2 are nonessential proteins in yeast. (**C**) Purification of the endogenous TIM complex. After affinity purification via Spot-tagged Tim17, the sample was subjected to Superose-6 size-exclusion chromatography (SEC). The gray box indicates fractions further analyzed by non-reducing SDS-PAGE and Coomassie staining (right panel). We note that copurified Tim50 often did not fully co-migrate with the other subunits in SEC, indicative of its dissociation over time. (**D**) As in C, but purification of the TIM complex from mitochondria co-overexpressing nine known subunits of the TIM complex except for mtHsp70. Fractions marked with asterisks are pooled and used for cryo-EM analysis. (**E**) As in D, but purification was performed with mitochondrial overexpressing only Tim17 (Spot-tagged), Tim23, Tim44. (**F**) As in D, but purification via Spot-tagged Tim21.

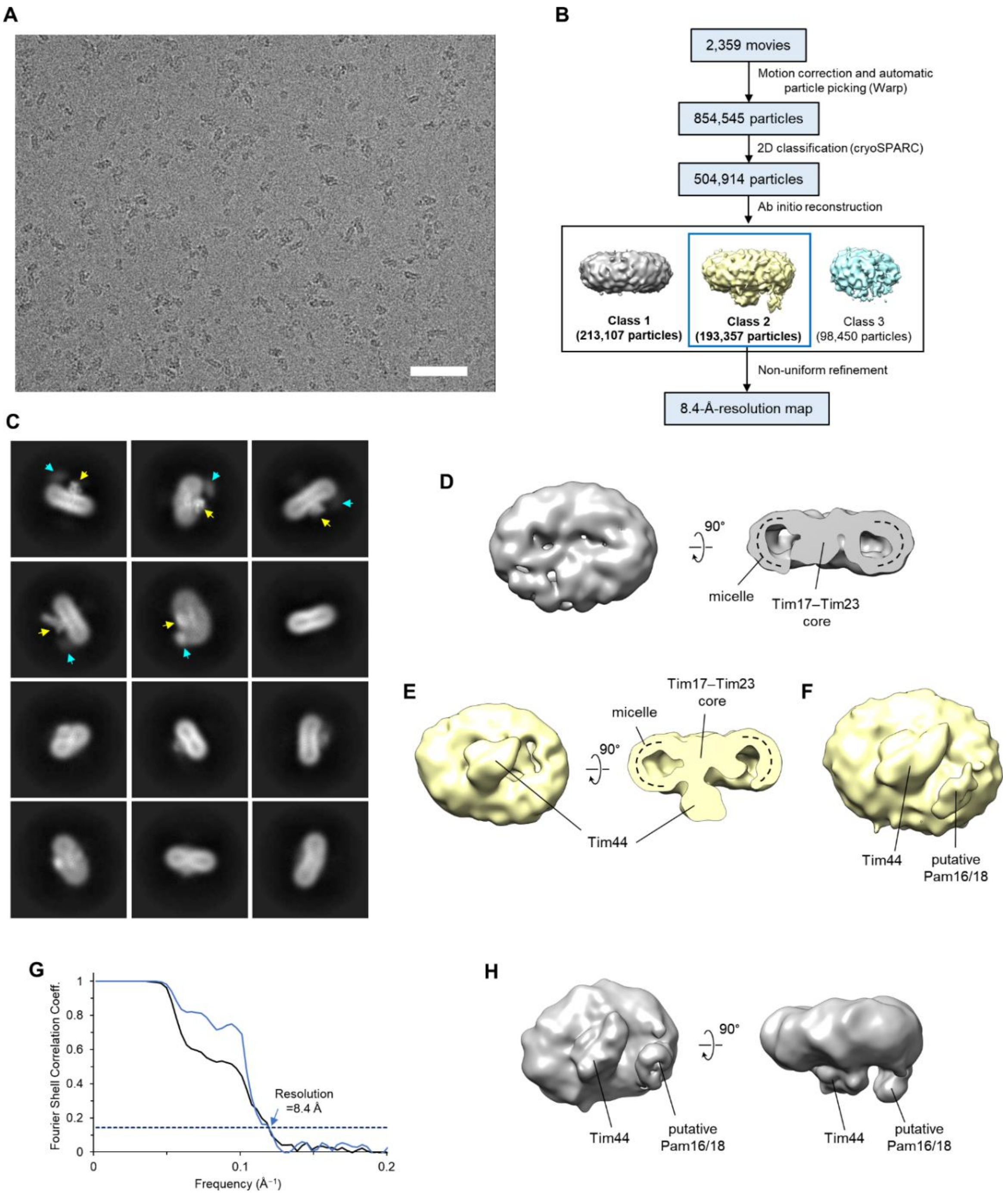

**Fig. S2. Low-resolution cryo-EM analysis of the TIM complex.**
(**A** and **B**) Representative cryo-EM image (A) and summary of single particle analysis (B) of the sample shown in Fig. S1D (co-overexpression of nine subunits). In panel A, scale bar is 50 nm. We note that other TIM samples behaved similarly. (**C**) Example two-dimensional (2D) classes of the TIM complex. Yellow arrowhead, central protruding feature (Tim44-CTD); cyan arrowhead, additional protruding feature (Pam16/18). (**D**) Class 1 from ab-initio refinement of particles shown in B. (**E**) As in D, but showing Class 2. (**F**) As in the left panel of E, but shown at a lower isosurface threshold value. (**G** and **H**), Fourier shell correlation (G) and density map (H) of the 8.4-Å-resolution final reconstruction of the Class 2 particles.

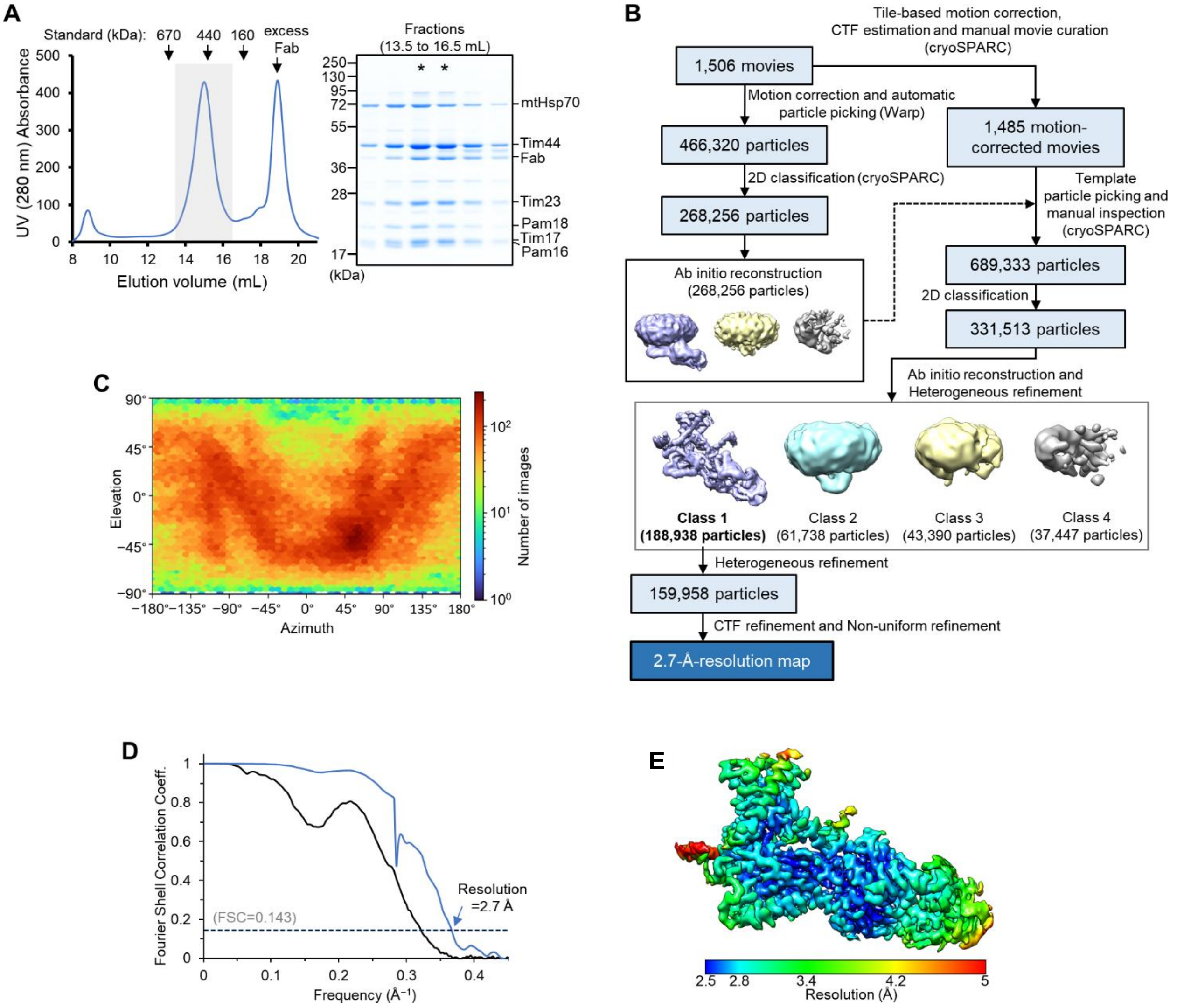

**Fig. S3. High-resolution cryo-EM structure of the TIM$^{MOTOR}$ complex (Tim17/23/44 + Pam16/18).**
(**A**) Purification of the Fab-bound TIM$^{MOTOR}$ complex. Left, Superose-6 SEC elution profile; right, Coomassie-stained non-reducing SDS gel of fractions indicated in the left panel by a gray box. Fractions marked with asterisks were pooled and used for cryo-EM analysis. Note that in this sample preparation, Tim50 was excluded from co-expression. We also note that throughout our cryo-EM study, we did not observe clear Tim50 features even when Tim50 was included in purified complexes. (**B**) Summary of single particle analysis of the Fab-bound TIM$^{MOTOR}$ complex (Tim17/23/44 + Pam16/18). (**C**–**E**) Particle view distribution (C), Fourier shell correlation (D), and local resolution distribution (E) for the Fab-bound TIM$^{MOTOR}$ complex.

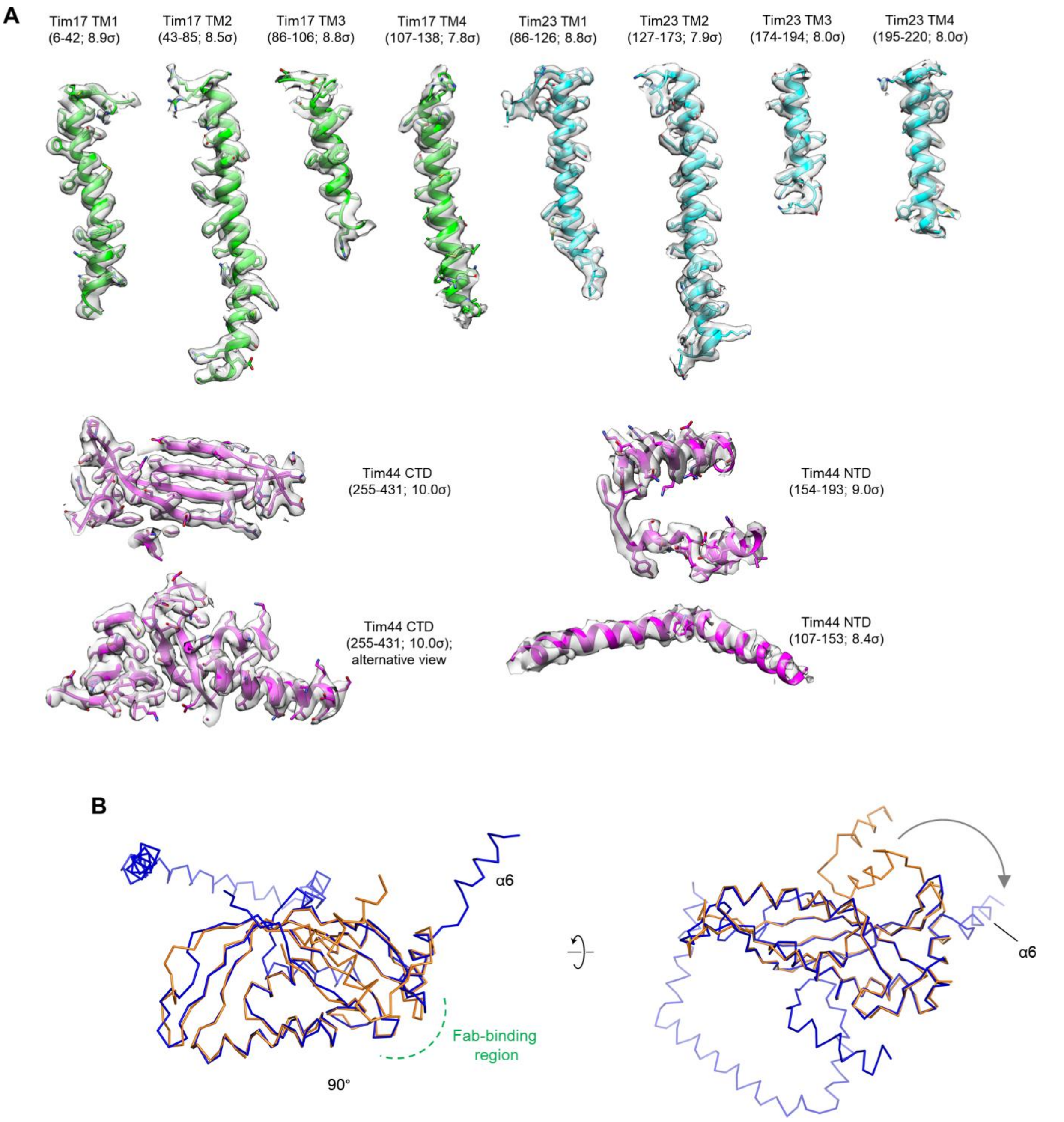

**Fig. S4. Atomic model of the core TIM complex.**
(**A**) Segmented EM densities and atomic models of the TIM complex (Tim17/23/44+Pam16/18). The amino acid ranges and isosurface threshold values are indicated. (**B**) Superposition between the Tim44-CTD structure from the core TIM$^{MOTOR}$ complex (blue) and the crystal structure of isolated Tim44-CTD (orange; PDB ID: 2FXT). The Fab-binding region is indicated with a green dashed line. Note that in the crystal structure, a segment immediately preceding the CTD (including α6) is in different position from the present cryo-EM structure (also see Fig. S10 A and B).

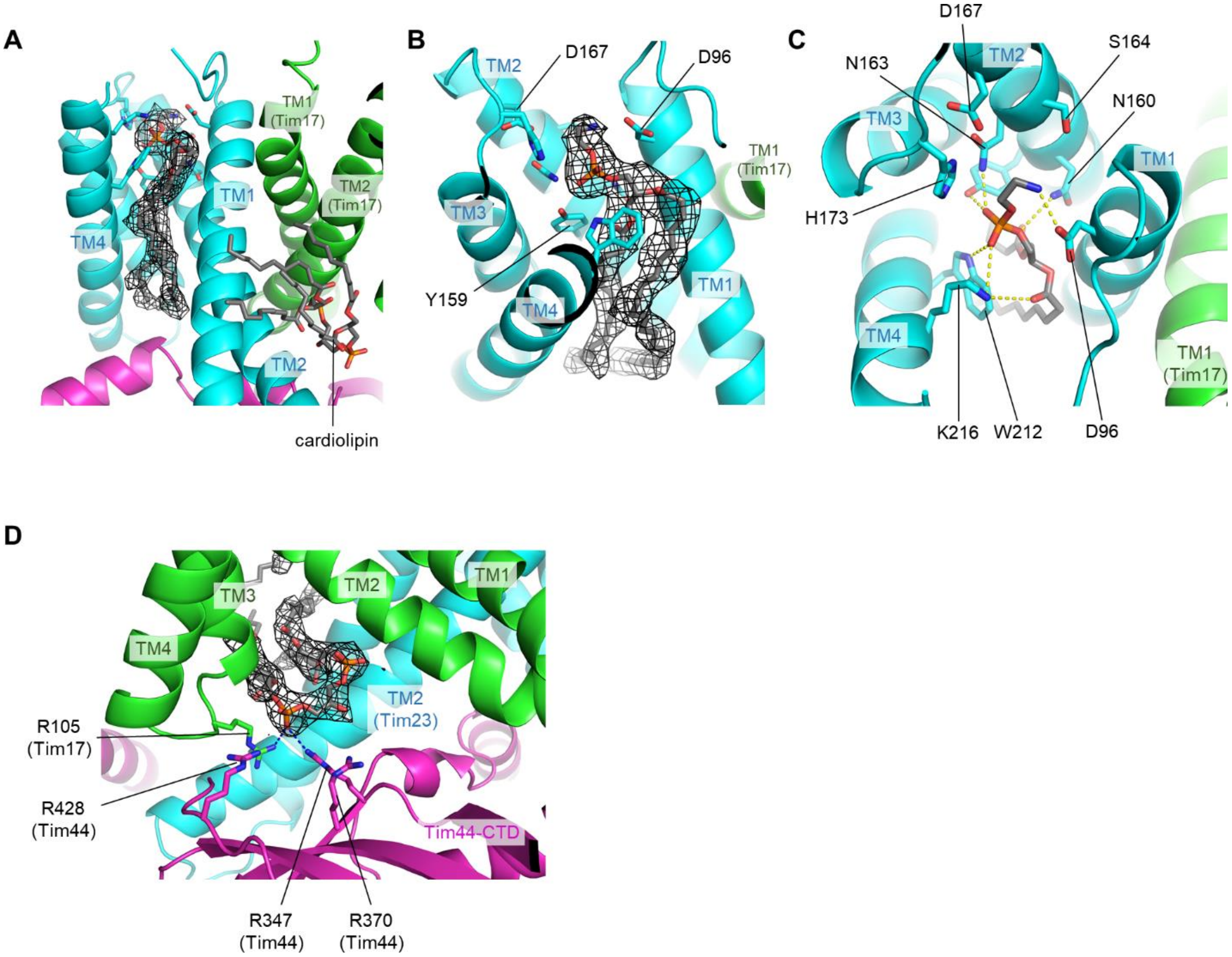

**Fig. S5. Lipid interactions in the TIM complex.**
(**A**–**C**) Phospholipid bound to the cavity of Tim23. Shown are a side view into the cavity (A), a tilted view (B), and a view from the IMS (C). The EM density of the phospholipid is shown as black mesh. Side chains contacting the lipid head group are shown as sticks. In C, polar interactions are indicated by yellow dashed lines. We note that, although the identity of the phospholipid could not be unambiguously determined, the EM density agrees very well with phosphotidylethnaolamine. (**D**) As in Fig. 1E, but showing a zoomed-in view for the cardiolipin molecule. The EM density is shown in black mesh. Polar interactions are indicated by dashed line.

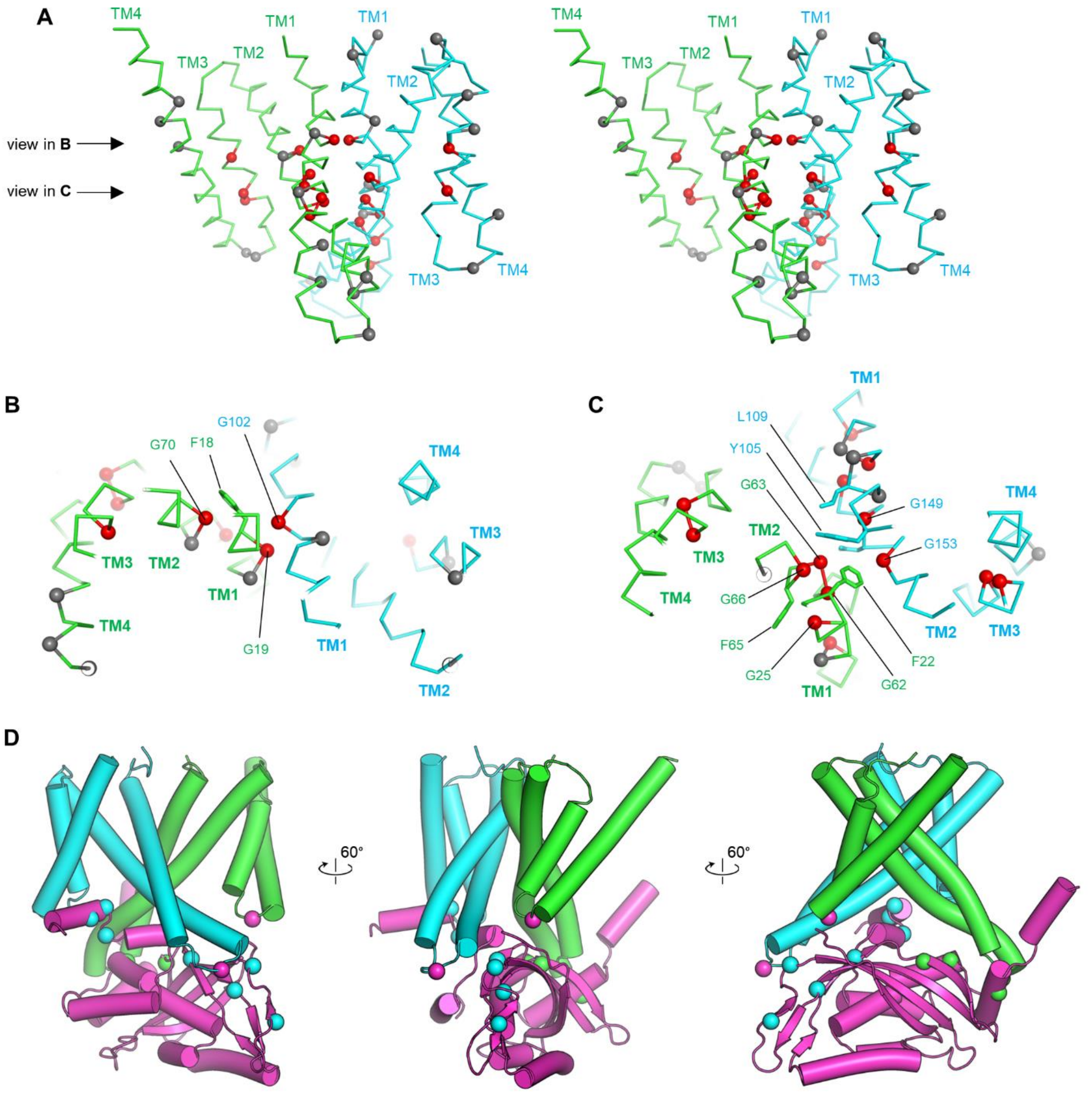

**E**

| Position of Bpa | Crosslinked subunit | Min. distance in structure to crosslinked subunit | References |
|---|---|---|---|
| Tim17-G106 | Tim44 | 3.4 Å (Tim44-Arg428 NH1) | (49) |
| Tim23-N127 | Tim44 | 8.3 Å (Tim44-Glu390 OE2) | (52) |
| Tim44-S160 | Tim23 | 8.9 Å (Tim23-Met204 CE) | (52) |
| Tim44-I163 | Tim23 | 5.2 Å (Tim23-Met204 CE) | (52) |
| Tim44-D165 | Tim23 | 4.1 Å (Tim23-Lys190 NZ) | (52) |
| Tim44-D345 | Tim23 | 3.5 Å (Tim23-Asn139 CD2) | (38) |
| Tim44-I374 | Tim23 | 6.2 Å (Tim23-Leu132 CD2) | (38) |
| Tim44-I385 | Tim23 | 12.5 Å (Tim23-Pro129 CG) | (38) |
| Tim44-A391 | Tim23 | 3.7 Å (Tim23-Leu132 CD1) | (38) |
| Tim44-S271 | Tim17 | 4.1 Å (Tim17-Arg46 NH2) | (38) |
| Tim44-R272 | Tim17 | 8.2 Å (Tim17-Leu43 CG) | (38) |
| Tim44-N286 | Tim17 | 5.5 Å (Tim17-Asn40 CB) | (38) |
| Tim44-E350 | Tim17 | 6.1 Å (Tim17-Arg57 CD) | (38) |

**Fig. S6. Inter-subunit contacts among Tim17, Tim23, and Tim44.**

(**A**) Stereo view showing the positions of glycine in Tim17 (green Cα-trace) and Tim23 (cyan Cα-trace). Positions of all glycine residues are shown as spheres (Cα atoms). Red spheres indicate positions in which mutation to a non-glycine amino acid would cause steric clashes. Gray spheres indicate positions in which mutation may not cause steric clashes. Shown is a side view along the plane of the Tim17–Tim23 interface. (**B** and **C**) As in A, but viewing down the planes indicated by arrows in A from the IMS. (**D**) Mapping of amino acid positions that have been shown to produce an inter-subunit photo-crosslinking with *p*-benzoyl-L-phenylalanine (Bpa). Tim17 (green), Tim23 (cyan), and Tim44 (purple) are shown in a cylindrical cartoon representation. Green and cyan spheres are positions in Tim44 that crosslink to Tim17 and Tim23, respectively. Purple spheres are positions in Tim17 and Tim23 that crosslink to Tim44. See panel E for details. (**E**) Summary of previously reported inter-subunit photocrosslinking results. The minimal distance is between the position (Cα for Tim17-G106 and Cβ atom for all other positions) into which Bpa was incorporated and the crosslinked subunit (atom is specified).

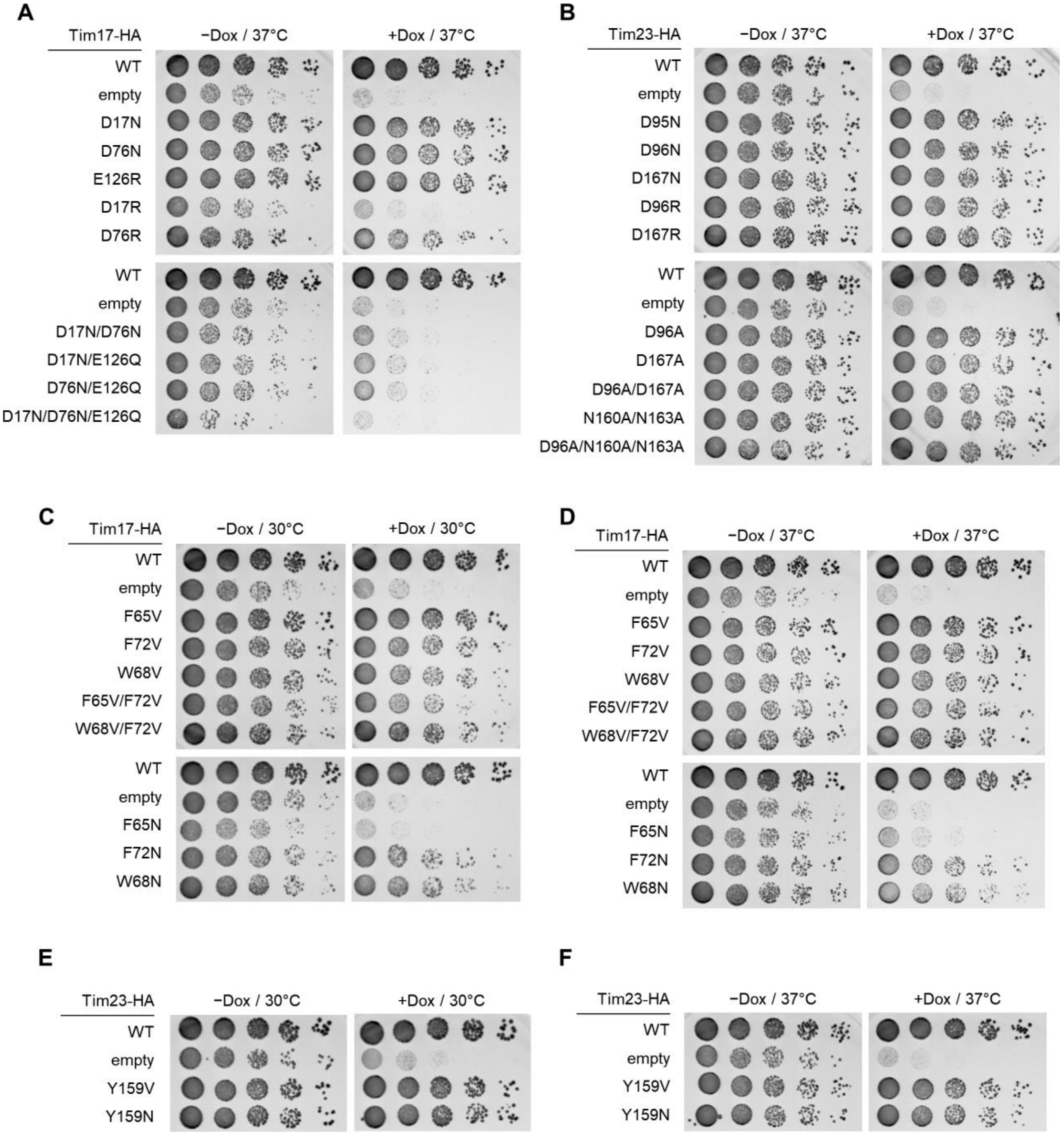

**Fig. S7. Mutational analysis of cavity-lining amino acids.**
(**A** and **B**) As in Fig. 3 B and D, but Tim17 (panel A) and Tim23 (panel B) mutants were tested at 37°C. Note that we have detected no clear growth defects with all indicated Tim23 mutants. (**C** and **D**) As in A, but testing mutations on aromatic residues lining the Tim17 cavity (Phe65, Phe72, and Trp68) of Tim17 at 30°C (panel C) and 37°C (panel D). (**E** and **F**) As in C and D, but testing Tyr159 of Tim23.

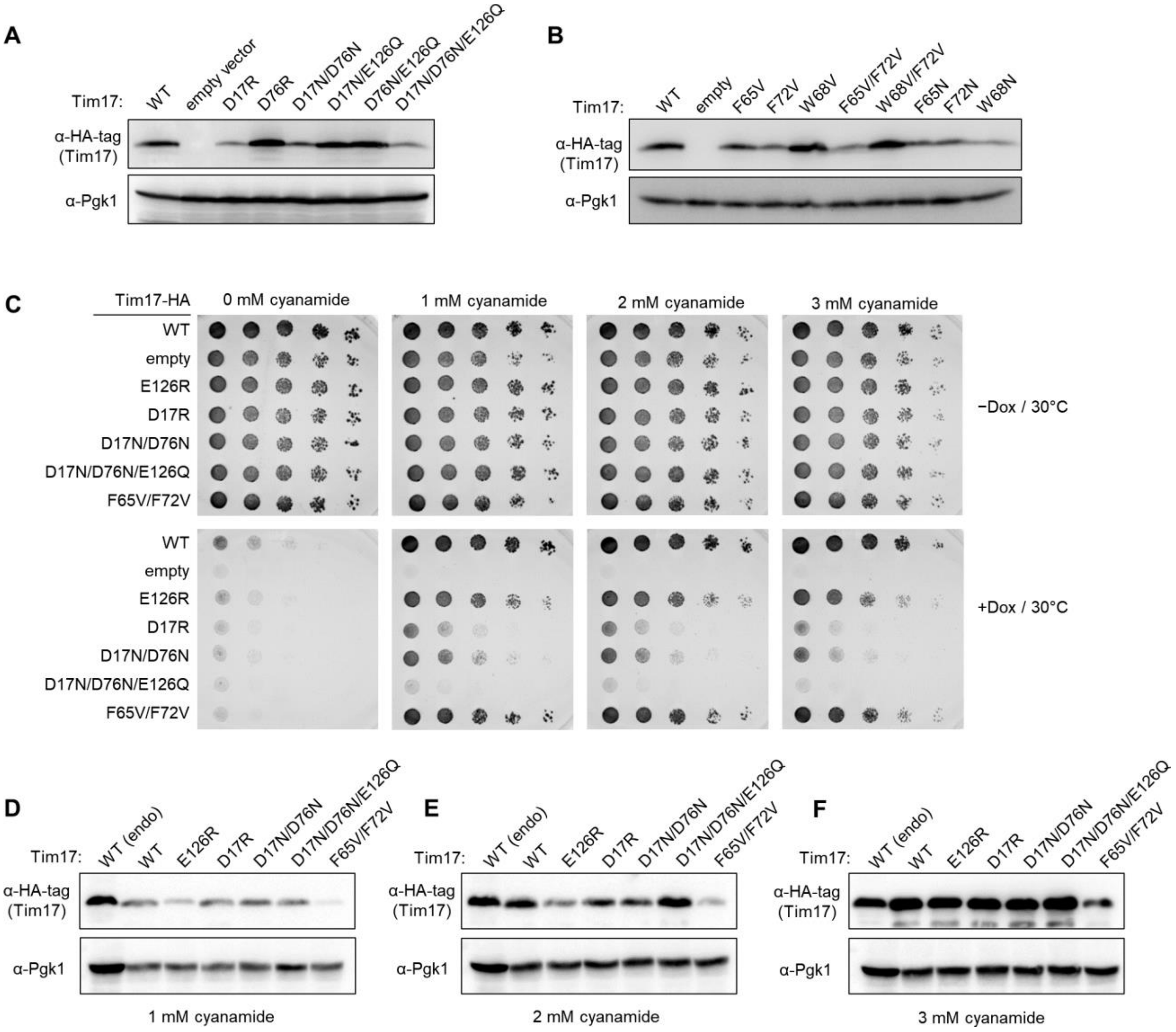

**Fig. S8. Expression levels of Tim17 mutants.**

(**A**) Expression levels of WT and indicated mutants were measured by western blotting. All Tim17 variants were expressed from the endogenous promoter. 3-phosphoglycerate kinase (Pgk1) was used as a loading control. (**B**) As in A, but with additional mutants. (**C**) As in Fig. 3 B and D, the experiments were performed with a 2μ plasmid expressing indicated mutants of HA-tagged Tim17 under a cyanamide-inducible *DDI2* promoter. Cells were spotted on synthetic complete without leucine (SC[–Leu]) plates supplemented with varying concentrations of cyanamide. The plates contain doxycycline (Dox), where indicated. (**D**–**F**) As in A, but measuring expression levels of Tim17 under the *DDI2* promoter in the presence of varying concentrations of cyanamide. As a control, expression of WT Tim17 under the endogenous promoter (endo) from a CEN/ARS plasmid was included.

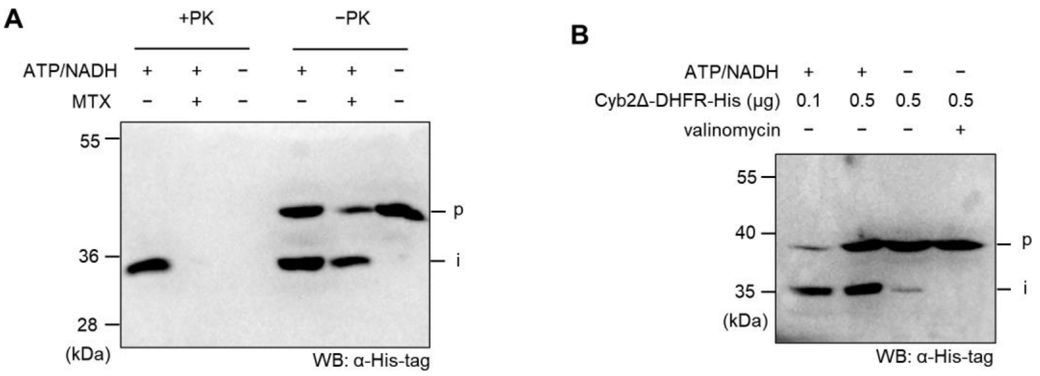

**Fig. S9. Generation of a translocation intermediate complex.**
(**A**) WT mitochondria (50 μg proteins) were incubated with 1 μg of purified Cyb2Δ-DHFR-His in a 100-μL reaction volume. Where indicated, ATP/NADH (2 mM each) and/or methotrexate (MTX; 2 μM) were included. After the 20-min incubation at 25°C, mitochondria were washed, and the reactions were split. One group was treated with proteinase K (+PK) and the other was not (−PK). The samples were quenched with phenylmethylsulfonyl fluoride and analyzed by SDS-PAGE and western blotting (WB) with anti-His-tag antibody. p, precursor form of Cyb2Δ-DHFR-His; i, intermediate (presequence-cleaved) form of Cyb2Δ-DHFR-His. (**B**) Additional controls including a reaction in the presence of 2 μM valinomycin to dissipate ΔΨ. Note that indicated amounts of Cyb2Δ-DHFR-His were added to 100-uL reactions and that all reactions contained MTX.

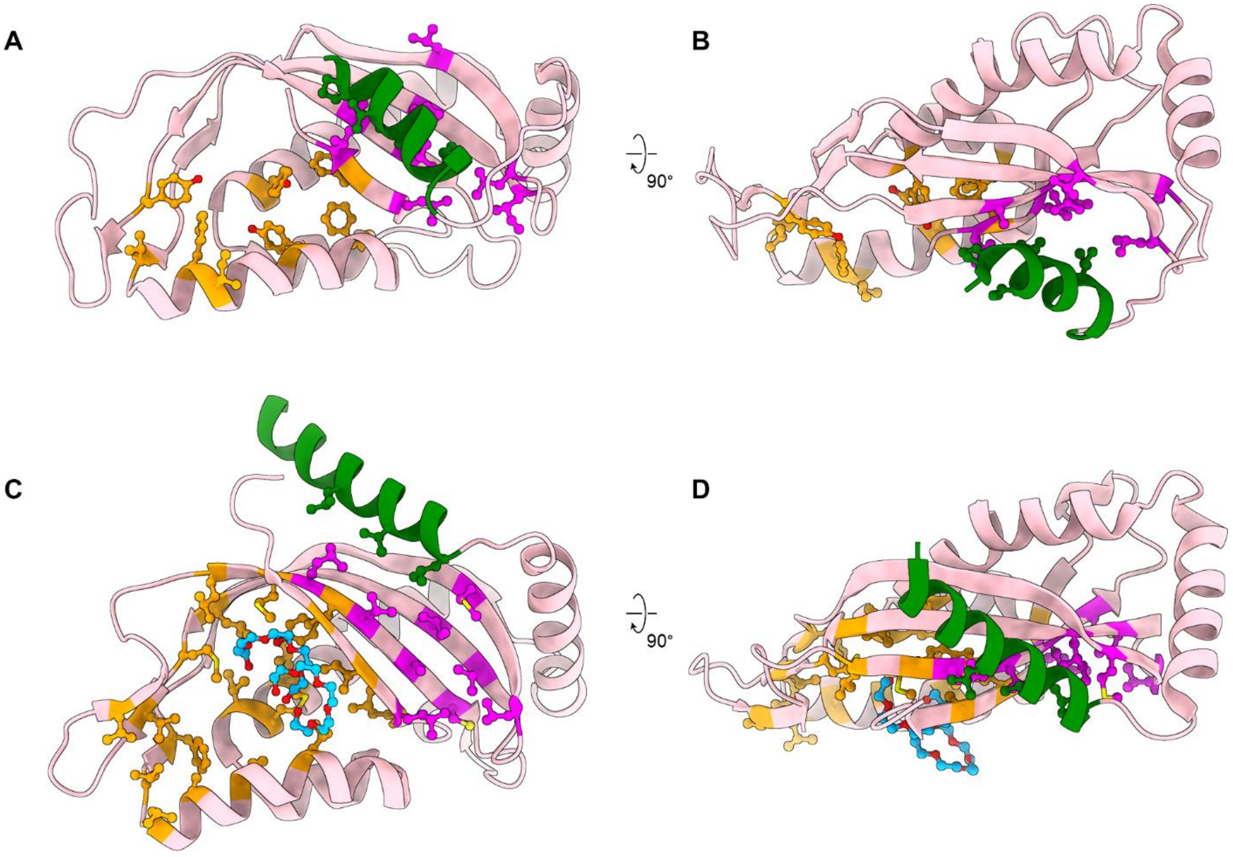

**Fig. S10. Hydrophobic clusters on the surface of Tim44-CTD.**
(**A** and **B**) Crystal structure of isolated yeast Tim44-CTD (PDB ID: 2FXT). The view angle in A is the same as in Fig. 5 A and B. Hydrophobic amino acids in Clusters 1 and 2 are shown in purple and orange, respectively. The green helix is the α6 segment (positions 247–260) immediately preceding the CTD that was included in the crystallization construct (hydrophobic side chains that interact with Cluster 1 are shown as sticks). (**C** and **D**) As in A and B, but showing the crystal structure of isolated human Tim44-CTD (PDB ID: 2CW9). The view angles are equivalent to A and B. The green helix is a segment of positions 270–287 (equivalent to α6). Note that in the human Tim44-CTD structure, Cluster 2 has a more open structure and binding of pentaethylene glycol molecules (blue) has been observed.

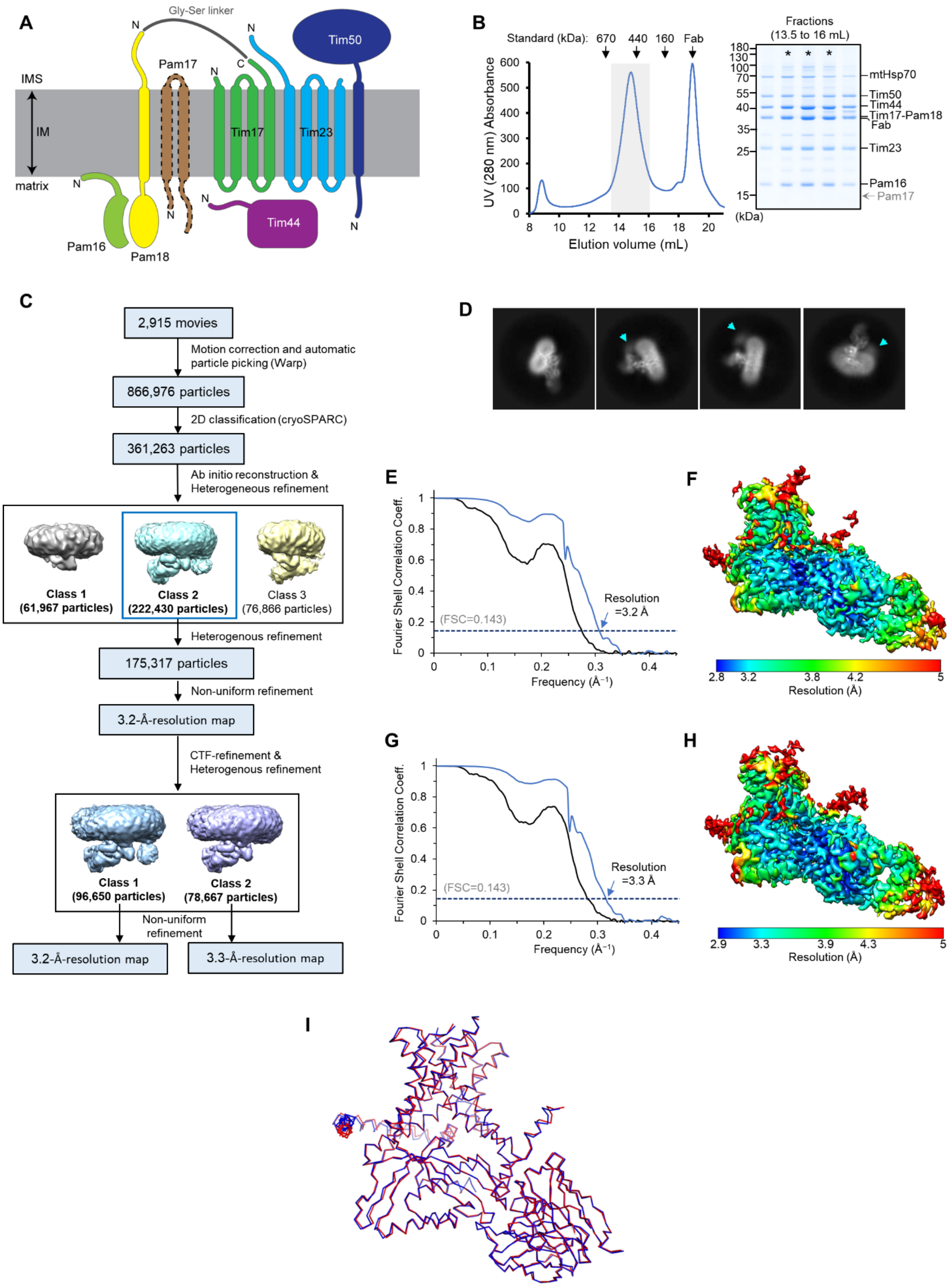

**Fig. S11. Cryo-EM analysis of the fusion TIM$^{MOTOR}$ construct.**

(**A**) Schematic diagram of the subunit topology of the fusion TIM$^{MOTOR}$ construct. The C-terminus of Tim17 was fused to the N-terminus of Pam18 via a Gly-Ser linker. (**B**) Purification of the fusion construct. After purification via Spot-tagged Tim23, the sample was subjected to Superose-6 size-exclusion chromatography (left). The gray box indicates fractions further analyzed by non-reducing SDS-PAGE and Coomassie staining (right). Fractions marked with asterisks are pooled and used for cryo-EM analysis. We note that we also co-expressed Pam17, but it hardly co-purified in accordance with a previous report (18). The expected band position for Pam17 is also indicated. (**C**) Summary of single particle analysis of the Fab-bound fusion TIM$^{MOTOR}$ complex. Note that Class 2 shows substantially weaker features of Pam16–Pam18 than Class 1. (**D**) Selected 2D class images. Cyan arrowhead, a putative feature of Pam16/18. (**E**–**F**) Fourier shell correlation (E) and local resolution distribution (F) for Class 1 of the fusion TIM$^{MOTOR}$ complex (see panel C). (**G**–**H**) As in E–F, but for the structure of Class 2. (**I**) Superposition between the structures of the Tim17/23/44+Pam16/18 (blue) complex, and the fusion TIM$^{MOTOR}$ construct (red). Cα RMSD is ~0.38 Å.

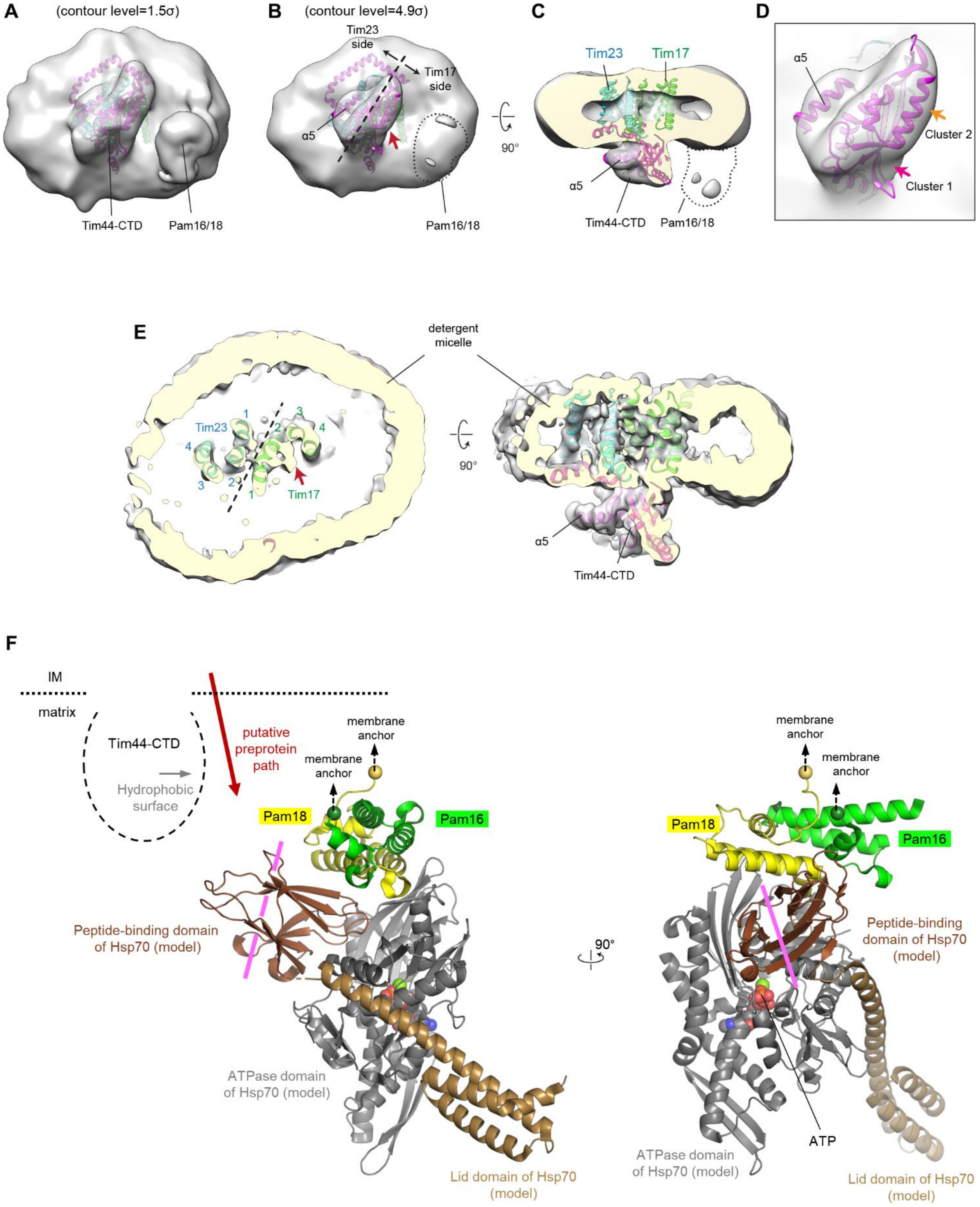

**Fig. S12. Orientation of Pam16/18 and mtHsp70 in the TIM complex.**
(**A**–**D**) Docking of the core TIM$^{MOTOR}$ model (purple, Tim44; green, Tim17; blue, Tim23) into the low-resolution cryo-EM reconstruction shown in Fig. S2H. Two different levels of the isosurface (semitransparent gray surface) are used to show the protruding features (panels B to D are the same contour level). In B, polypeptide exit site is indicated by a

red arrowhead. The dashed line indicates the plane of the Tim17-Tim23 interface. Panels C and D show a cutaway side view and a zoom-in view into the Tim44-CTD feature, respectively. In D, directions of hydrophobic patches (Clusters 1 and 2) are indicated. (**E**) As in Fig. 5 D and E, but showing the structure of Class 2 without the putative Pam16/18 globular domain (see Fig. S11C). (**F**) Modelling the position of mtHsp70. We placed the crystal structure of Pam16/18 (PDB ID: 2GUZ), based on the low-resolution cryo-EM feature, (positions of the membrane-matrix interface an Tim44-CTD are indicated by dashed lines). We oriented the Pam16–Pam18 model based on the positions of the N-termini (spheres) of the Pam16 and Pam18 construct that are linked to a membrane anchor in the full-length proteins. We then dock the model of the bacterial Hsp70 DnaK (PDB ID: 5NRO), which exhibits ~50% sequence identity to yeast mtHsp70 (Ssc1). This DnaK crystal structure contains a co-crystalized J protein (DnaJ), which allowed us to orient the Hsp70 model with respect to the J-protein Pam18. A substrate peptide is expected to bind to the pocket of the peptide-binding domain (dark brown) with the peptide arranged along the purple line. The putative preprotein exit direction from the Tim17 cavity is indicated with a red arrow.

### Table S1. List of yeast strains

| Name | Genotype/Description | Reference |
|---|---|---|
| BY4741 | *MAT***a** *his3-1, leu2-0, met15-0, ura3-0* | Horizon Discovery |
| yMLT62 | *MAT***a** *leu2-0*::*pACT1-GEV*::*HIS3, rps9Δ, mek1Δ, his3-1, met15-0, ura3-0* | A gift from J. Thorner |
| R1158 | BY4741 *URA*::$P_{CMV}$*-tTA* | Horizon Discovery |
| ySS078 | yMLT62 *TIM17-3C-Spot*::*HphMX* | This study |
| ySS025 | yMLT62 *ura3-0*::pSS011(Tim23/17-Spot/50/44/21)::*URA3* *HO*::pSS015(Pam16/17/18/Mgr2)::*LEU2* | This study |
| ySS027 | yMLT62 *ura3-0*::pSS013(Tim23/17/50/44/21-Spot)::*URA3* *HO*::pSS015(Pam16/17/18/Mgr2)::*LEU2* | This study |
| ySS047 | yMLT62 *ura3-0*::pSS077(Tim23/17-Spot/44[–GS])::*URA3* | This study |
| ySS055 | yMLT62 *ura3-0*::pSS077(Tim23/17-Spot/44[–GS])::*URA3* *HO*::pSS082(Pam16/18)::*LEU2* | This study |
| ySS107 | yMLT62 *ura3-0*::pSS107(Tim23-Spot/Tim17-15xGS-Pam18/Tim50/Tim44/Pam16)::*URA3* *HO*::pSS109(Pam17-His)::*LEU2* | This study |
| ySS121 | BY4741 *TIM50-HA*::*NatMX TIM23-Myc*::*HphMX* | This study |
| yYC17 | R1158 $P_{TIM17}$::*KanMX-tetO7*-$P_{CYC1}$ | This study |
| yYC23 | R1158 $P_{TIM23}$::*KanMX-tetO7*-$P_{CYC1}$ | This study |
| yYC02 | BY4741 *TIM17-HA*::*NatMX TIM23-Myc*::*HphMX* | This study |
| yYC03 | BY4741 *TIM17-HA*::*NatMX TIM23-Myc*::*HphMX HO*::*TIM17*(WT)-Spot::*LEU2* | This study |
| yYC04 | BY4741 *TIM17-HA*::*NatMX TIM23-Myc*::*HphMX HO*::*tim17*(D17N/E126Q)-*Spot*::*LEU2* | This study |
| yYC05 | BY4741 *TIM17-HA*::*NatMX TIM23-Myc*::*HphMX HO*::*tim17*(D76N/E126Q)-*Spot*::*LEU2* | This study |

### Table S2. List of plasmids

| Name | Description | Reference |
|---|---|---|
| pYTK001 to pYTK096 | Original MoClo YTK plasmids | (80) |
| pYTK-e106 | *HO* integration vector containing a *LEU2* marker | (65) |
| pYTK-e112 | CEN/ARS plasmid containing a *LEU2* marker | (65) |
| pYTK-e115 | CEN/ARS plasmid containing an *NatMX* marker. Assembled from pYTK084, pYTK008, pYTK047, pYTK073, pYTK078, and pYTK081. | This study |
| pYTK-e122 | 2μ plasmid containing a *LEU2* marker. Assembled from pYTK084, pYTK008, pYTK047, pYTK073, pYTK075, and pYTK082. | This study |
| pYTK-e201 | MoClo YTK part (type 4a) for 3C-2xSpot (Amino acid sequence: GSASGTLEVLFQGPTASGPDRVRA-VSHWSSGGGSGGGSTPDRVRAVSHWSS*; also see Table S4) | This study |
| pYTK-e203 | MoClo YTK part (type 4a) for 7xHis (Amino acid sequence: SGHHHHHHH*; also see Table S4) | This study |
| pYTK001-Tim23 | MoClo YTK part (type 3) for Tim23 (also see Table S4) | This study |
| pYTK001-Tim17 | MoClo YTK part (type 3) for Tim17 (also see Table S4) | This study |
| pYTK001-Tim50 | MoClo YTK part (type 3) for Tim50 (also see Table S4) | This study |
| pYTK001-Tim44 | MoClo YTK part (type 3) for Tim44 (also see Table S4) | This study |
| pYTK001-Tim21 | MoClo YTK part (type 3) for Tim21 (also see Table S4) | This study |
| pYTK001-Pam16 | MoClo YTK part (type 3) for Pam16 (also see Table S4) | This study |
| pYTK001-Pam17 | MoClo YTK part (type 3) for Pam17 (also see Table S4) | This study |
| pYTK001-Pam18 | MoClo YTK part (type 3) for Pam18 (also see Table S4) | This study |

| | | |
|---|---|---|
| pYTK001-Mgr2 | MoClo YTK part (type 3) for Mgr2 (also see Table S4) | This study |
| pYTK095-Tim23 | MoClo YTK expression cassette for *pGAL1*-Tim23. Assembled from pYTK095, pYTK002, pYTK030, pYTK001-Tim23, pYTK051, and pYTK067. | This study |
| pYTK095-Tim23-Spot | MoClo YTK expression cassette for *pGAL1*-Tim23-Spot. Assembled from pYTK095, pYTK002, pYTK030, pYTK001-Tim23, pYTK-e201, pYTK061, and pYTK067. | This study |
| pYTK095-Tim17 | MoClo YTK expression cassette for *pGAL1*-Tim17. Assembled from pYTK095, pYTK003, pYTK030, pYTK001-Tim17, pYTK051, and pYTK068. | This study |
| pYTK095-Tim17-Spot | MoClo YTK expression cassette for *pGAL1*-Tim17-Spot. Assembled from pYTK095, pYTK003, pYTK030, pYTK001-Tim17, pYTK-e201, pYTK061, and pYTK068. | This study |
| pYTK095-Tim17-15xGS-Pam18 | MoClo YTK expression cassette for *pGAL1*-Tim17-15xGS-Pam18 fusion protein (15xGS linker and Pam18 CDS were introduced into pYT095-Tim17 by PCR). | This study |
| pYTK095-Tim50 | MoClo YTK expression cassette for *pGAL1*-Tim50. Assembled from pYTK095, pYTK004, pYTK030, pYTK001-Tim50, pYTK051, and pYTK069. | This study |
| pYTK095-Tim44 | MoClo YTK expression cassette for *pGAL1*-Tim44. Assembled from pYTK095, pYTK005, pYTK030, pYTK001-Tim44, pYTK051, and pYTK070. | This study |
| pYTK095-Tim44 (–GS) | MoClo YTK expression cassette for *pGAL1*-Tim44 (contains a stop codon after the last amino acid; thus, a GlySer linker is absent at the C-terminus). | This study |
| pYTK095-Tim21 | MoClo YTK expression cassette for *pGAL1*-Tim21. Assembled from pYTK095, pYTK006, pYTK030, pYTK001-Tim21, pYTK051, and pYTK071. | This study |
| pYTK095-Tim21-Spot | MoClo YTK expression cassette for *pGAL1*-Tim21-Spot. Assembled from pYTK095, pYTK006, pYTK030, pYTK001-Tim21, pYTK-e201, pYTK061, and pYTK071. | This study |
| pYTK095-Pam16 | MoClo YTK expression cassette for *pGAL1*-Pam16. Assembled from pYTK095, pYTK002, pYTK030, pYTK001-Pam16, pYTK052, and pYTK067. | This study |
| pYTK095-Pam17 | MoClo YTK expression cassette for *pGAL1*-Pam17. Assembled from pYTK095, pYTK003, pYTK030, pYTK001-Pam17, pYTK052, and pYTK068. | This study |
| pYTK095-Pam17-7xHis | MoClo YTK expression cassette for *pGAL1*-Pam17-7xHis. Assembled from pYTK095, pYTK003, pYTK030, pYTK001-Pam17, pYTKe203, pYTK062, and pYTK068. | This study |
| pYTK095-Pam18 | MoClo YTK expression cassette for *pGAL1*-Pam18. Assembled from pYTK095, pYTK004, pYTK030, pYTK001-Pam18, pYTK052, and pYTK069. | This study |
| pYTK095-Mgr2 | MoClo YTK expression cassette for *pGAL1*-Mgr2. Assembled from pYTK095, pYTK005, pYTK030, pYTK001-Mgr2, pYTK052, and pYTK070. | This study |
| pYTK095 L2-RE | MoClo YTK filler cassette (ConL2-Spacer-ConRE). Assembled from pYTK095, pYTK004, pYTK048, and pYTK072. | This study |
| pYTK095 L3-RE | MoClo YTK filler cassette (ConL3-Spacer-ConRE). Assembled from pYTK095, pYTK005, pYTK048, and pYTK072. | This study |
| pYTK095 L4-RE | MoClo YTK filler cassette (ConL4-Spacer-ConRE). Assembled from pYTK095, pYTK006, pYTK048, and pYTK072. | This study |
| pYTK095 L5-RE | MoClo YTK filler cassette (ConL5-Spacer-ConRE). Assembled from pYTK095, pYTK007, pYTK048, and pYTK072. | This study |
| pYTK095 LS-R1 | MoClo YTK filler cassette (ConLS-Spacer-ConR1). Assembled from pYTK095, pYTK002, pYTK048, and pYTK067. | This study |
| pSS011 | pYTK096 (Tim23, Tim17-Spot, Tim50, Tim44, Tim21). Assembled from pYTK096, pYTK095-Tim23, pYTK095-Tim17-Spot, pYTK095-Tim50, pYTK095-Tim44, pYTK095-Tim21, and pYTK095-L5-RE. | This study |

| | | |
|---|---|---|
| pSS013 | pYTK096 (Tim23, Tim17, Tim50, Tim44, Tim21-Spot). Assembled from pYTK096, pYTK095-Tim23, pYTK095-Tim17, pYTK095-Tim50, pYTK095-Tim44, pYTK095-Tim21-Spot, and pYTK095 L5-RE. | This study |
| pSS015 | pYTK-e106 (Pam16, Pam17, Pam18, Mgr2). Assembled from pYTK-e106, pYTK095-Pam16, pYTK095-Pam17, pYTK095-Pam18, pYTK095-Mgr2, and pYTK095 L4-RE. | This study |
| pSS077 | pYTK096 (Tim23, Tim17-Spot, Tim44 (–GS)). Assembled from pYTK096, pYTK095-Tim23, pYTK095-Tim17-Spot, pYTK095 L2-R3 (spacer), pYTK095-Tim44 (–GS), and pYTK095 L4-RE. | This study |
| pSS082 | pYTK-e106 (Pam16, Pam18). Assembled from pYTK-e106, pYTK095-Pam16, pYTK095 L1-R2 (spacer), pYTK095-Pam18, and pYTK095 L3-RE. | This study |
| pSS107 | pYTK096 (Tim23-Spot, Tim17-15xGS-Pam18, Tim50, Tim44, Pam16). Assembled from pYTK096, pYTK095-Tim23-Spot, pYTK095-Tim17-15xGS-Pam18, pYTK095-Tim50, pYTK095-Tim44 (–GS), pYTK095-Pam16, and pYTK095 L5-RE. | This study |
| pSS109 | pYTK-e106 (Pam17-His). Assembled from pYTK-e106, pYTK095 LS-R1, pYTK095-Pam17-His, and pYTK095 L2-RE. | This study |
| pSS122 | pYTK-e112 *pTIM17*-Tim17-Spot (also see Table S4) | This study |
| pSS140 | pYTK-e112 *pTIM17*-Tim17 (also see Table S4) | This study |
| pSS070 | pETDuet-1 6xHis-3C-Tim44-CTD (residues 210–431) (also see Table S4) | This study |
| pYC17a | pYTK-e115 *pTIM17*-Tim17-HA (also see Table S4) | This study |
| pYC17b | pYTK-e122 *pDDI2*-Tim17-HA (also see Table S4) | This study |
| pYC23 | pYTK-e115 *pTIM23*-Tim23-HA (also see Table S4) | This study |
| pYC002 | pET32a Cyb2Δ-DHFR (also see Table S4) | This study |

### Table S3. List of primers

| Name | Sequence (including notes) |
|---|---|
| EP_442 | GATGCACTAAGAGGCAAACATGAC<br>(To confirm chromosomal integration at *TIM23* locus) |
| EP_443 | GGACGGCTCTGACAGTTTCG<br>(To confirm chromosomal integration at *TIM23* locus) |
| EP_446 | GTTGGAGGCATACAAGGAACAG<br>(To confirm chromosomal integration at *TIM17* locus) |
| EP_447 | GCACTAGCTTTTGGCTTGTTG<br>(To confirm chromosomal integration at *TIM17* locus) |
| EP_450 | AGAATACAGCAGGAGCAAATGG<br>(To confirm chromosomal integration at *TIM50* locus) |
| EP_451 | GCATCAGATCATTAGGTGTGTCTACATC<br>(To confirm chromosomal integration at *TIM50* locus) |
| SS_1696 | cctcttctactctctttgcctgtacatactacacgttatagcgttaacaaaagcagatagGGCGTTAGTATCGAATCG<br>(To replace the endogenous *TIM23* promoter with the TRE promoter. Uppercase for sequence specific to TRE-kanMX and lower case for sequences homologous to yeast chromosomal sequences directly before the starting codon of Tim23) |
| SS_1697 | ttggccgcccacggcagcattcgcatcatcggtaggtgtcttatctccaaaaagccacgacatGGATCCCCCGAATTG<br>(To replace the endogenous *TIM23* promoter with the TRE promoter. Uppercase for sequence specific to TRE-kanMX and lower case for sequences homologous to the N-terminus of Tim23) |
| SS_1698 | GGTTATTGCATTCGCC<br>(To confirm chromosomal integration of the TRE cassette into the *TIM23* locus) |
| SS_1699 | CAGGACCTGATATTATGTTATTG<br>(To confirm chromosomal integration of the TRE cassette into the *TIM23* locus) |
| SS_1700 | ccaaatccagaggataaaagcactattctcatcaaaagatggaaaagctgtgaaagagtcGGCGTTAGTATCGAATCG |

| | |
|---|---|
| | (To replace the endogenous *TIM17* promoter with the TRE promoter. Uppercase for sequence specific to TRE-kanMX and lower case for sequences homologous to yeast chromosomal sequences directly before the starting codon of Tim17) |
| SS_1701 | agcaccaccgaaatcatttagtatgactataggacatggatctctcgaatgatcggctgacatGGATCCCCCGAATTG<br>(To replace the endogenous *TIM17* promoter with the TRE promoter. Uppercase for sequence specific to TRE-kanMX and lower case for sequences homologous to the N-terminus of Tim17) |
| SS_1702 | ACTGCTATTGTTCAACAAAG<br>(To confirm chromosomal integration of the TRE cassette into the *TIM17* locus) |
| SS_1703 | GCTCACCTAATGGCG<br>(To confirm chromosomal integration of the TRE cassette into the *TIM17* locus) |
| YC_1760 | gaaagaattcCTCCAGCATTATAAAGC<br>(To amplify a DNA segment expressing Tim17 with a 5' *Eco*RI site) |
| YC_1761 | gaaaggatccAGTTTCTGCACTAGC<br>(To amplify the expression cassette of Tim17 with a 3' *Bam*HI site) |
| YC_1762 | gaaagaattcATTGAAAAAAAGAGAAAATACTG<br>(To amplify a DNA segment expressing Tim23 with a 5' *Eco*RI site) |
| YC_1763 | gaaaggatccGCCATCGAAAACAATAG<br>(To amplify a DNA segment expressing Tim23 with a 3' *Bam*HI site) |
| YC_1981 | gaaagaattcCAGCCCACATACTAC<br>(To amplify a DNA segment of the *DDI2* promoter with a 5' *Eco*RI site) |
| YC_1982 | gaaaggtaccGATTGATTCTTTTGAAGAGAAG<br>(To amplify a DNA segment of the *DDI2* promoter with a 3' *KpnI* site) |

### Table S4. DNA sequences

(Note, sequences are inserts only; the plasmid backbones are not included.)

| Name | DNA sequence | Reference |
|---|---|---|
| pYTK-e201 | GGTCTCaatccgctagtggtaccctggaggtgttatttcagggcccgactgctagcggccctgatagagttagagcagtctcacattggtcttctggtggaggttctggcggtggttcaactccagatagagtacgtgctgtttctcactggagttcttaatggctGAGACC<br>(Uppercase, *Bsa*I sites) | This study |
| pYTK-e203 | GGTCTCaatccggtcaccatcaccaccatcatcactaatggctGAGACC<br>(Uppercase, *Bsa*I sites) | This study |
| TIM17-3C-Spot (PCR product to introduce a Spot-tag to chromosomal *TIM17*) | GATATGCTGCTTGGCAAGCCAAACCTATGGCTCCTCCTTTGCCCGAAGCACCTTCCTCTCAACCTCTGCAAGCTgctagtggtaccctggaggtgttatttcagggcccgactgctagcggccctgatagagttagagcagtctcacattggtcttctggtggaggttctggcggtggttcaactccagatagagtacgtgctgtttctcactggagttcttgataaggcgcgccacttctaaataagcgaatttcttatgatttatgatttttattattaaataagttataaaaaaaataagtgtatacaaattttaaagtgactcttaggttttaaaacgaaaattcttattcttgagtaactctttcctgtaggtcaggttgctttctcaggtatagtatgaggtcgctcttattgaccacacctctaccggcagatccgctagggataacagggtaatatagatctgtttagcttgcctcgtccccgccgggtcacccggccagcgacatggaggcccagaataccctccttgacagtcttgacgtgcgcagctcaggggcatgatgtgactgtcgcccgtacatttagcccatacatccccatgtataatcatttgcatccatacattttgatggccgcacggcgcgaagcaaaaattacggctcctcgctgcggacctgcgagcagggaaacgctcccctcacagacgcgttgaattgtccccacgccgcgcccctgtagagaaatataaaaggttaggatttgccactgaggttcttctttcatatacttccttttaaaatcttgctaggatacagttctcacatcacatccgaacataaacaacc**atg**ggtaaaaagcctgaactcaccgcgacgtctgtcgagaagtttctgatcgaaaagttcgacagcgtctccgacctgatgcagctctcggagggcgaagaatctcgtgctttcagcttcgatgtaggagggcgtggatatgtcctgcgggtaaatagctgcgccgatggtttctacaaagatcgttatgtttatcggcactttgcatcggccgcgctcccgattccggaagtgcttgacattggggaattcagcgagagcctgacctattgcatctcccgccgtgcacagggtgtcacgttgcaagacctgcctgaaaccgaactgcccgctgttctgcagccggtcgcggaggccatggatgcgatcgctgcggccgatcttagccagacgagcgggttcggcccattcggaccgcaaggaatcggtcaatacactacatggcgtgatttcatatgcgcgattgctgatccccatgtgtatcactggcaaactgtgatggacgacaccgtcagtgcgtccgtcgcgcaggctctcgatgagctgatgctttgggccgaggactgccccgaagtccggcacctcgtgcacgcggatttcggctccaacaatgtcctgacggacaatggccgcataacagcggtcattgactggagcgaggcgatgttcggggattcccaatacgaggtcgccaacatcttcttctggaggccgtggttggcttgtatggagcagcagacgcgctacttcgagcggaggcatccggagcttgcaggatcgccgcggctccgggcgtatatgctccgcattggtcttgaccaactctatcagagcttggttgacggcaatttcgatgatgcagcttgggcgcagggtcgatgcgacgcaatcgtccgatccggagccgggactgtcgggcgtacacaaatcgcccgcagaagcgcggccgtctggaccgatggctgtgtagaagtactcgccgatagtggaaaccgacgccccagcactcgtccgagggcaaaggaa**taa**tcagtactgacaataaaaagattcttgttttcaagaacttgtcatttgtatagtttttttatattgtagttgttctattttaatcaaatgttagc | This study |

| | | |
|---|---|---|
| | gtgatttatattttttttcgcctcgacatcatctgcccagatgcgaagttaagtgcgcagaaagtaatatcatgcgtcaatcgtatgt<br>gaatgctggtcgctatactgctgtcgattcgatactaacgccgccatccagtgtcgaaaacgagctccaattcatcgatgatat<br>cagatccactagtggcctatgcggGACGTATCGCACTAGCCTCCTTTACGTTTTTACTTTATTTC<br>AGCCTTTTATTTCAAGATTACCAACCATTTCTC<br>(Uppercase, homology arms; bold/underlined, start and stop codons of hygromycin B phosphotransferase) | |
| TIM17-HA<br>(PCR product to introduce an HA-tag to chromosomal *TIM17*) | GATATGCTGCTTGGCAAGCCAAACCTATGGCTCCTCCTTTGCCCGAAGCACCTTCCT<br>CTCAACCTCTGCAAGCTgctggaggggctaccacggctagtggtaccggcgaaaatttatactttcaaggtactg<br>ctagcgggggcggttcttatccctatgacgtcccggactatgcaggatcctatccatatgacgttccagattacgcttaaggcg<br>cgccacttctaaataagcgaatttcttatgatttatgatttttattattaaataagttataaaaaaaataagtgtatacaaattttaaa<br>gtgactcttaggttttaaaacgaaaattcttattcttgagtaactctttcctgtaggtcaggttgctttctcaggtatagtatgaggtcg<br>ctcttattgaccacacctctaccggcagatccgctagggataacagggtaatatagatctgtttagcttgcctcgtccccgccgg<br>gtcacccggccagcgacatggaggcccagaataccctccttgacagtcttgacgtgcgcagctcaggggcatgatgtgact<br>gtcgcccgtacatttagcccatacatccccatgtataatcatttgcatccatacattttgatggccgcacggcgcgaagcaaaa<br>attacggctcctcgctgcagacctgcgagcagggaaacgctcccctcacagacgcgttgaattgtccccacgccgcgcccc<br>tgtagagaaatataaaaggttaggatttgccactgaggttcttctttcatatacttccttttaaaatcttgctaggatacagttctcac<br>atcacatccgaacataaacaacc**<u>atg</u>**ggtaccactcttgacgacacggcttaccggtaccgcaccagtgtcccggggac<br>gccgaggccatcgaggcactggatgggtccttcaccaccgacaccgtcttccgcgtcaccgccaccggggacggcttcac<br>cctgcgggaggtgccggtggacccgcccctgaccaaggtgttccccgacgacgaatcggacgacgaatcggacgacgg<br>ggaggacggcgacccggactcccggacgttcgtcgcgtacggggacgacggcgacctggcgggcttcgtggtcgtctcgt<br>actccggctggaaccgccggctgaccgtcgaggacatcgaggtcgccccggagcaccgggggcacggggtcgggcgc<br>gcgttgatggggctcgcgacggagttcgcccgcgagcggggcgccgggcacctctggctggaggtcaccaacgtcaacg<br>caccggcgatccacgcgtaccggcggatggggttcaccctctgcggcctggacaccgccctgtacgacggcaccgcctcg<br>gacggcgagcaggcgctctacatgagcatgccctgcccc**<u>taa</u>**tcagtactgacaataaaaagattcttgttttcaagaacttg<br>tcatttgtatagtttttttatattgtagttgttctattttaatcaaatgttagcgtgatttatattttttttcgcctcgacatcatctgcccagat<br>gcgaagttaagtgcgcagaaagtaatatcatgcgtcaatcgtatgtgaatgctggtcgctatactgctgtcgattcgatactaac<br>gccgccatccagtgtcgaaaacgagctcgaattcatcgatgatatcagatccactagtggcctatgcggccGACGTATC<br>GCACTAGCCTCCTTTACGTTTTTACTTTATTTCAGCCTTTTATTTCAAGATTACCAACC<br>ATTTCTC<br>(Uppercase, homology arms; bold/underlined, start and stop codons of nourseothricin N-acetyl transferase) | This study |
| TIM23-myc<br>(PCR product to introduce a myc-tag to chromosomal *TIM23*) | GGTTTGAAACCCATGGGTTATTCCTCGGCAATGGTGGCCGCTGCGTGCGCCGTCTG<br>GTGTAGTGTCAAGAAAAGACTACTTGAAAAAgctagtggtaccctggaggtgttatttcagggcccgact<br>gctagcggcgaacaaaagttgatttctgaagaagatttgaacggtgaacaaaagctaatctccgaggaagacttgtgataa<br>ggcgcgccacttctaaataagcgaatttcttatgatttatgatttttattattaaataagttataaaaaaaataagtgtatacaaattt<br>taaagtgactcttaggttttaaaacgaaaattcttattcttgagtaactctttcctgtaggtcaggttgctttctcaggtatagtatgag<br>gtcgctcttattgaccacacctctaccggcagatccgctagggataacagggtaatatagatctgtttagcttgcctcgtccccg<br>ccgggtcacccggccagcgacatggaggcccagaataccctccttgacagtcttgacgtgcgcagctcaggggcatgatgt<br>gactgtcgcccgtacatttagcccatacatccccatgtataatcatttgcatccatacattttgatggccgcacggcgcgaagca<br>aaaattacggctcctcgctgcggacctgcgagcagggaaacgctcccctcacagacgcgttgaattgtccccacgccgcg<br>cccctgtagagaaatataaaaggttaggatttgccactgaggttcttctttcatatacttccttttaaaatcttgctaggatacagttc<br>tcacatcacatccgaacataaacaacc**<u>atg</u>**ggtaaaaagcctgaactcaccgcgacgtctgtcgagaagtttctgatcgaa<br>aagttcgacagcgtctccgacctgatgcagctctcggagggcgaagaatctcgtgctttcagcttcgatgtaggagggcgtg<br>gatatgtcctgcgggtaaatagctgcgccgatggtttctacaaagatcgttatgtttatcggcactttgcatcggccgcgctcccg<br>attccggaagtgcttgacattggggaattcagcgagagcctgacctattgcatctcccgccgtgcacagggtgtcacgttgca<br>agacctgcctgaaaccgaactgcccgctgttctgcagccggtcgcggaggccatggatgcgatcgctgcggccgatcttag<br>ccagacgagcgggttcggcccattcggaccgcaaggaatcggtcaatacactacatggcgtgatttcatatgcgcgattgct<br>gatccccatgtgtatcactggcaaactgtgatggacgacaccgtcagtgcgtccgtcgcgcaggctctcgatgagctgatgct<br>ttgggccgaggactgccccgaagtccggcacctcgtgcacgcggatttcggctccaacaatgtcctgacggacaatggccg<br>cataacagcggtcattgactggagcgaggcgatgttcggggattcccaatacgaggtcgccaacatcttcttctggaggccgt<br>ggttggcttgtatggagcagcagacgcgctacttcgagcggaggcatccggagcttgcaggatcgccgcggctccgggcgt<br>atatgctccgcattggtcttgaccaactctatcagagcttggttgacggcaatttcgatgatgcagcttgggcgcagggtcgatg<br>cgacgcaatcgtccgatccggagccgggactgtcgggcgtacacaaatcgcccgcagaagcgcggccgtctggaccgat<br>ggctgtgtagaagtactcgccgatagtggaaaccgacgccccagcactcgtccgagggcaaaggaa**<u>taa</u>**tcagtactga<br>caataaaaagattcttgttttcaagaacttgtcatttgtatagtttttttatattgtagttgttctattttaatcaaatgttagcgtgatttata<br>ttttttttcgcctcgacatcatctgcccagatgcgaagttaagtgcgcagaaagtaatatcatgcgtcaatcgtatgtgaatgctg<br>gtcgctatactgctgtcgattcgatactaacgccgccatccagtgtcgaaaacgagctccaattcatcgatgatatcagatcca<br>ctagtggcctatgcggACTGATGGCGCTTGTATATAGCATTTGAAAAATAATAGTACGTAACG<br>CAGAAAACAAACCAATGAA<br>(Uppercase, homology arms; bold/underlined, start and stop codons of hygromycin B phosphotransferase) | This study |

| | | |
|---|---|---|
| TIM50-HA<br>(PCR product to introduce an HA-tag to chromosomal *TIM50*) | TATTTGAAGAGGAAAAGAAAAAGAAGAAGATTGCTGAATCCAAAgctggaggggctaccacg<br>gctagtggtaccggcgaaaatttatactttcaaggtactgctagcgggggcggttcttatccctatgacgtcccggactatgca<br>ggatcctatccatatgacgttccagattacgcttaaggcgcgccacttctaaataagcgaatttcttatgatttatgatttttattatta<br>aataagttataaaaaaaataagtgtatacaaattttaaagtgactcttaggttttaaaacgaaaattcttattcttgagtaactcttt<br>cctgtaggtcaggttgctttctcaggtatagtatgaggtcgctcttattgaccacacctctaccggcagatccgctagggataac<br>agggtaatatagatctgtttagcttgcctcgtccccgccgggtcacccggccagcgacatggaggcccagaataccctccttg<br>acagtcttgacgtgcgcagctcaggggcatgatgtgactgtcgcccgtacatttagcccatacatccccatgtataatcatttgc<br>atccatacattttgatggccgcacggcgcgaagcaaaaattacggctcctcgctgcagacctgcgagcagggaaacgctc<br>ccctcacagacgcgttgaattgtccccacgccgcgcccctgtagagaaatataaaaggttaggatttgccactgaggttcttct<br>ttcatatacttccttttaaaatcttgctaggatacagttctcacatcacatccgaacataaacaacc**<u>atg</u>**ggtaccactcttgacg<br>acacggcttaccggtaccgcaccagtgtcccgggggacgccgaggccatcgaggcactggatgggtccttcaccaccga<br>caccgtcttccgcgtcaccgccaccggggacggcttcaccctgcgggaggtgccggtggacccgcccctgaccaaggtgtt<br>ccccgacgacgaatcggacgacgaatcggacgacggggaggacggcgacccggactcccggacgttcgtcgcgtacg<br>gggacgacggcgacctggcgggcttcgtggtcgtctcgtactccggctggaaccgccggctgaccgtcgaggacatcgag<br>gtcgccccggagcaccgggggcacggggtcgggcgcgcgttgatggggctcgcgacggagttcgcccgcgagcgggg<br>cgccgggcacctctggctggaggtcaccaacgtcaacgcaccggcgatccacgcgtaccggcggatggggttcaccctct<br>gcggcctggacaccgccctgtacgacggcaccgcctcggacggcgagcaggcgctctacatgagcatgccctgcccc**<u>ta</u>**<br>**<u>a</u>**tcagtactgacaataaaaagattcttgttttcaagaacttgtcatttgtatagtttttttatattgtagttgttctattttaatcaaatgtta<br>gcgtgatttatattttttttcgcctcgacatcatctgcccagatgcgaagttaagtgcgcagaaagtaatatcatgcgtcaatcgta<br>tgtgaatgctggtcgctatactgctgtcgattcgatactaacgccgccatccagtgtcgaaaacgagctcgaattcatcgatgat<br>atcagatccactagtggcctatgcggccCACACCCTCATTTTGTTACTGTCATGTGAATAAAACTTA<br>TGTAT<br>(Uppercase, homology arms; bold/underlined, start and stop codons of nourseothricin N-acetyl transferase) | This study |
| pYTK001-Tim23 | GGTCTCatatgtcgtggctttttggagataagacacctaccgatgatgcgaatgctgccgtgggcggccaagatacaacc<br>aagcctaaggaactatcgttgaagcagagtttaggtttcgagccaaacatcaataacataatatcaggtcctggtggaatgc<br>atgtcgacaccgctaggctgcatcctttggctggtctagacaagggtgtggagtatttagatctggaagaagaacaactatcct<br>cgttagaaggctcacagggtctgatcccttcccgtgggtggaccgatgacctatgttacggtaccggtgccgtctacctgctgg<br>gacttggtatcggagggttttctggtatgatgcagggtctacagaatattccgcccaatagtcccggaaaattgcaattgaaca<br>ccgtcctgaatcacattactaagagaggtcccttcttaggtaataatgcggggattctcgcgttgagctacaatatcatcaattct<br>acaatagatgcactaagaggcaaacatgacaccgcgggctccattggcgctggggccctcacgggcgctttgttcaagtctt<br>caaaaggtttgaaacccatgggttattcctcggcaatggtggccgctgcgtgcgccgtctggtgtagtgtcaagaaaagacta<br>cttgaaaaaggatcctGAGACC<br>(Uppercase, *Bsa*I sites) | This study |
| pYTK001-Tim17 | GGTCTCatatgtcagccgatcattcgagagatccatgtcctatagtcatactaaatgatttcggtggtgcttttgccatgggtg<br>ccattggtggtgttgtttggcatgggattaaaggttttagaaattcgccattaggtgagcgtggttcaggagctatgagcgccatt<br>aaagcgcgtgctcccgtactgggtggtaattttggtgtgtggggtggtttattttcgacttttgattgcgctgtgaaggccgttagaa<br>agagagaggacccatggaatgctatcattgcagggtttttcactggtggcgctttagctgtaagaggtggttggaggcataca<br>aggaacagttcgatcacgtgtgcttgtttgttgggtgtgattgaaggtgtgggactaatgtttcaaagatatgctgcttggcaagc<br>caaacctatggctcctcctttgcccgaagcaccttcctctcaacctctgcaagctggatcctGAGACC<br>(Uppercase, *Bsa*I sites) | This study |
| pYTK001-Tim50 | GGTCTCatatgctgtccattttaagaaattcggtgagactaaattcaagggctcttagggttgtgccatctgccgccaacac<br>cttaacatcggttcaagcatccagaagacttttaacaagttattcaagttttttacaaaaagaaacaaaagacgacaagccta<br>aatccattcttacagatgatatgctgttcaaggccggtgttgacgtcgatgagaagggccaaggcaaaaacgaggaaacat<br>caggagaaggaggagaggacaagaatgaaccttcttccaaaagtgaaaaatctagaagaaaaagacaaacttctacag<br>atattaaaagagaaaagtatgctaactggttttacatttttttcgttgtctgcgttgacaggtactgcaatctacatggcaagggatt<br>gggagcctcaagagtctgaagaattgaagaaagacatcgataatggctacactttatcacttatgtataaaagattcaaggc<br>caggttcaactcaatgttcacctacttccaagagccacctttccctgatttactacctccaccaccaccaccaccgtaccaaag<br>gccattaactcttgttatcacattggaagatttttttggttcattctgagtggtcacaaaagcatggttggagaacggccaaaaga<br>cctggtgctgactacttcttgggttacctatcgcagtattacgaaattgttttgttttcatccaactatatgatgtactctgacaaaatc<br>gctgaaaaattagatccaatccatgcattcgtatcttataatttgttcaaggaacactgtgtttacaaagacggtgtgcacattaa<br>ggatctgtcaaaattgaatagagatttgagtaaagtaatcattattgacactgaccctaacagttacaaattgcaacctgaaaa<br>tgctattccaatggagccatggaatggtgaagctgatgacaaattagtaagattgattccatttttggagtaccttgccactcaac<br>aaaccaaggatgttagaccaatcttgaacagctttgaagacaagaagaacctagcagaagaatttgatcatcgtgtgaaaa<br>aattgaaggataaattttacggagatcataaatctggtggcaactgggcaatgacggcactaggtctaggaaattccctggg<br>cggcagcaccaagttcccgctcgatttgattcatgaagaaggacaaaagaactatttaatgttcatgaagatgattgaggaa<br>gaaaaggaaaaaattagaatacagcaggagcaaatgggcgggcaaacatttacgctgaaagactatgttgaaggtaact<br>tgccttcgccagaagaacaaatgaaaatacaattggagaagcagaaggaggtagacgccttatttgaagaggaaaagaa<br>aaagaagaagattgctgaatccaaaggatcctGAGACC<br>(Uppercase, *Bsa*I sites) | This study |

| | | |
|---|---|---|
| pYTK001-Tim44 | GGTCTCatatgcacagatccacttttatcaggacgtccggcacgagctctaggacactaaccgcaaggtacagatcgc<br>agtacacaggattactcgttgcccgagtattattctccacctctacgacccgtgcgcaaggtggaaaccctcgatcaccactc<br>cagattttccgcgatacattcaagaaggaatgggagaagtctcaggaactacaggagaacataaagacgctgcaagatg<br>cttcgggaaagttaggcgagtctgaggcctacaaaaaggctagggaagcatatttgaaagctcagagaggctccacaatt<br>gtgggtaaaactctgaaaaagaccggtgaaaccatggaacatatagccactaaggcctgggagtccgaactcggtaaga<br>acacaagaaaggcggctgccgccacggcgaagaagctggatgagagttttgagccagtgagacagacgaagatttaca<br>aggaagtctcagaagtcattgacgatggggagagttcccgatacggtgggtttatcacgaaagagcaaaggagacttaaa<br>cgtgagagagatctggcctctgggaaaagacacagggcagtaaagagcaatgaagatgcaggaacagcagtggttgcg<br>acaaatatcgagtctaaagaatcgtttggtaagaaagtggaggatttcaaggagaaaaccgtggttggccgttctatacaatc<br>tttaaagaacaaattgtgggatgaaagtgaaaaccccttaattgttgtcatgaggaaaataaccaacaaagtgggcggtttctt<br>tgcagaaacagaatcctcccgtgtttacagtcaatttaagctaatggacccaaccttttcgaacgaaagcttcaccagacactt<br>aagagaatacattgttcccgagattctcgaagcgtatgtgaaaggcgatgtcaaagttctcaaaaaatggttcagcgaggcg<br>ccattcaatgtttacgccgcccaacagaaaatcttcaaagaacaggatgtttacgccgatggccgtatcctagatatcagggg<br>cgttgaaatagtgagtgcaaagttattggctccccaagacatcccagtcttggtggtcgggtgtagagcacaagaaatcaac<br>ctttacaggaaaaagaaaactggcgagattgcggctggtgacgaagctaatatcttgatgagctcttatgccatggttttcacc<br>agagatccagagcaaatcgacgatgacgaaacagaagggtggaagatcttggagtttgtgcgcgggggttctagacaatt<br>caccggatcctGAGACC<br>(Uppercase, *Bsa*I sites) | This study |
| pYTK001-Tim21 | GGTCTCatatgagctcaagtttgcctaggtctttactccgtttaggacatagaaagcctttgtttccacgatataatactttcgt<br>gaactcatcggtaataactcatacgtcactgcttagaacaagattatatagtaatggcaccggcgcaacatcaggaaagaa<br>ggatgacaagactaggaataaacctaagccattatggcctcaagtaaaatccgcttctacgttcaccttttctggcatacttgtg<br>ataggagctgttggtatatctgctattgttatttacctaattctttcagaactattttcgccttcaggtgatacacagcttttcaacaga<br>gcagtttctatggtagagaaaaacaaagatataagaagtttgttacagtgcgacgatggcattacgggaaaagaaagattg<br>aaagcgtacggtgagcttataacgaatgacaaatggacaagaaacaggcctatagtatctaccaaaaaattagataaaga<br>aggtaggacacatcactatatgagattccacgttgaatccaaaaagaaaatagcgttggttcacttagaggctaaagaatcc<br>aaacagaattatcaacctgactttatcaacatgtacgttgatgttcccggagagaagcgttactatttgatcaagccaaaattgc<br>atccggtttctaattcgaagggctttctgggaattagatggggccccagaaaagatggatcctGAGACC<br>(Uppercase, *Bsa*I sites) | This study |
| pYTK001-Pam16 | GGTCTCatatggctcacagggctttcatacaggttataatcacaggaactcaagtttttggaaaagcattcgccgaggcgt<br>atagacaagcggcttcacaatcagtgaaacaaggtgctacaaatgcatcaagaaggggaacaggaaaaggcgaatatg<br>gtggtattacgttggatgagagttgtaaaattttaaatattgaagaatccaagggcgatttaaacatggacaagattaataaca<br>ggtttaactatctatttgaggttaacgataaagaaaaaggtggaagcttctacttacagagcaaagtttatcgagcagcagaa<br>aggttaaaatgggaactggctcagagagaaaaaaatgcgaaggcgaaagcaggggacgcttcaacagcgaaacctcc<br>tccgaattcaacaaattcatctggtgcagataatagtgcaagcagcaatcagggatcctGAGACC<br>(Uppercase, *Bsa*I sites) | This study |
| pYTK001-Pam17 | GGTCTCatatgtttaccagtgccattagattgtcatcgcaaagactgttcgctagtcaaccttctgtcaccgctgcggcattg<br>cgctcgactgctacaaccttacccttaagatcatattctcagcccgcatcccttcaagactccagtatcttgacatggtctgattttt<br>tcaaattgaggaaacagcagcgtagaatcaatgttggttcttcgctgtttactgctcttttgggctgtaacgtttcatgggcttacctt<br>tccacaatggaaatagacccgactcaaatgctattcggattcgacccattaactgtaatttcagctgggataatagcctctggt<br>gcactaggctacttgttgggtccgatagttggttcgcaagttttcaaactttcccataaccaacaattggcacagttcaacaaca<br>aaaataaagagtttctaaaacatatcatcaataacagggtcgatgcctcttctcaaagtttcagtaatcctgttccagattattac<br>ggtgaaaagataggttccttaaaggaatataagcaatggttaagagattgtcacgcttacgcaaagaaagccaaagaatttt<br>tgggatcctGAGACC<br>(Uppercase, *Bsa*I sites) | This study |
| pYTK001-Pam18 | GGTCTCatatgagttctcaaagtaatactggtaattctattgaggcaccacaactacccattcctggtcaaactaatggctct<br>gcgaacgttactgttgatggagctggtgttaatgtcggtatccagaatggttcgcagggtcaaaagaccggaatggacctttat<br>tttgatcaagctttgaactacatgggagaacatcctgtgataacaggttttggggcctttttaactttatattttacagccggtgcat<br>ataaatcaatatcgaagggacttaacggtggaaaatccactactgccttcttgaaaggcggatttgacccgaaaatgaattct<br>aaagaggctctacagattttgaatttgacagaaaatacattgactaaaaaaaagttgaaagaggttcataggaaaattatgtt<br>agctaatcatcctgacaaaggtggttctccatttttggccactaagataaacgaagctaaggactttttggaaaaaaggggtat<br>tagcaaaggatcctGAGACC<br>(Uppercase, *Bsa*I sites) | This study |
| pYTK001-Mgr2 | GGTCTCatatgcctcctcttccacaaaattatgcgcaacagcagccttcgaattgggacaaattcaaaatggggttgatga<br>tgggtactaccgtcggtgtctgcacaggcatcctatttggtggatttgccatcgcaactcaaggcccaggtcctgatggtgtagtt<br>agaacactagggaaatacattgctggttcagcgggtacctttgggctatttatgtccatcgggtctataatcagaagtgatagtg<br>aaagtagtccaatgtcccatcctaacctgaacctacagcaacaggcaagactggaaatgtggaagcttcgtgccaaatacg<br>gtatacgtaaggacggatcctGAGACC<br>(Uppercase, *Bsa*I sites) | This study |
| pSS122 | GAATTCctccagcattataaagcatatctaacaataccattcgggttatactgaatagcccacgcacgggttcggggttgt<br>aacgattatatgcatttcataacatgccactgtcagtggcgccacggaaggatagcgggactttatcaaatcattaatttctgtg | This study |

| | | |
|---|---|---|
| ($P_{TIM17}$-Tim17-Spot) | cataaagtgtatagtatattcatatagatatgtatatatatcattgaatacagaaatcttcgggtgtatacaacgaacattcatcgg<br>acgcctttttttactgctattgttcaacaaagtgggtaacccggcattcttgcgccaaagtttccatttttatgccaaatccagaggat<br>aaaagcactattctcatcaaaagatggaaaagctgtgaaagagtcaagttcagaagaaagcgttaagaaaaatagagta<br>cacgggagcgtt**atg**tcagccgatcattcgagagatccatgtcctatagtcatactaaatgatttcggtggtgcttttgccatggg<br>tgccattggtggtgttgtttggcatgggattaaaggttttagaaattcgccattaggtgagcgtggttcaggagctatgagcgcca<br>ttaaagcgcgtgctcccgtactgggtggtaattttggtgtgtggggtggtttattttcgacttttgattgcgctgtgaaggccgttaga<br>aagagagaggacccatggaatgctatcattgcagggtttttcactggtggcgctttagctgtaagaggtggttggaggcatac<br>aaggaacagttcgatcacgtgtgcttgtttgttgggtgtgattgaaggtgtgggactaatgtttcaaagatatgctgcttggcaag<br>ccaaacctatggctcctcctttgcccgaagcaccttcctctcaacctctgcaagctactagtcctgatagagttagagcagtctc<br>acattggtcttctggtggaggttctggcggtggttcaactccagatagagtacgtgctgtttctcactggagttct**tag**gcatgtag<br>acattatatgagccatttttcatcgttgttggaagtacccttattcggcatgtttttgttacataaatgacgtatcgcactagcctcct<br>ttacgtttttactttatttcagccttttatttcaagattaccaaccatttctctcaaccatgtacatattatatattgaaaaagtaccatac<br>ttcatctctgaagagaaaatatcaacaagccaaaagctagtgcagaaactGGATCCgcaCTGCAG<br>(Uppercase, *Eco*RI, *Bam*HI, *Pst*I sites; bold/underlined, start and stop codons) | |
| pSS140<br>($P_{TIM17}$-Tim17) | GAATTCctccagcattataaagcatatctaacaataccattcgggttatactgaatagcccacgcacgggttcggggttgt<br>aacgattatatgcatttcataacatgccactgtcagtggcgccacggaaggatagcgggactttatcaaatcattaatttctgtg<br>cataaagtgtatagtatattcatatagatatgtatatatatcattgaatacagaaatcttcgggtgtatacaacgaacattcatcgg<br>acgcctttttttactgctattgttcaacaaagtgggtaacccggcattcttgcgccaaagtttccatttttatgccaaatccagaggat<br>aaaagcactattctcatcaaaagatggaaaagctgtgaaagagtcaagttcagaagaaagcgttaagaaaaatagagta<br>cacgggagcgtt**atg**tcagccgatcattcgagagatccatgtcctatagtcatactaaatgatttcggtggtgcttttgccatggg<br>tgccattggtggtgttgtttggcatgggattaaaggttttagaaattcgccattaggtgagcgtggttcaggagctatgagcgcca<br>ttaaagcgcgtgctcccgtactgggtggtaattttggtgtgtggggtggtttattttcgacttttgattgcgctgtgaaggccgttaga<br>aagagagaggacccatggaatgctatcattgcagggtttttcactggtggcgctttagctgtaagaggtggttggaggcatac<br>aaggaacagttcgatcacgtgtgcttgtttgttgggtgtgattgaaggtgtgggactaatgtttcaaagatatgctgcttggcaag<br>ccaaacctatggctcctcctttgcccgaagcaccttcctctcaacctctgcaagct**tag**gcatgtagacattatatgagccatttttt<br>tcatcgttgttggaagtacccttattcggcatgtttttgttacataaatgacgtatcgcactagcctcctttacgtttttactttatttcag<br>ccttttatttcaagattaccaaccatttctctcaaccatgtacatattatatattgaaaaagtaccatacttcatctctgaagagaaa<br>atatcaacaagccaaaagctagtgcagaaactGGATCCgcaCTGCAG<br>(Uppercase, *Eco*RI, *Bam*HI, *Pst*I sites; bold/underlined, start and stop codons) | This study |
| pSS070<br>(pETDuet-1<br>6xHis-3C-<br>Tim44CTD) | GGATCCgctggaagtgctgtttcagggcccgacaaatatcgagtctaaagaatcgtttggtaagaaagtggaggatttca<br>aggagaaaaccgtggttggccgttctatacaatctttaaagaacaaattgtgggatgaaagtgaaaacccccttaattgttgtca<br>tgaggaaaataaccaacaaagtgggcggtttctttgcagaaacagaatcctcccgtgtttacagtcaatttaagctaatggac<br>ccaacctttttcgaacgaaagcttcaccagacacttaagagaatacattgttcccgagattctcgaagcgtatgtgaaaggcg<br>atgtcaaagttctcaaaaaatggttcagcgaggcgccattcaatgtttacgccgcccaacagaaaatcttcaaagaacagg<br>atgtttacgccgatggccgtatcctagatatcaggggcgttgaaatagtgagtgcaaagttattggctccccaagacatccca<br>gtcttggtggtcgggtgtagagcacaagaaatcaacctttacaggaaaaagaaaactggcgagattgcggctggtgacga<br>agctaatatcttgatgagctcttatgccatggttttcaccagagatccagagcaaatcgacgatgacgaaacagaagggtgg<br>aagatcttggagtttgtgcgcggggggttctagacaattcacc**taa**CTCGAG<br>(Uppercase, *Bam*HI and *Xho*I sites; bold/underlined, stop codon) | This study |
| pYC17a<br>(pYTK-e115<br>*TIM17*-HA) | GAATTCctccagcattataaagcatatctaacaataccattcgggttatactgaatagcccacgcacgggttcggggttgt<br>aacgattatatgcatttcataacatgccactgtcagtggcgccacggaaggatagcgggactttatcaaatcattaatttctgtg<br>cataaagtgtatagtatattcatatagatatgtatatatatcattgaatacagaaatcttcgggtgtatacaacgaacattcatcgg<br>acgcctttttttactgctattgttcaacaaagtgggtaacccggcattcttgcgccaaagtttccatttttatgccaaatccagaggat<br>aaaagcactattctcatcaaaagatggaaaagctgtgaaagagtcaagttcagaagaaagcgttaagaaaaatagagta<br>cacgggagcgtt**atg**tcagccgatcattcgagagatccatgtcctatagtcatactaaatgatttcggtggtgcttttgccatggg<br>tgccattggtggtgttgtttggcatgggattaaaggttttagaaattcgccattaggtgagcgtggttcaggagctatgagcgcca<br>ttaaagcgcgtgctcccgtactgggtggtaattttggtgtgtggggtggtttattttcgacttttgattgcgctgtgaaggccgttaga<br>aagagagaggacccatggaatgctatcattgcagggtttttcactggtggcgctttagctgtaagaggtggttggaggcatac<br>aaggaacagttcgatcacgtgtgcttgtttgttgggtgtgattgaaggtgtgggactaatgtttcaaagatatgctgcttggcaag<br>ccaaacctatggctcctcctttgcccgaagcaccttcctctcaacctctgcaagctactagttatccctatgacgtcccggactat<br>gcaggatcctatccatatgacgttccagattacgct**tag**gcatgtagacattatatgagccatttttttcatcgttgttggaagtacc<br>cttattcggcatgtttttgttacataaatgacgtatcgcactagcctcctttacgtttttactttatttcagccttttatttcaagattacca<br>accatttctctcaaccatgtacatattatatattgaaaaagtaccatacttcatctctgaagagaaaatatcaacaagccaaaa<br>gctagtgcagaaactGGATCCgcaCTGCAG<br>(Uppercase, *Eco*RI, *Bam*HI and *Pst*I sites; bold/underlined, start and stop codons of Tim17-HA) | |
| pYC17b<br>(pYTK-e122 $P_{DDI2}$-TIM17-HA) | GAATTCcagcccacatactactttttccttttgtttttttttttatatttcaaaggttaaactcgcttagactatgtctataatataaa<br>aaaaaaatagctctattccgtttcttttattctatttgatatttccatcacactttcatcttaatcacggatgtatactgataatagggttg<br>actgcgcctgtacggattacagtgccctcttcaattggaaaatccaagctttcaagatgggtaactgttattcaaaggatcctct<br>aagataaaacacagatcgacagatccgagagttggcttctgtgccttgggctcaaattcctttcccacctcatgcaaattgatttt | |

| | |
|---|---|
| | tctgactccaaaaaaagacagagccctgcgatagttcccgaatgttgtaacatcaaagccaagcactcctttatagaagtcg<br>catgaacgttgaatactagcagctggtgaaactacagggtctaaactaactagtatccatatctttttgagagcattgaaagtat<br>acggagtacaagctgggttagaaggaatttttatcttaacagcaatgaaaatcaactttctagactgaatccctcaagaaaatt<br>gcaaaagactaactgatactggtttaaaagagaaagatgtcaaatatgcggagttataccatcaaacaactttggacggcc<br>ccgaaacaaatgtccgcaaaaaagatcttattaaagtgcatggacactatcatttctataatacaaaatactccaccgcaca<br>atagtttgtcgggaagtcatcaatcaatcttgtacgagctttacaaataactttttaggatcggtcccctcataaaattatatataa<br>atgggttagtttccttcttcttctgttaacatgaagttgcttcgtactgtttttttgccttgcttctcttcaaaagaatcaatcggtacc**atg**t<br>cagccgatcattcgagagatccatgtcctatagtcatactaaatgatttcggtggtgcttttgccatgggtgccattggtggtgttgt<br>ttggcatgggattaaaggttttagaaattcgccattaggtgagcgtggttcaggagctatgagcgccattaaagcgcgtgctcc<br>cgtactgggtggtaattttggtgtgtggggtggtttattttcgacttttgattgcgctgtgaaggccgttagaaagagagaggaccc<br>atggaatgctatcattgcagggtttttcactggtggcgctttagctgtaagaggtggttggaggcatacaaggaacagttcgatc<br>acgtgtgcttgtttgttgggtgtgattgaaggtgtgggactaatgtttcaaagatatgctgcttggcaagccaaacctatggctcct<br>cctttgcccgaagcaccttcctctcaacctctgcaagctactagttatccctatgacgtcccggactatgcaggatcctatccata<br>tgacgttccagattacgct**tag**gcatgtagacattatatgagccatttttttcatcgttgttggaagtacccttattcggcatgttttttgt<br>tacataaatgacgtatcgcactagcctcctttacgtttttactttatttcagccttttatttcaagattaccaaccatttctctcaaccat<br>gtacatattatatattgaaaaagtaccatacttcatctctgaagagaaaatatcaacaagccaaaagctagtgcagaaactG<br>GATCCgcaCTGCAG<br>(Uppercase, *Eco*RI, *Bam*HI and *Pst*I sites; bold/underlined: start and stop codons of Tim17-HA) |
| pYC23<br>(pYTK-e115<br>*TIM23*-HA) | GAATTCattgaaaaaaagagaaaatactgaaaaaaaaaagacaccgacaaaaaaaagagagaaaggaaccttta<br>cgtagagcaaaagggaaacatttgttgcgtttaattaatagaaataataggttattgcattcgccctccattgcagaaaaaaaa<br>aaaaaaaagaccattttcctcttctactctctttgcctgtacatactacacgttatagcgttaacaaaagcagatagaaaaaaa<br>aaaaataaccaagataatataggtatacttgtttaacagatcacacacaatc**atg**tcgtggctttttggagataagacacctac<br>cgatgatgcgaatgctgccgtgggcggccaagatacaaccaagcctaaggaactatcgttgaagcagagtttaggtttcga<br>gccaaacatcaataacataatatcaggtcctggtggaatgcatgtcgacaccgctaggctgcatcctttggctggtctagaca<br>agggtgtggagtatttagatctggaagaagaacaactatcctcgttagaaggctcacagggtctgatcccttcccgtgggtgg<br>accgatgacctatgttacggtaccggtgccgtctacctgctgggacttggtatcggagggttttctggtatgatgcagggtctgc<br>agaatattccgcccaatagtcccggaaaattgcaattgaacaccgtcctgaatcacattactaagagaggtcccttcttaggta<br>ataatgcggggattctcgcgttgagctacaatatcatcaattctacaatagatgcactaagaggcaaacatgacaccgcggg<br>ctccattggcgctggggccctcacgggcgctttgttcaagtcttcaaaaggtttgaaacccatgggttattcctcggcaatggtg<br>gccgctgcgtgcgccgtctggtgtagtgtcaagaaaagactacttgaaaaaactagttatccctatgacgtcccggactatgc<br>aggatcctatccatatgacgttccagattacgct**tag**gcaacacaagaacctactctctctctctttctctctctctccctcttc<br>gcttttcccccatgcactgatggcgcttgtatatagcatttgaaaaataatagtacgtaacgcagaaaacaaaccaatgaaag<br>tagaaacccggagaaaagatctaaaaaaacaaaaaaaaaagatggaaagagcgcatgtatgtgtagatatgtacatata<br>tacaactacttgtatgttatttgccttttgtactgttaagctattgttttcgatggcGGATCCgcaCTGCAG<br>(Uppercase, *Eco*RI, *Bam*HI and *Pst*I sites; bold/underlined: start and stop codons of Tim23-HA) |
| pYC002<br>(pET32a-Cyb2Δ-DHFR) | CAT**ATG**ctgaaatacaaacctttactgaaaatctcgaagaactctgaggctgctatcctgcgcgcgtctaagactcgcttg<br>aacacaatccgcgcgtacggttctaccgttccaaaatccaagtcgttcgaacaagactcaagttccgtggcgtatctgaactg<br>gcataatggccaaatcgacaacgagccgaaactggatatgaataaacaaaagatttcgcccgctgaagttgccaagcata<br>acaagcccgatgattcgtgggttgtgatcaatggttacgtatacgacttaacgcgtttcctgccaaatcatccaggtgggcagg<br>atgttatcaagtttaacgccgggaaagatgtcactgctatttttgaaccactgcacgctcctaatgtcatcgataagtatattgctc<br>ccgaaaaaaaattgggtcccctgcaaggatccggcggtaccgttcgtccattgaactgcatcgtcgccgtgtcccaaaatat<br>ggggattggcaagaacggtgaccttccctggcctccgctccgcaacgagttcaagtacttccaacgtatgaccacaacctctt<br>cagtggaaggtaaacagaatctggtgattatgggtcgcaaaacctggttctccattcctgagaagaatcgtcctttaaaggac<br>cgtattaatattgttctcagtcgtgaactcaaagaaccaccacgtggagctcattttcttgccaaaagtttggatgatgccttacgt<br>cttattgaacaaccggaattggcaagtaaagtagacatggtttggattgtcggaggcagttctgtttaccaggaagccatgaat<br>caaccaggccacctccgcctctttgtgacacgcatcatgcaggaatttgaaagtgacacgtttttcccagaaattgatttgggg<br>aaatataaacttctcccagaatacccaggcgtcctctctgaggtccaggaggaaaaaggcatcaagtataagtttgaagtct<br>acgagaagaaagacCTCGAGcaccaccaccaccaccac**tga**<br>(Uppercase, *Nde*I and *Xho*I sites; bold/underlined: start and stop codons) |
